# Hybrid Drosophila genomes rapidly purge minority ancestry but repeatedly retain selected regions

**DOI:** 10.64898/2026.10.15.759374

**Authors:** Xinjun Zhang, Swapan Mallick, Nadin Rohland, Nick Patterson, David C. Page, David Reich, Jacob L. Mueller

## Abstract

Hybridization is increasingly recognized as a widespread evolutionary process across animals, plants, and fungi, and plays a key role in driving speciation, shaping genetic diversity, and facilitating adaptation. Initial hybrid genomes are rarely observed empirically, making it challenging to observe in real time how competing forces like purifying selection and adaptive introgression interact to shape post-hybridization genomes. Here, we use replicated experimental hybrid populations between *Drosophila simulans* and *D. sechellia* to track directly genome-wide evolutionary dynamics across approximately 45 generations post-hybridization. We show that hybrid populations with primarily *D. simulans* ancestry rapidly recover wild-type levels of fertility and fecundity, whereas populations carrying predominantly *D. sechellia* ancestry experience persistent fitness reductions and frequent extinction. Genome-wide ancestry trajectories show pervasive purifying selection against minority-ancestry alleles, leading to rapid regression toward majority ancestry across the genome, with stronger purging on the X chromosome than autosomes. Following the initial purging, neutral minority ancestry stabilized at low frequencies across the genome. In striking contrast, we observe repeated increases in *D. sechellia* ancestry on chromosome 4 and multiple regions on chromosome 3R within *D. simulans*-majority populations, providing evidence for adaptive introgression. High-density sequencing revealed that these regions were repeatedly retained across independently evolving populations despite near-complete genome-wide purging. Simulations support a model in which recombination rapidly uncouples beneficial introgressed alleles from linked incompatibility-associated alleles during the initial generations, allowing positive selection to act. Our temporal allele-frequency analyses suggest that hybrid genome stabilization of the introgressed regions is highly repeatable but not identical, yielding parallel outcomes across replicate populations. We establish a tractable experimental framework to directly observe how hybrid genomes reorganize in real time, and demonstrate how purifying selection, positive selection, recombination, and stochastic processes interact to restructure hybrid genomes shortly after admixture.

## Introduction

Hybridization and introgression are pervasive and recurrent features of speciation across animals, plants, and fungi^1–6^. Most current understandings of hybridization and introgression are inferred from retrospective studies of present-day genomes, which reflect evolutionary outcomes long after the initial hybridization^7–13^. Theoretical and empirical work suggests hybrid genomes initially undergo rapid reconstruction^11,14,15^. This reconstruction is driven by both purging deleterious introgressed ancestry due to hybrid incompatibility and recombination-mediated decoupling of beneficial introgressed alleles from linked deleterious alleles. However, retrospective and theoretical studies of hybridization cannot directly observe how purging, recombination, and adaptive introgression interact shortly after hybridization, and whether independent hybrid populations converge on similar ancestry outcomes.

Prospective studies of hybrid populations allow tracking of genome evolution through direct measurements of ancestry dynamics immediately after hybridization. Previous prospective studies of laboratory-generated *Drosophila* hybrids revealed disproportionate barriers to introgression on sex chromosomes relative to X chromosomes^16,17^, but endpoint genomic sampling, small numbers of replicate populations, or the unusual biology of neo-sex chromosomes limited these studies. Studies of natural populations of swordtail fish (*Xiphophorus*) and experimentally generated sunflower hybrids further demonstrated genome-wide purging of minority ancestry, recombination-associated retention of introgressed ancestry, and repeatable genomic outcomes^11,18–20^, however these studies generally lacked the combination of complementary majority-ancestry backgrounds, sufficient sampling from early post-hybridization generations, and large numbers of independently evolving replicate populations that were critical to directly resolve the evolutionary forces shaping hybrid genome stabilization, which we define here as the process by which hybrid genomes are reorganized following hybridization through the combined effects of selection, recombination, and drift.

We performed a prospective study of hybridization between *Drosophila simulans* and *D. sechellia,* a particularly powerful model system for studying early hybrid genome evolution. These two species offer advantages over previous prospective hybridization studies: we can backcross the same F1 hybrid separately to either parental species, track many independently evolving replicate hybrid populations, quantify X chromosome versus autosome selection, and track both increases and declines in genome-wide ancestry through time. This design allows us to distinguish negative selection against minority ancestry from positive selection consistently favoring one parental species. These two species produce fertile F1 females but sterile F1 males, in which hybrid populations maintain strong postzygotic reproductive isolation while preserving the capacity for recombination in females. *D. sechellia* has a distinct evolutionary history: it is an island population ecologically specialized to a toxic fruit, has an excess of deleterious alleles^10^, and an effective population size an order of magnitude less than *D. simulans*^21,22^. Additionally, extensive genomic resources, short generation time, and well-characterized hybrid incompatibilities within the *D. simulans* clade provide an ideal framework for studying the genomic consequences of hybridization.

Here, we established replicated hybrid populations via a single F1 female backcross to each parental species, resulting in populations with ∼75% ancestry from either *D. simulans* or *D. sechellia*, and tracked genome-wide ancestry dynamics over approximately 45 generations after hybridization. We show that hybrid genomes rapidly regress toward majority ancestry through pervasive purifying selection against minority ancestry, with the strongest purging on the X chromosome. Despite the widespread purging, specific *D. sechellia* regions that initially decline with the rest of the genome subsequently reverse trajectory and repeatedly increase in frequency within predominantly *D. simulans* populations, suggesting adaptive introgression. High-density sequencing further revealed that these regions are repeatedly retained across independently evolving populations. Simulations support a model in which recombination uncouples beneficial introgressed alleles from linked incompatibility-associated ancestry, allowing positive selection to act on introgressed variation. These findings provide a direct experimental view of how hybrid genomes stabilize, revealing (1) systematic purging of minority ancestry in both genomic backgrounds, (2) quantitatively stronger purging on the X chromosome that is two- to three-fold stronger than the autosomes, (3) specific autosomal loci that reversed the genome-wide purging trajectory and increased frequency under positive selection, and (4) highly repeatable but not deterministic genomic outcomes across independently evolving populations.

## Results

### A single backcross produces replicated hybrid populations and reveals asymmetric fitness recovery

To study the onset of post-hybridization genome dynamics, we established replicate hybrid populations between *D. simulans* and *D. sechellia* by backcrossing fertile F1 hybrid females to either parental species (Fig. 1A). This design generated 46 populations with 75% *D. simulans* ancestry and 45 populations with 75% *D. sechellia* ancestry, each propagated as a freely breeding population via non-overlapping generations. F1 hybrid males were confirmed to be infertile regardless of cross type (Supplementary Table 1), establishing an immediate selective pressure against hybrid males in both population backgrounds. Because F1 hybrid females are fertile, the hybrid populations retain the capacity for recombination and continued segregation of introgressed alleles.

**Figure 1:**
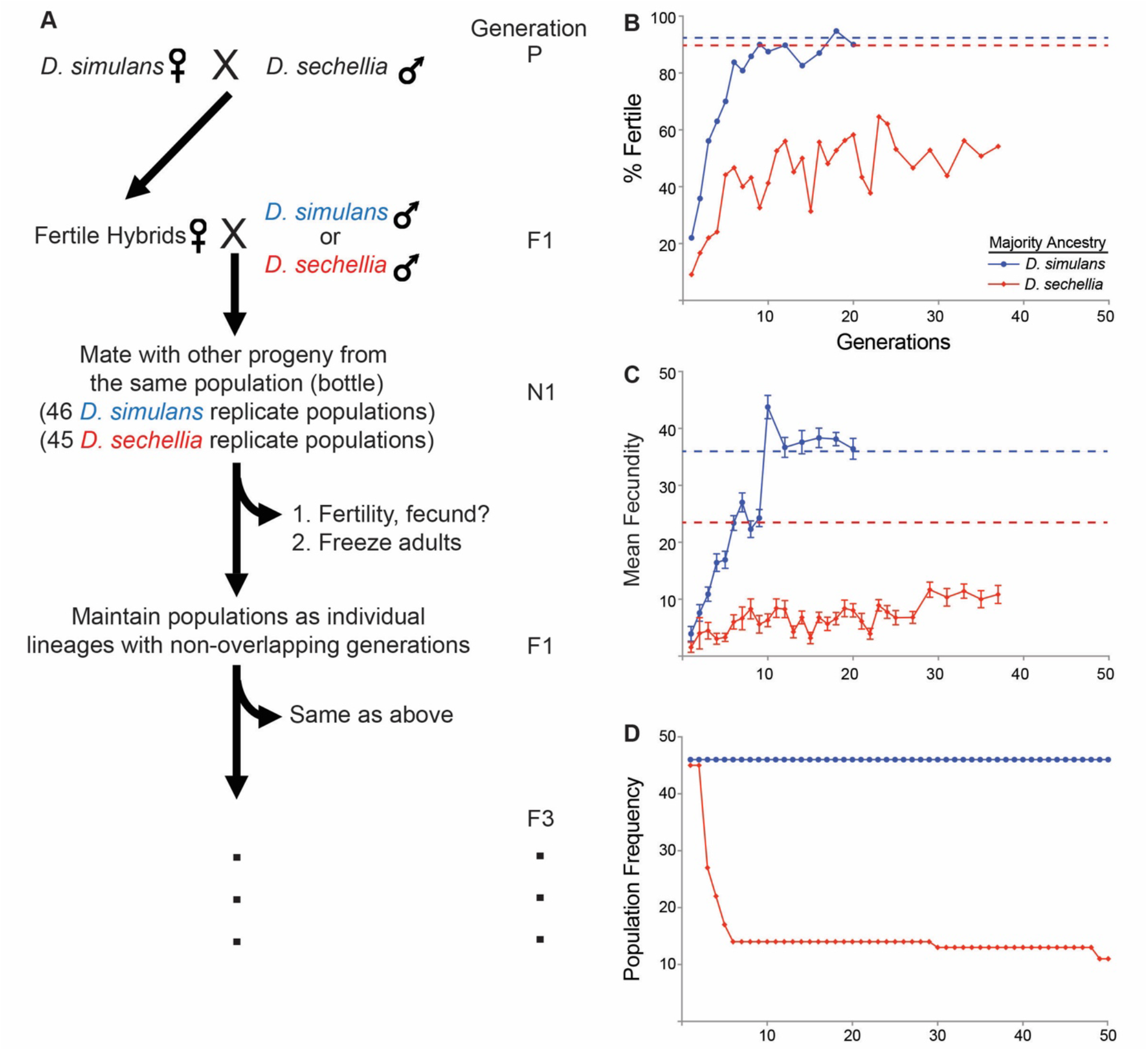
Hybrid populations can rapidly recover wild-type fertility and fecundity levels to avoid extinction. (A) Experimental design of hybrid population formation. Male and female symbols indicate multiple flies per cross. (B) The percentage of infertile single-pair matings in the *D. simulans* (red) and *D. sechellia* (blue) majority-ancestry hybrid populations that sired offspring. The dotted lines indicate the percent of infertile single-pair matings in each parental line. (C) The mean level of fecundity from each single-pair mating for each hybrid population. The dotted lines indicate the percent of infertile single-pair matings in each parental line. Error bars represent standard errors from the mean. (D) The number of populations that survived over 50 generations.

We first observe that both hybrid populations experience a substantial fitness reduction immediately following hybridization. For long-term population survival, the ability to rapidly restore fertility and fecundity is critical. We observe markedly different efficiencies in recovering the two hybrid genetic backgrounds. By randomly sampling mating pairs from each hybrid population during the initial 37 generations, we find that hybrid populations with primarily *D. sechellia* ancestry cannot regain wild-type fertility and fecundity levels (Fig. 1B, 1C). Consistent with low rates of fertility and fecundity in *D. sechellia*^23^, 31 *D. sechellia* majority-ancestry populations went extinct over the first six generations, one at generation 29 and two at generation 48 (Fig. 1D). In contrast, hybrid populations with *D. simulans* majority ancestry restore wild-type fertility and fecundity levels within 11 generations (Fig. 1B and 1C). Consistent with restoring wild-type fertility and fecundity levels, all *D. simulans* majority-ancestry populations survived (Fig. 1D). These starkly different outcomes highlight the importance of hybrid animals rapidly restoring wild-type fertility and fecundity for the persistence of hybrid populations, and that the capacity to achieve this is highly asymmetric depending on the majority ancestry background.

### Rapid post-hybridization purging of minority ancestry alleles, with faster regression on the X chromosome

We hypothesize that the survival of hybrid populations depends on purging incompatibility loci that dominate hybrid genomes and cause male infertility or inviability. To directly track how hybrid genomes evolve following hybridization, we genotyped 118 ancestry-informative genetic markers at approximately 1 MB intervals across the genome in individual flies sampled from each population over 45 generations (Supplementary Table 2). At the chromosome level, across both hybrid backgrounds, minority ancestry declined rapidly during the first 10 generations (Fig. 2A), resulting in genome-wide regression toward majority ancestry. Regression was faster in *D. simulans* majority-ancestry populations, likely due to the more severe fitness consequences (reduced fertility and fecundity) of minority ancestry in *D. sechellia* majority-ancestry backgrounds.

**Figure 2:**
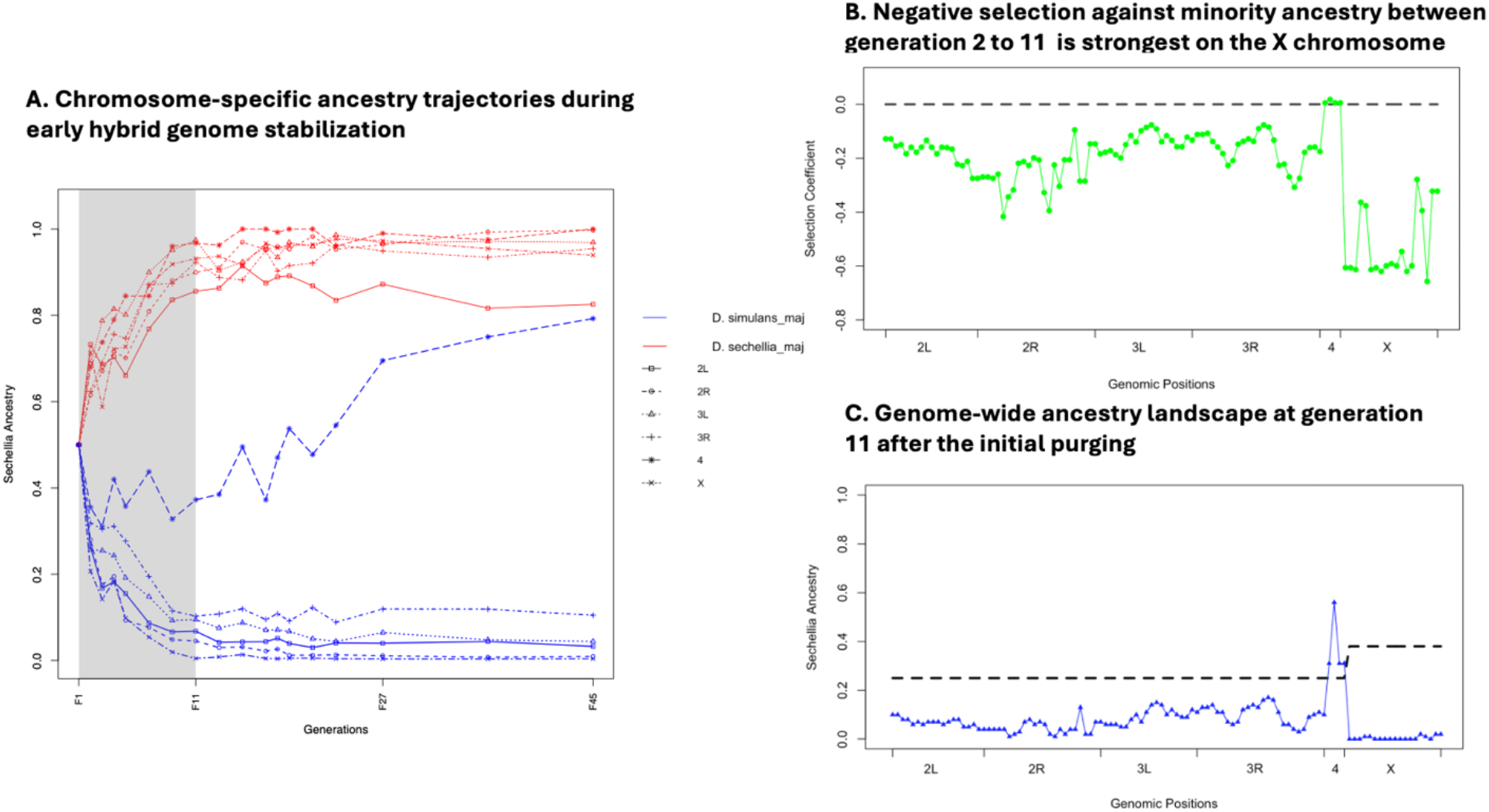
Rapid genome-wide purging reshapes hybrid genomes during the first 10 generations after hybridization. (A) Chromosome-specific ancestry trajectories in D.simulans-majority hybrid populations. Minority *D. sechellia* ancestry declines rapidly across all chromosomes during the first 10 generations after hybridization. (B) Selection coefficients estimated from allele frequency changes between generation 2 and generation 11 showed that purifying selection is strongest on the X chromosome. (C) Genome-wide ancestry landscape at generation 11 shows most genomic regions contained low *D. sechellia* ancestry after the initial phase of purging, except a small number of regions that escaped the purging and subsequently became candidates for adaptive introgression.

A striking asymmetry was observed between the X chromosome and autosomes, such that the introgressed *D. simulans* ancestry was eliminated from the X chromosome significantly faster than from the autosomes in *D. sechellia* majority-ancestry hybrids (Fig. 2A; Wilcoxon one-sided test, p-value 9.984e-06). The rapid purging of X-linked loci is consistent with the expectation that hybrid incompatibility alleles are purged more strongly from sex chromosomes than from autosomes because sex chromosomes harbor disproportionately more incompatibility alleles^24–26^, and expose them to stronger selection via hemizygosity^27–29^. To further quantify the strength of purifying selection in the first 10 generations (generations 2 through 11), we estimated selection coefficients at all genotyped loci on the autosomes and the X chromosome (Fig. 2B, Supplementary Fig. 1). Average purifying selection strength is about 2-3 fold stronger on the X chromosome than on autosomes 2 and 3 (Fig. 2B, X: s=-0.523; 2L+R: s=-0.220; 3L+R: s=-0.154; 4: s=0.008), consistent with known X-linked hybrid incompatibility loci between *D. simulans* and *D. sechellia*^23^.

Although purging eliminates most introgressed ancestry within the first 10 generations, it does not completely remove minority ancestry. Instead, genome-wide introgressed ancestry stabilizes at low frequencies across much of the genome (Fig. 2C), suggesting that a subset of introgressed alleles escapes complete elimination and persists in hybrid populations. This residual introgression likely represents a combination of neutral loci and very weakly selected alleles that were not efficiently removed by purifying selection. However, ancestry persistence is not uniform across the genome. While most loci remain at low frequencies, a small number of genomic regions subsequently diverge from the genome-wide trend and increase in frequency, indicating that hybrid genome stabilization involves both widespread ancestry loss and selective retention of specific introgressed haplotypes.

### Repeated retention of specific *D. sechellia* genomic regions during hybrid genome stabilization

In contrast to the pervasive genome-wide elimination of minority ancestry, we observe that *D. sechellia* ancestry repeatedly increases in frequency in several regions. Using our 1MB marker panel, we identify three regions that consistently escape purging in *D. simulans*-majority populations: the entire chromosome 4, and two regions on chromosome 3R near 5.9 MB and 12.9 MB (Fig. 3A). All three regions initially decline in frequency alongside the rest of the genome, likely due to the purging of linked incompatibility alleles with strong negative selection coefficients. However, starting in generation 11, *D. sechellia* allele frequencies at these loci diverge sharply from the genome-wide trend and increase, indicating positive selection for introgressed alleles that becomes effective after recombination breaks their linkage to strongly deleterious alleles (Fig. 3B).

**Figure 3:**
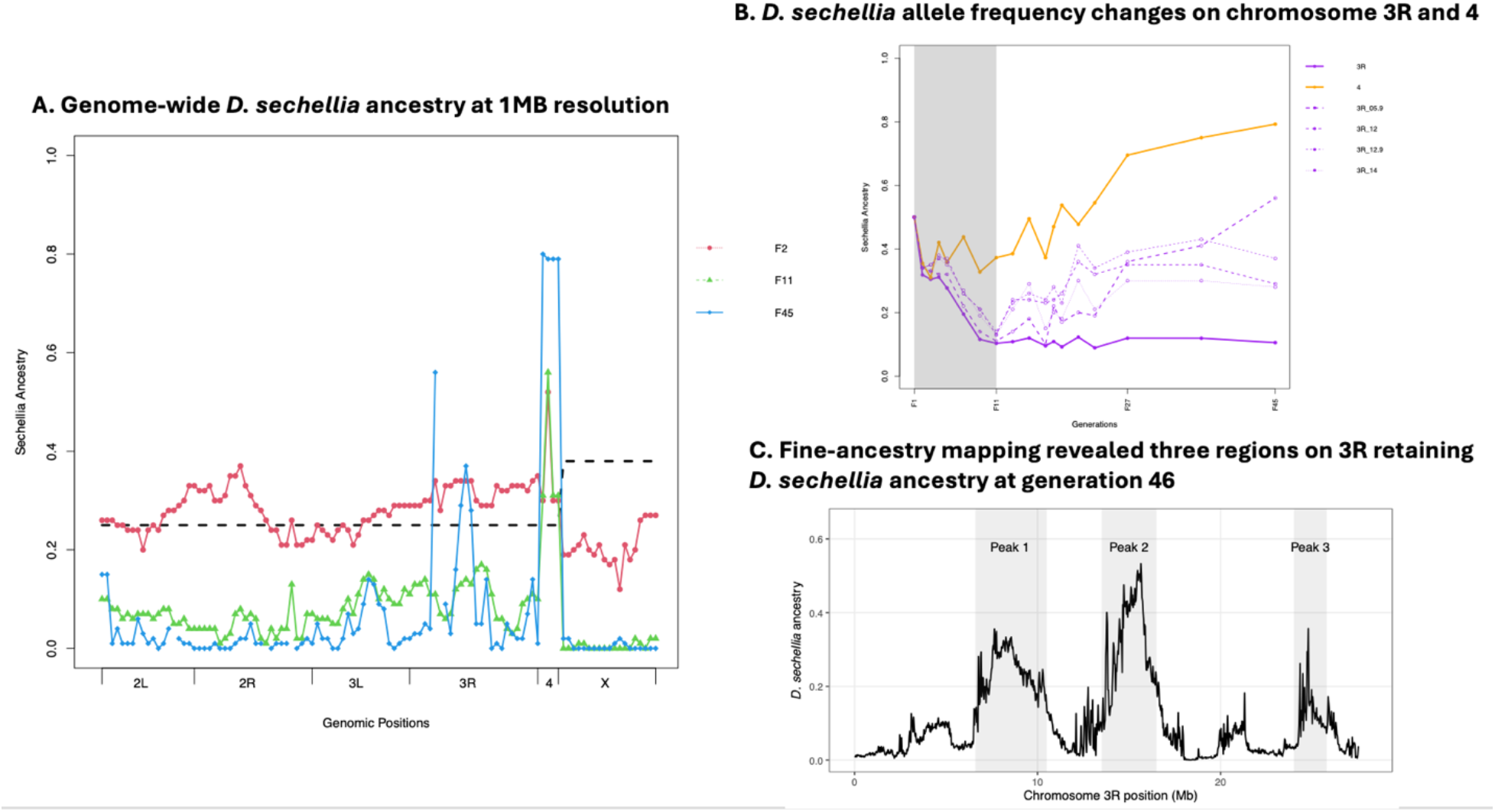
Chromosome 4 and three 3R regions are repeatedly retained in hybrid populations with *D. simulans* majority ancestry. (A) Genome-wide *D. sechellia* ancestry at generations 2, 11, and 45 with ancestry-informative markers at ∼1 MB resolution. Most of the genome retains little minority-ancestry DNA after intensive post-hybridization purging, while the chromosome 4 and 3R regions show elevated *D. sechellia* ancestry in a *D. simulans* majority genome ancestry. (B) Temporal trajectories of chromosome 4 and 3R ancestry-informative 1MB markers in *D. simulans*-majority populations. Chromosome 4 and several 3R markers increased in frequency after generation 11 while most introgressed ancestry declined rapidly following hybridization (C) Fine-ancestry mapping of 3R using 30kb windows showed elevated *D. sechellia* ancestry at 6.6-10.5MB, 13.5-16.5MB, and 24.0-25.8MB.

To further determine the mechanism underlying the trajectory at these regions, we generated a high-density Illumina sequencing dataset from all surviving *D. simulans*-majority populations at generation 46. We show that, genome-wide, *D. sechellia* ancestry declines to approximately 0.5% (Supplementary Fig. 2), confirming the intensive purging of minority ancestry. High-resolution ancestry mapping identifies the 3R introgressed regions and reveals a third region on chromosome 3R with increased frequency near 24-26 MB that is undetectable with low-density markers (Fig. 3C), indicating chromosome 4 and three 3R regions may be under strong positive selection.

We quantified the strength of positive selection driving the introgressions of chromosome 4 and 3R during generations 11 through 45. The *D. sechellia* chromosome 4 exhibits strong positive selection (mean s=0.055, Supplementary Fig. 3), consistent with the expectation that positive selection acts more efficiently on a non-recombining chromosome^30,31^. The two originally identified 3R loci exhibit positive selection of different magnitudes (3R:5.9MB s=0.069; 3R:12.9MB s=0.037), reflecting the expected reduced efficiency of positive selection in recombining chromosomes linked to deleterious, incompatibility-associated alleles. These findings suggest that while most introgressed ancestry is deleterious, a small subset of introgressed *D. sechellia* alleles confer a selective advantage in the *D. simulans* ancestry background and repeatedly escape genome-wide purging, and rise in frequency, contributing to the stabilization of hybrid genomes.

### Recombination rapidly decouples beneficial introgressed variants from linked incompatibility loci

We hypothesize that the delayed increase in *D. sechellia* alleles on chromosome 3R can be explained by the time required for recombination to uncouple beneficial introgressed alleles from the linked incompatibility-associated haplotypes into which they initially arrived. To test this, we simulate the decay of linkage disequilibrium among ancestry-informative markers on chromosome 3R in populations of size N=50, matching the effective size of our experimental bottles (Supplementary Fig. 4). We show that for markers approximately 4 MB apart (such as 3R at 9.0 MB and 12.9 MB), LD decays rapidly during the first 10 generations and approaches 0 by generation 20. For markers approximately 2MB apart (such as 3R at 2.9 MB and 5.9 MB), LD also decays substantially during the initial generations, reaching 0 by generation 100. This LD delay is consistent with the timing of the observed allele-frequency trajectories, suggesting that the initial 10 generations of decline reflect the purging of linked incompatibility haplotypes, while the subsequent increase reflects positive selection on beneficial alleles once they escaped purging via recombination.

To further evaluate the dynamics between positive selection of introgression and purifying selection of purging of incompatibility loci, we simulated the chromosome 3R markers using empirically estimated selection coefficients, with average s = 0.05 for positively selected chromosome 3R loci (5.9MB and 12.9MB) and average s = -0.2 for flanking loci (Fig. 4). Simulations reproduce the empirically observed allele frequency trajectories, whereas neutral simulations do not. Our findings support a model in which recombination rapidly reshapes hybrid genomes by separating beneficial introgressed alleles from linked deleterious loci.

**Figure 4:**
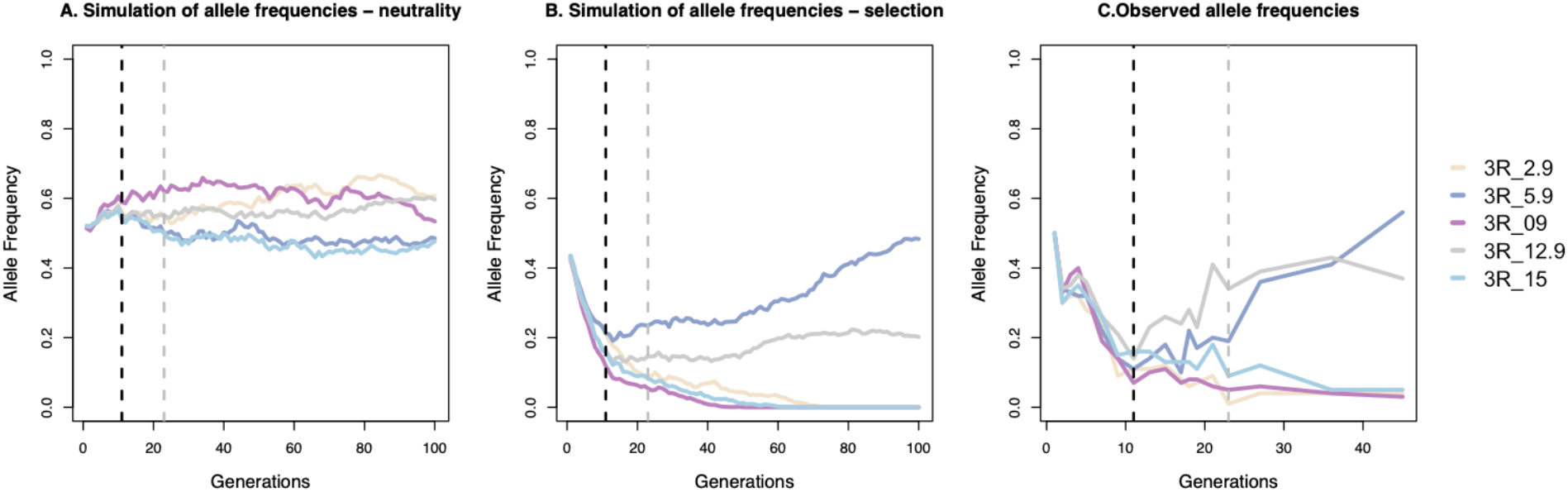
Simulations of positive selection facilitate understanding of the increase in allele frequencies on 3R. (A) Simulations of 5 SNP markers, 3R_2.9, 3R_5.9, 3R_09, 3R_12.9, and 3R_15 under neutrality. We plot the mean allele frequency at each marker for 100 generations. (B) Simulations of natural selection acting on the marker alleles, with 3R_5.9 and 3R_12.9 under positive selection and the other markers under purifying selection. (C) Observed allele frequency changes in empirical data.

### Independent positive selection repeatedly retains *D. sechellia* chromosome 4 and 3R loci in a *D. simulans* genetic background, but it is not deterministic

The increase in frequency of the chromosome 4 and 3R regions is highly repeatable across independently evolving replicate populations (Fig. 5A), however this retention is not deterministic, as independent populations show substantial variation in ancestry levels. For example, although chromosome 4 was retained in the vast majority of populations, it is nearly lost in a replication bottle (Fig. 5A). Similarly, the 3R regions show incomplete retention across populations despite evidently experiencing positive selection (Fig. 5A+B). These observations suggest that hybrid genome stabilization follows repeatable evolutionary trajectories while remaining influenced by stochastic processes such as drift.

**Figure 5:**
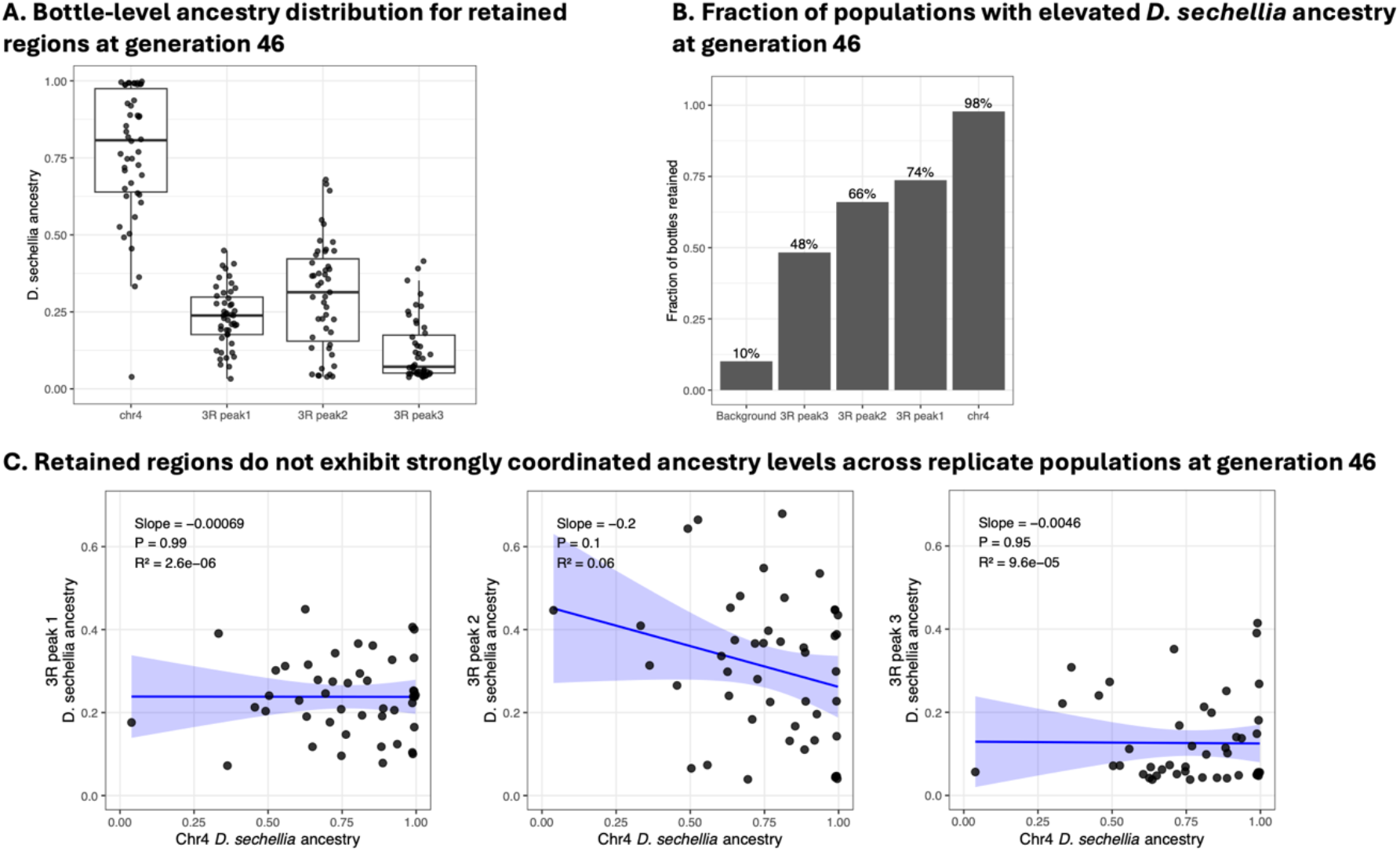
Illumina sequencing at generation 46 revealed repeated retention of introgressed regions producing parallel but non-identical ancestry outcomes across independently evolving populations. (A) Distribution of *D. sechellia* ancestry across replicate populations for chromosome 4 and the three 3R regions. Each population represents a point, and the boxplot summarizes the distribution of ancestry values. We show that although retained regions consistently exceeded the genomic background, ancestry levels vary substantially among populations across all regions. (B) Proportion of replicate populations with elevated ancestry within each retained region. Chromosome 4 was retained in nearly all populations, whereas the 3R regions showed intermediate levels of repeatability. Background genomic windows showed little evidence of elevated ancestry. These results show that hybrid genome stabilization is highly repeatable but not deterministic. (C) Relationship between chromosome 4 ancestry and the three retained 3R regions across replicate populations. Linear regression revealed little evidence of coordinated ancestry change among regions (all p-values >0.1). Although chromosome 4 and 3R regions were repeatedly retained across populations, frequency changes differed among populations and were not statistically predictive across loci.

The rise in allele frequency in the chromosome 4 and 3R regions could be explained by epistasis or independent positive selection. To distinguish between these two possibilities, we examine coordinated allele-frequency changes across replicate populations from generations 11 to 27, corresponding to the period when retained regions began to increase in frequency, and at generation 46. From generations 11 to 27, most allele pairs of loci show patterns consistent with independent positive selection, while chromosome 4 and loci within the 12-13MB region of 3R locus pairs exceed the empirical 95^th^ percentile of the genome-wide distribution (25 bottles; Supplementary Figure 5), suggestive of epistasis. However, at generation 46, contrary to expectations under a simple epistasis model, chromosome 4 ancestry did not strongly correlate with ancestry levels at the major 3R peaks (Fig. 5C). Therefore, although we identified coordinated changes in temporal allele frequencies, the endpoint generation 46 ancestry patterns across replicate populations did not provide strong support for persistent epistasis. These observations suggest that interactions among retained loci may be more complex than a simple pairwise epistatic model; however, regardless of the causal mechanism, our results indicate that hybrid genome stabilization does not depend solely on locus-specific selection and may involve coordinated evolutionary responses among introgressed regions.

## Discussion

The *D. simulans* and *D. sechellia* species establish a powerful, systematic experimental framework for studying real-time hybridization dynamics. By experimentally generating and tracking replicated hybrid populations between *D. simulans* and *D. sechellia*, we capture the real-time reconstruction of hybrid genomes. Our results reveal four features of hybrid genome stabilization: 1) rapid genome-wide purging of minority ancestry driven by hybrid incompatibility in both genomic backgrounds, 2) disproportionately strong purifying selection on the X chromosome, 3) repeated retention of a small number of introgressed regions enabled by positive selection that reverse the genome-wide trajectory, and 4) parallel but non-identical evolutionary outcomes across replicate populations (Fig. 6). These joint forces ultimately lead to a structured mosaic genomic landscape of ancestry, rapidly shaped by the interplay between purifying selection, positive selection, recombination and stochastic processes over short timescales (Fig. 6). More broadly, our results show that the forces shaping hybrid genome evolution can be strong enough to produce repeatable patterns across populations, yet insufficient to drive populations toward a single, fixed genomic outcome.

**Figure 6:**
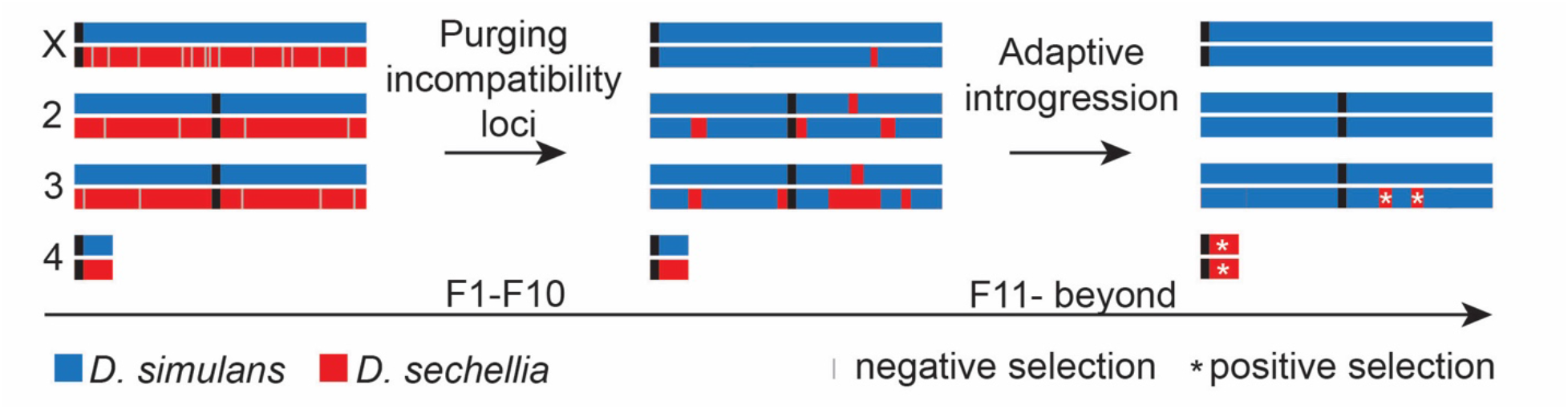
Model of early hybridization events. Strong purifying selection removes hybrid incompatibility loci during the initial 10 generations, with even stronger selection on the X chromosome, followed by introgression of selectively advantageous loci.

Our observations of early hybrid genome dynamics in real time are consistent with previous experimental evolution studies of hybrids in flies^17^ and swordtails^18^ and with evolutionary theory^29^ of how deleterious alleles are purged. We extend these previous studies in several ways. First, we show that minority ancestry is systematically purged despite substantial differences in fertility and fecundity between *D. simulans*- and *D. sechellia*-majority populations. Second, we directly quantify the large X-effect as it develops over time, showing that purifying selection is 2-3 times stronger on the X chromosome than on autosomes, plausibly explained by both hemizygous exposure of incompatibility alleles in hybrid males and the disproportionately enriched fitness-related loci on sex chromosomes. Third, we observe that minority ancestry is not eliminated but instead stabilizes at low frequencies across much of the genome, likely reflecting neutral genetic drift and weakly deleterious alleles that escape initial purging. Whether these low-frequency introgressed alleles have long-term functional effects is an open question.

Our findings also directly capture the transition from genome-wide purging to locus-specific adaptive introgression enabled by recombination on chromosome 3R. Most studies of adaptive introgression focus on empirical data long after hybridization, whereas our experimental system observed positive selection in the early stages following initial hybridization, showing that beneficial alleles can begin to spread as early as 10 generations post-hybridization. Our simulations support a model in which recombination decouples beneficial alleles from deleterious haplotypes, allowing positive selection to act despite pervasive genome-wide purifying selection. This provides a mechanistic explanation for why high-recombination regions harbor more introgressed ancestry in natural populations (Fig. 6). By contrast, the chromosome 4 dynamics likely reflect a different evolutionary process. Because chromosome 4 is non-recombining, both beneficial and deleterious mutations are expected to remain linked. The rapid increase of *D. sechellia* chromosome 4 therefore suggests that the chromosome’s net fitness effect became advantageous within the *D. simulans*-majority genomic background, despite its deleterious variants.

We propose three non-mutually exclusive mechanistic explanations for why chromosome 4 from *D. sechellia* is transmitted more frequently than from *D. simulans*. One possibility is a transmission advantage, where the *D. sechellia* chromosome 4 acquired a “selfish” transmission ratio distorter. The 4^th^ chromosome was ancestrally a sex chromosome^32^, and sex chromosomes are known to be enriched for transmission ratio distorters^33^. 2) A second possibility is a structural or chromosomal mechanism, where structural differences of chromosome 4 between *D. simulans* and *D. sechellia* lead to nondisjunction in meiosis. Chromosome 4 in *Drosophila* is known to undergo nondisjunction at a higher rate than other chromosomes^34^, which could favor greater transmission of *D. sechellia*. The *D. sechellia* chromosome 4 differs from the *D. simulans* chromosome 4 in having telocentric centromeres, newly generated by non-LTR retroelements^35^, which could lead to nondisjunction, favoring *D. sechellia* chromosome 4 transmission. 3) A third possibility is positive selection for fitness benefits of the *D. sechellia* 4^th^ chromosome. In another Drosophila lineage, the 4^th^ chromosome acquired a new gene that influences fitness^36^, a pattern that may also have occurred in *D. sechellia*. Distinguishing among these three possibilities requires future studies to assess transmission rates in males and females, examine chromosomal nondisjunction, and functionally characterize candidate *D. sechellia* 4^th^-chromosome genes.

Our results demonstrate that hybrid genome stabilization is highly repeatable but also not deterministic. Selection consistently favors particular regions while stochastic evolutionary processes continue to influence their endpoint frequencies. Considering the unique island population history of *D. sechellia*, it still provides opportunities to generate novel combinations of selectively advantageous alleles when hybridized with *D. simulans*. The rapid introgression of a few genomic regions (on 3R and chromosome 4) from *D. sechellia* into *D. simulans* suggests it could also occur in the wild. Indeed, *D. sechellia*/*D. simulans* hybrids have been detected in the Seychelles^37^, but whether the *D. sechellia* chromosome 4 and regions of 3R have introgressed into *D. simulans* in the wild is unknown. More broadly, and over longer timescales, introgression is observed throughout Drosophila evolution^2^, raising the question of how much of the genome introgresses between any two closely related species. Our laboratory-based hybridization strategy, using a single backcross and replicate populations, provides an opportunity to assess how much of an organism’s genome can fly under the radar as selectively advantageous in closely related species.

## Acknowledgments

We thank G. Dokshin, P. Muller, J. Neubauer, and L. Okumura for technical assistance; C. Aquadro, A. Clark, D. Presgraves, S. Sankararaman, and J. Sitnick for technical advice; and T. Orr-Weaver for using her fly room. David Page and David Reich are Investigators of the Howard Hughes Medical Institute, and this research was also supported by startup funds from Harvard Medical School to David Reich.

## Methods

### Fly stocks and population husbandry

*D. simulans* – South Africa (#14021-0251.169) and *D. sechellia* – Cousin Island (#14021-0248.07) stocks were obtained from the UCSD species stock center. All populations were maintained in bottles on a standard medium containing cornmeal, molasses, and yeast. In the initial cross to generate F1 hybrids and the subsequent backcross, 3-5 days old (peak of sexual maturity) males and females were collected, yielding ∼30 males and ∼20 females per bottle. To generate F1 hybrid males and females, 2061 *D. simulans* virgin females were crossed to *D. sechellia* males. F1 hybrid males were crossed to *D. simulans*, *D. sechellia*, or F1 hybrid virgin females to assay hybrid male infertility.

To initiate the hybrid populations, ∼2000 F1 hybrid females were backcrossed to either *D. simulans* or *D. sechellia* males in roughly equal proportions (∼1000 F1 females to each) with ∼30 males and ∼20 females per population (each population was propagated in bottles). This generated 46 populations with primarily *D. simulans* ancestry and 45 with primarily *D. sechellia* ancestry. F1 hybrid females and parental males were allowed to breed freely for nine days, and then all flies were frozen to allow the first generation of backcrossed hybrids to eclose. To have non-overlapping generations, three and six days after the first-generation progeny eclosed, adults were transferred to a new bottle, which would yield only second-generation progeny. Nine days after the first flies were transferred to the new bottle, all flies were frozen to allow only the subsequent generation progeny to populate the bottle. This breeding scheme was repeated across all populations, each generation for 50 generations.

### Fertility and fecundity assays

Single-pair matings were performed with hybrid males and females to measure the presence of progeny (fertility) and progeny output (fecundity). Virgin hybrid male and female mating pairs, 3-5 days of age (peak of sexual maturity), were randomly selected from the populations and placed in vials. After two days, the male was removed, and the female was allowed to lay eggs for eight days. If any mating pair produced progeny, they were considered fertile. To estimate fecundity, the total number of adult progeny produced was counted 21 days after mating. Fecundity was measured in two ways: including all single-pair matings (overall fecundity) or only those matings producing offspring. At least 30 single-pair matings were performed for *D. simulans* and *D. sechellia* majority-ancestry populations at all time points except two: *D. sechellia* majority-ancestry generations 2 and 7. For *D. simulans* majority ancestry populations, matings were assayed at generations 1-10, 12, 14, 16, 18, and 20. We stopped assaying fertility and fecundity in the *D. simulans* majority-ancestry populations at generation 20 because the hybrid populations had consistently maintained wild-type levels of fertility and fecundity over multiple generations (Fig. 1A and 1B). For *D. sechellia,* for most ancestry populations, matings were assayed at generations 1-25, 27, 29, 31, 33, 35, and 37. All single-pair matings were performed in fly vials containing standard medium with cornmeal, molasses, and yeast.

### Identification and genotyping of ancestry informative markers

We utilized two strategies to genotype flies. First, to track the ancestry of each generation, we used a 1-MB-interval map of ancestry-informative genetic markers that were fixed nucleotide differences between *D. simulans* and *D. sechellia*. Second, we used a high-density map to confirm the 1MB-interval map and fine-map specific regions at generation 46. We used females for all genotyping to obtain higher X-chromosome allele frequencies. Thus, all mitochondria were of *D. simulans* origin, and the origin of the Y chromosome depended on whether the population was initiated with a backcross to *D. simulans* or *D. sechellia* males (Fig. 1A), which we confirmed experimentally.

### 1MB-interval map

We used a targeted approach to identify species-specific ancestry-informative markers at ∼1 MB resolution across the Drosophila genome. At the onset of our experiment, the *D. simulans* and *D. sechellia* genomes had been sequenced^38^, allowing us to identify an evenly spaced collection of ancestry informative markers.

Sequenom’s MassARRAY Designer software was used to pool primers (30-40 markers per pool) for 118 ancestry informative markers (Supplementary Table 2). All primers for PCR and iPLEX reactions were ordered unmodified, with standard and purified from Integrated DNA Technologies. Forward and reverse PCR primers were ordered in 96-well deep-well plates, mixed by the vendor at final, equimolar 240 μM concentrations. PCR primers underwent further dilution to a working concentration of 1 μM each. Probes (for iPLEX extension) were ordered unmixed in 96-well deep-well plates at 250 to 450 μM.

With our collection of 118 ancestry informative markers (Supplementary Table 2), each marker was assayed on genomic DNA isolated from single female flies, frozen and stored at -80**°**C, and put in 96-well plates. We also tested each marker on genomic DNA isolated from single female flies, as previously described^39^, from the parental *D. simulans* and *D. sechellia* lines used in this experiment to confirm it accurately distinguished between the two species. We assayed a single female from each population at generations 1-4, 6, 10, 12, 14, 16, 18, and 20. We sampled each population’s genome ancestry deeper by assaying five individual females at generations 8, 17, 26, 35, and 44. Genomic DNA was prepared for genotyping on the Sequenom MASSARRAY system (Sequenom, San Diego, Calif). This assay consists of a marker-specific PCR amplification followed by a single base extension using mass-modified ddNTP terminators.

The marker-specific PCR was performed in a total volume of 6 μL with ten ng genomic DNA, a final primer concentration of 1 μmol/L, 2.5 mmol/L MgCl2, 0.2 mmol/L of each deoxynucleotide triphosphate (dNTP), and 0.1 U HotStarTaq DNA Polymerase (QIAGEN, Hilden, Germany). Hot start was performed at 94°C for 15 minutes, followed by 45 cycles of 94°C for 20 seconds, 56°C for 30 seconds, and 72°C for 1 minute, and finished at 72°C for 3 minutes. To remove excess dNTPs, shrimp alkaline phosphatase (Amersham, Freiburg, Germany) was added to the PCR products for 40 minutes at 37°C, followed by inactivation of the phosphatase at 85°C for 10 minutes.

The base-specific extension reaction was performed using Sequenom’s iPLEX GOLD reagent kit per their recommendations. For the base extension reaction, the denaturation was performed at 94°C for 2 minutes, followed by 94°C for 5 seconds, 56°C for 5 seconds, and 72°C for 5 seconds for 40 cycles.

Extension products were then identified using MALDI-TOF mass spectrometry. From this point on, the Broad Institute’s Genetic Analysis Platform performed all genotyping steps using Sequenom technology, including resin clean-up, spotting of probes on the chip, and detection. SpectroTYPER software was used to inspect and manually curate the genotype output.

### High-density map

We performed Illumina sequencing on genomic DNA of all primarily *D. simulans* populations at generation 46. Genomic DNA was isolated, as previously described, from 10 females per population, including the two parental lines. To prepare the genomic DNA for Illumina sequencing, we sonicated all DNAs using a Covaris Focused-Ultrasonicator (E210) to obtain sheared fragments averaging ∼150bp. Illumina sequencing barcoded libraries were generated using the sheared genomic DNA as previously described^40^. Finished libraries were pooled according to TaqMan® real-time PCR quantifications and test-sequenced on a single lane of Illumina GAII for 36 cycles, single-end. After re-pooling the libraries according to the test-sequencing quantification estimates, six additional lanes were sequenced for 72 cycles, single-end. We generated deeper sequencing coverage of the parental strains’ genomes to identify species-specific nucleotide differences as ancestry-informative markers.

### Mapping to ancestry-informative markers

Illumina sequences from the parental lines and individual bottles were mapped to *D. simulans* and *D. sechellia* genome assemblies, downloaded from FlyBase in 2010 release archives. The genomes corresponded to the 12 Drosophila genomes project assemblies dsim_r1.4 and dsec_r1.3^41^. We obtained ∼3-fold coverage of the parental populations and ∼0.3-fold coverage of each hybrid population bottle. We determined ancestry-informative sites by pairwise alignment of the genomes and recording the consistent differences.

### Processing of high-density Illumina sequencing data

We used processed SNP-level read count files generated from the alignments for all downstream analyses, and we make these fully available as a Supplementary Data File. Initial processing identified approximately 2.4 million candidate SNPs between *D. simulans* and *D. sechellia* parental genomes. We filter SNPs that are 1) covered by a typical number of reads across populations (14-44 total reads), and 2) are covered by at least one read in both parental populations, and 3) show complete agreement between parental reads and expected species-specific alleles. After filtering, we retained 673,183 ancestry-informative SNPs. For each SNP, *D. sechellia* ancestry was estimated as *n_sech_/(n_sim_+n_sech_),* where *n_sech_* and *n_sim_* are the numbers of *D. sechellia*-like and *D. simulans*-like reads.

### Genome-wide ancestry inference from high-density Illumina sequencing

SNPs from Illumina sequencing data were aggregated into non-overlapping 300kb windows to determine ancestry across the genome. For each window, *D. sechellia* ancestry was estimated as the depth-weighed mean ancestry across all SNPs within the window. Windows with insufficient coverage were excluded from the analysis. We used a 300 kb window size because it provides robust ancestry estimates and minimizes stochastic variation among individual SNPs, and this window size is comparable to the expected extent of LD block sizes after 45 generations of evolution post-hybridization. To refine candidate regions identified by the 300kb analysis, we repeated the ancestry estimation using non-overlapping 30kb windows to localize ancestry peaks.

### Bottle-level repeatability analysis

We estimate ancestry for each *D. simulans*-majority populations at generation 46 to determine whether elevated *D. sechellia* ancestry is a repeatable pattern or from a small number of outlier populations. For each bottle and genomic window, we quantify the proportion of bottles exhibiting elevated ancestry within each genomic region. Regions repeatedly retained across independent populations are interpreted as evidence for repeatable evolutionary dynamics.

### Temporal allele frequency trajectory analysis

We examine allele frequency change across generations using the 118-marker ancestry panel. For each marker, *D. sechellia* allele frequencies were estimated at each generation, and trajectories were compared by direction and magnitude of allele frequency change over time. This analysis was used to identify loci exhibiting coordinated evolutionary dynamics, and because fitness was not measured directly from these loci, this analysis was not intended for a formal test of epistasis.

### Selection coefficient estimation

We estimate locus-specific selection coefficients from observed allele frequency changes through time. Let *p*_0_ and *p_n_* denote the frequency of *D. sechellia* ancestry at the initial generation and after n generations, respectively. Because the experimental populations were predominantly *D. simulans*, we treated *D. sechellia* ancestry as the focal minority ancestry and assigned relative fitness as *w_sec_*_ℎ_ = 1 and *w_sim_* = 1 − *s*, where the positive values of selection strength s indicate that *D. sechellia* ancestry has higher relative fitness and negative values indicate selection against *D. sechellia* ancestry.

Under this parameterization, at generation 0, the expected contributions of the two ancestry components are proportional to *p*_0_ and 1 − *p*_0_. Assuming their relative fitness difference remains constant across generations, after n generations these contributions are proportional to *p*_0_ and 1 − *p*_0_/(1 − *s*)*^n^* respectively. Normalizing by their sum gives the expected *D. sechellia* ancestry frequency:

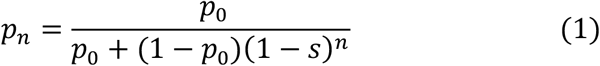

Equivalently, the probability of *D. simulans* relative to *D. sechellia* ancestry change is by a factor of (1-s) per generation, which is, 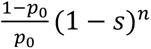. And solving for s gives:

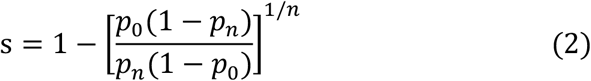

### Simulation of LD decay and allele frequency changes

We used SLiM to simulate the evolution of a 50-individual population across a 13MB genomic segment, matching the size of the 3R genomic region and the effective population size of the bottles. At the 500kb (3R_2.9), 3.5MB (3R_5.9), 6.5MB (3R_09), 10.5MB (3R_12.9), and 12.5MB (3R_15) positions, we inserted neutral marker alleles at 50% frequency, representing the introgressed *D. sechellia* ancestry. The simulated segment has a uniform mutation rate (1.5e-9) and recombination rate (2.3e-8), matching the empirical estimates in *D. simulans*^42,43^. At every generation, we computed the linkage disequilibrium measurement r^2^ using marker allele and haplotype frequencies and plotted the decay of linkage disequilibrium (LD) over 100 generations. To observe allele frequency and LD changes under selection, we repeated the simulations using estimated selection coefficients from the observed marker data. We show that under neutrality and selection, the LD between markers decayed substantially by the 11^th^ generation, coinciding with the time that allele frequency trajectories of 3R_5.9 and 3R_12.9 deviated from the rest of the autosomes. By the 23^rd^ generation, most introgressed alleles linked on the same haplotype have decoupled, allowing positive selection to act on loci at and nearby 3R_5.9 and 3R_12.9. We show that the observed allele frequencies and simulated results under selection matched well, and neither were observed under neutral simulations, suggesting that the increase of allele frequencies on 3R was driven by positive selection rather than stochastic processes. The simulations were designed to investigate the two 3R loci identified from a 1MB marker panel on temporal changes in ancestry. The third candidate region was identified only from the Illumina data on generation 46 and therefore could not be incorporated into the temporal simulations.

## Data availability

The high-density Illumina sequencing data generated in this study have been deposited in the European Nucleotide Archive (ENA) under accession PRJEB127256. Processed SNP-level read counts used for downstream ancestry analyses are provided as a Supplementary Data file. Ancestry informative markers and sequencing barcodes are provided in Supplementary Tables 2 and 3. All other data are available within the manuscript and Supplementary Information.

## Code availability

Custom scripts used for ancestry analyses, estimation of selection coefficients, population-level repeatability analyses, statistical analyses, and forward-time simulations in SLiM are available upon request.

