## SupplementaryFiguresandTables for "Hybrid Drosophila genomes rapidly purge minority ancestry but repeatedly retain selected regions"

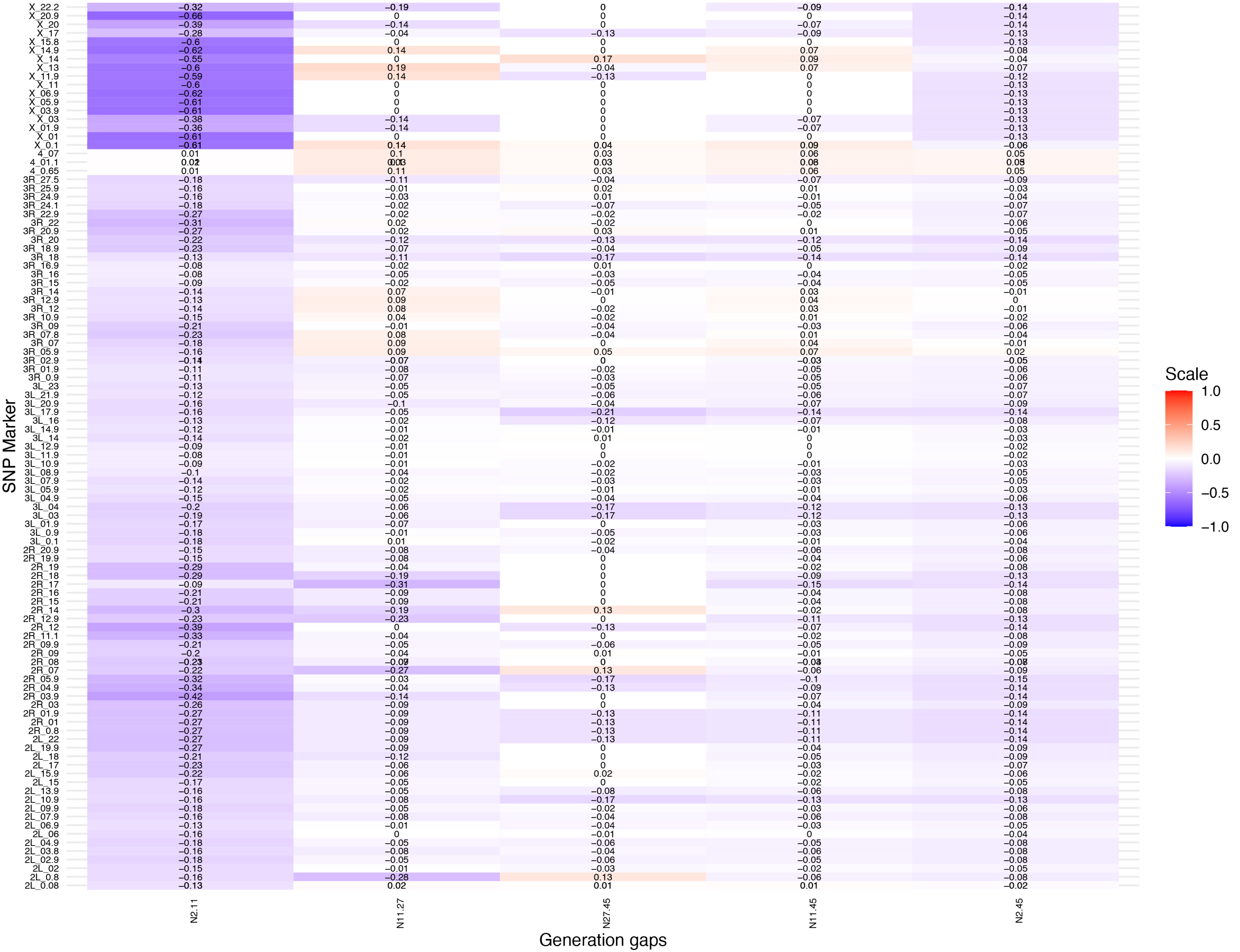


**Supplementary Figure 1. Genome-wide selection coefficient estimates across different generation intervals.** Heatmap showing estimated selection coefficients for every ancestry-informative SNP marker across the four generation intervals (N2-11, N11-27, N27-45, N11-45). Rows correspond to SNP markers ordered by genomic position, and columns represent generation intervals. Negative values indicate decreases in *D. sechellia* ancestry, whereas positive values indicate increases, together with persistent positive selection at chromosome 4 and several chromosome 3R regions across later generations.

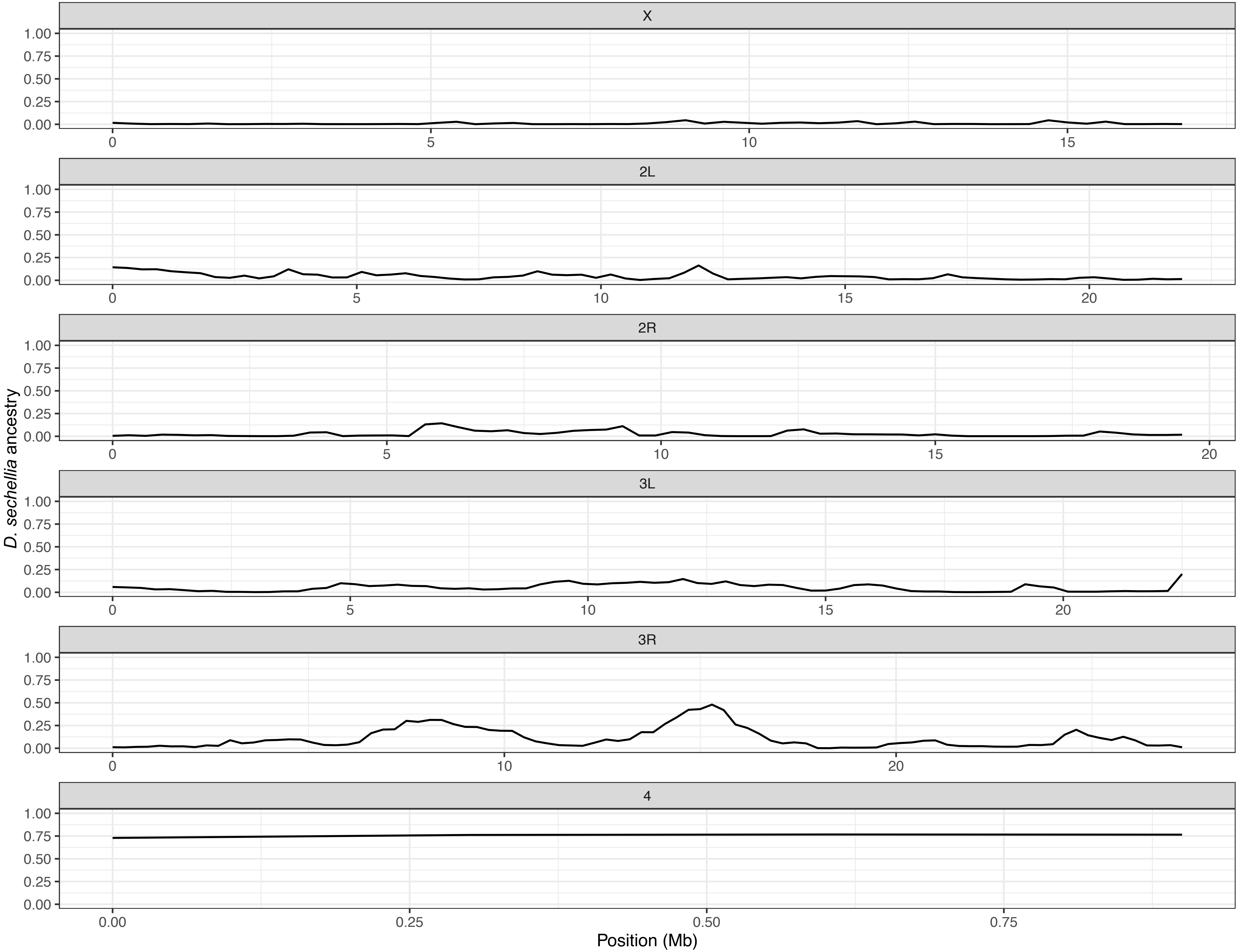


**Supplementary Figure 2. High-density sequencing reveals chromosome 4 and three chromosome 3R regions repeatedly retained in hybrid populations.** Genome-wide *D. sechellia* ancestry at generation 46 estimated from 673,183 ancestry-informative SNPs using non-overlapping 300 kb windows. Most of the genome retained little minority-ancestry DNA following intensive post-hybridization purging, whereas chromosome 4 and three chromosome 3R regions showed elevated *D. sechellia* ancestry in a predominantly *D. simulans* genetic background.

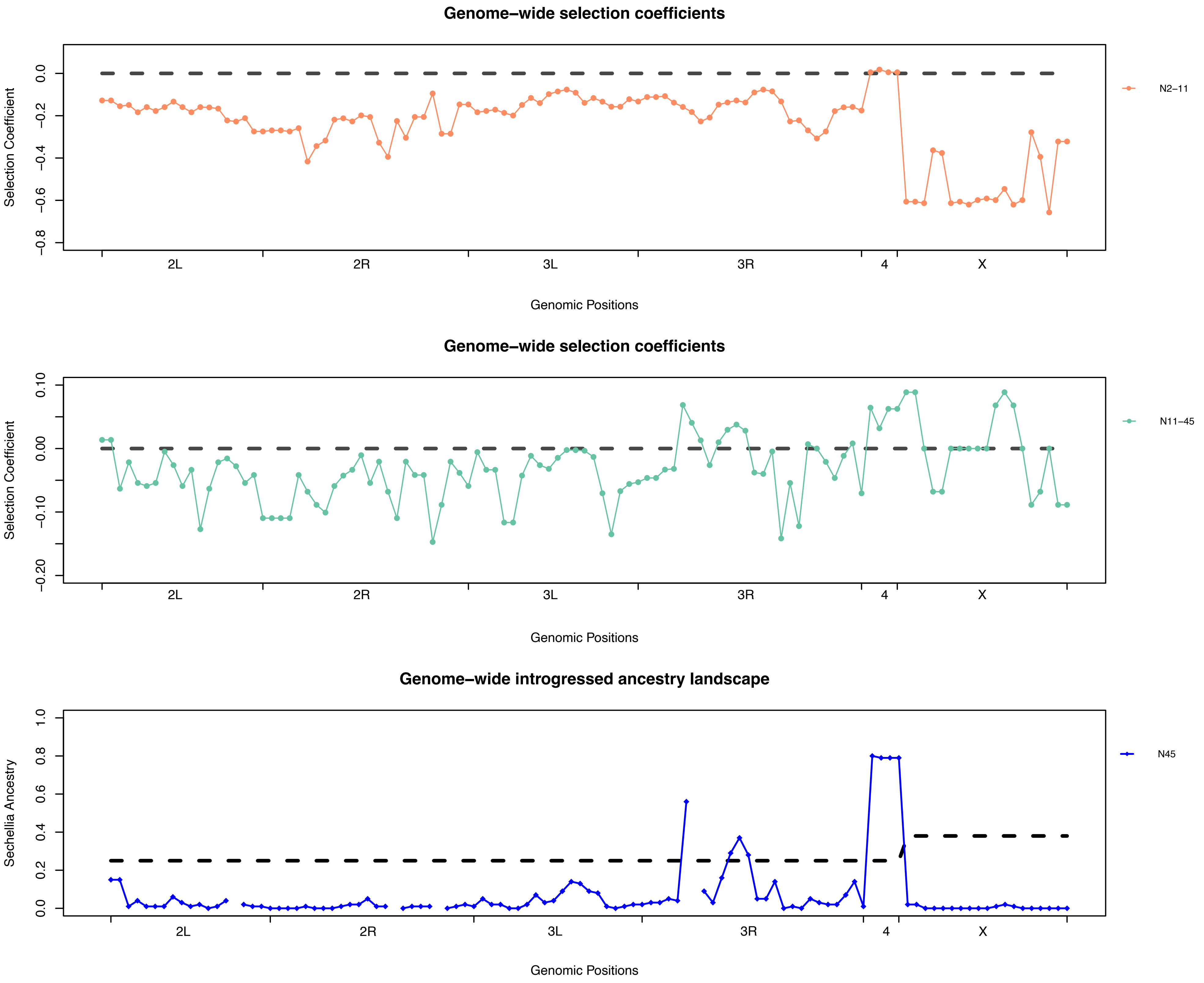


**Supplementary Figure 3. Genome-wide selection coefficients estimated during different phases of hybrid genome stabilization.** Top panel shows selection coefficients estimated from allele frequency changes between generations 2 and 11, during the initial phase of genome-wide purging. The middle panel shows selection coefficients estimated between generations 11 and 45, after most minority ancestry had been eliminated. Bottom panel shows the genome-wide *D. sechellia* ancestry landscape at generation 45 for comparison. Positive values indicate selection favoring *D. sechellia* ancestry, whereas negative values indicate purifying selection against *D. sechellia* ancestry.

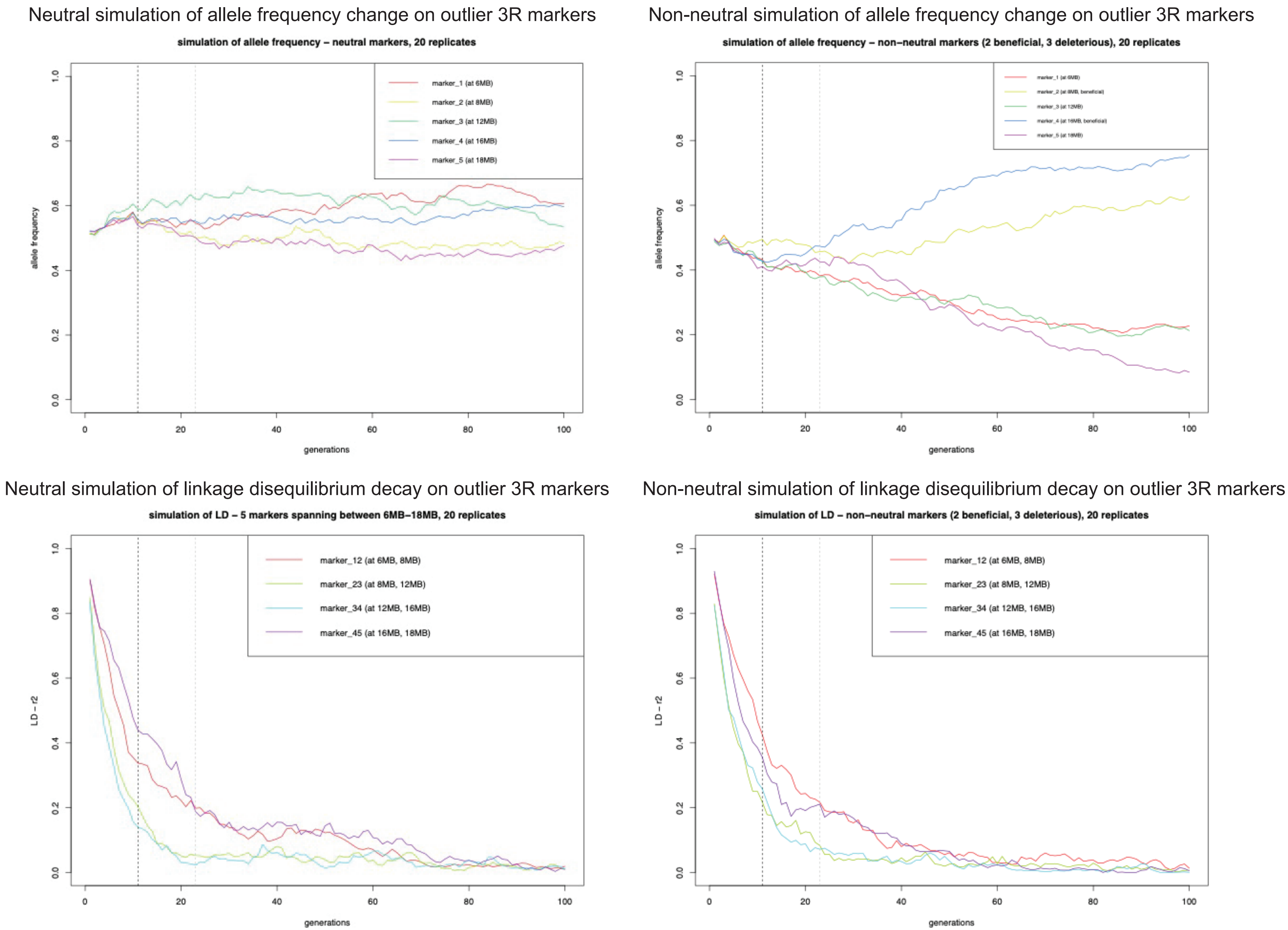


**Supplementary Figure 4. Simulations of allele frequency changes and linkage disequilibrium decay under neutral and selective conditions for chromosome 3R markers.** Top panels show simulated allele-frequency trajectories under neutral (left) and selective (right) models. Bottom panels show corresponding decay of linkage disequilibrium among simulated chromosome 3R markers. Under neutrality, allele frequencies fluctuate primarily by drift, whereas positive selection produces trajectories consistent with the empirical increase of the retained chromosome 3R regions despite continuing recombination.

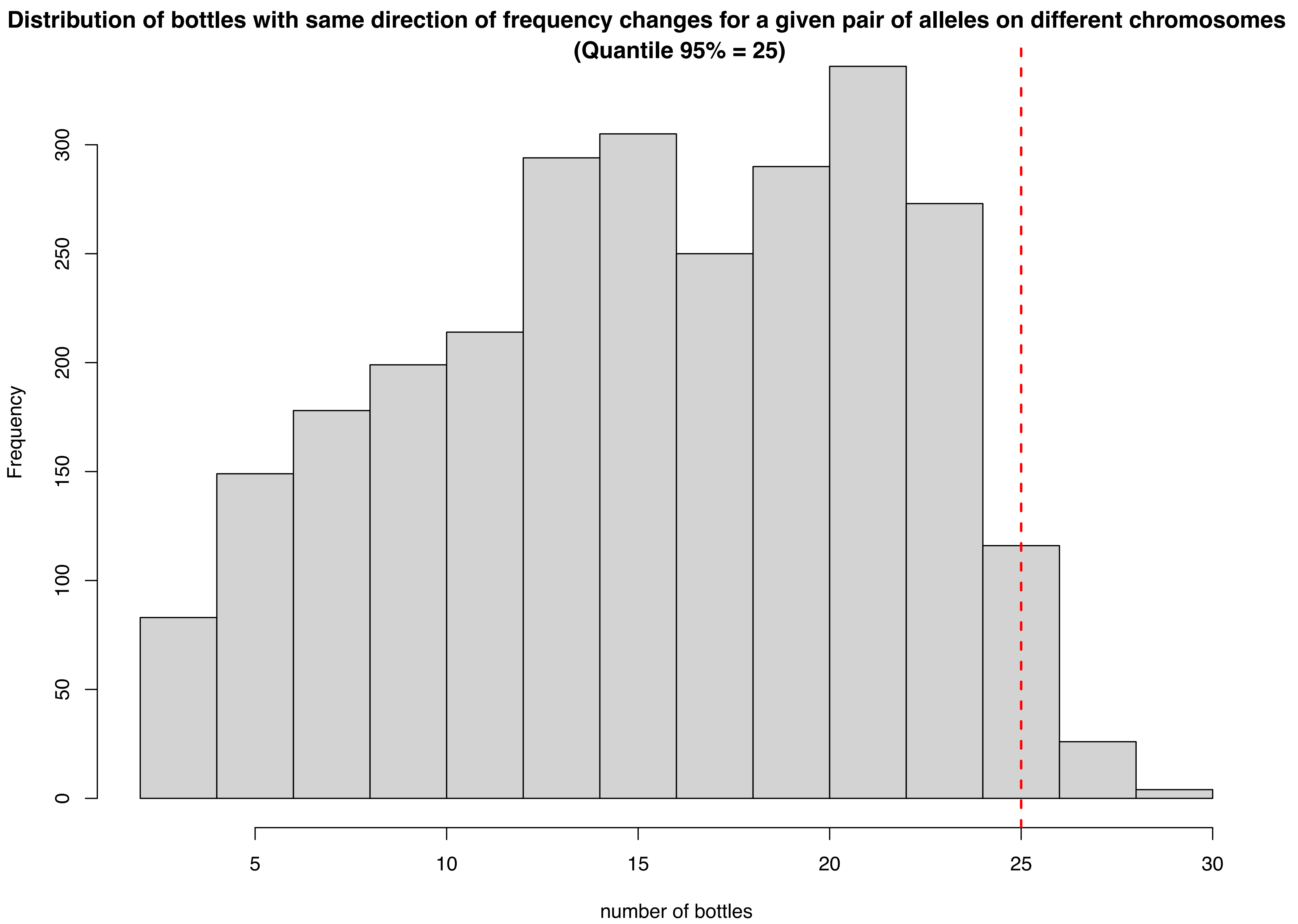


**Supplementary Figure 5. Distribution of coordinated allele-frequency changes among chromosome 4 and chromosome 3R loci across replicate populations.** For each pair of loci on different chromosomes, we counted the number of replicate populations in which allele frequencies changed in the same direction during generations 11–27. The histogram shows the genome-wide distribution of coordinated changes across locus pairs. The red dashed line indicates the empirical 95th percentile (25 populations), above which locus pairs were considered candidates for coordinated evolutionary dynamics.

### Supplementary Table 1

**F1 Hybrid Male Sterility**

| Virgin Female Strain | # of bottles set-up | # of progeny |
|---|---|---|
| *D. simulans* | 20 | 0 |
| *D. sechellia* | 17 | 0 |
| F1 Hybrid Females | 5 | 0 |

F1 hybrid males, from *D. sechellia* males crossed to *D. simulans* females, were crossed to three types of virgin females. In each case, ~30 virgin males and ~20 virgin females were included in each bottle. Consistent with previous reports of hybrid male sterility, none of the bottles produced any progeny.

**Supplementary Table 3**

Barcodes of sequences from each hybrid or parental population

*D. simulans* majority ancestry population #

| | fly bottle | barcode |
|---|---|---|
| 1 | A15.47 | CAGGTT |
| 2 | A25.47 | TCACAA |
| 3 | A35.47 | ACATCA |
| 4 | A55.47 | AGCGCA |
| 5 | A65.47 | CATCAA |
| 6 | A75.47 | GCTATT |
| 7 | B15.47 | TAGATC |
| 8 | B25.47 | CATGAC |
| 9 | B35.47 | GAATCG |
| 10 | B45.47 | TCTTCT |
| 11 | B55.47 | ATTCCG |
| 12 | B65.47 | GGAATT |
| 13 | C15.47 | ACGGTG |
| 14 | C25.47 | CTCAGC |
| 15 | C35.47 | TCCGGT |
| 16 | C45.47 | TGCAGT |
| 17 | C65.47 | TTCATA |
| 18 | D15.47 | ATACAC |
| 19 | D25.47 | CGTTAT |
| 20 | D35.47 | CTCGGA |
| 21 | F15.47 | TGTGTG |
| 22 | F25.47 | ACCGCG |
| 23 | F35.47 | GATCGG |
| 24 | F45.47 | TCACGG |
| 25 | F55.47 | ATTACT |
| 26 | G15.47 | CTTAGA |
| 27 | G25.47 | GCAGCT |
| 28 | G35.47 | TCCTCC |
| 29 | G45.47 | GAACTA |
| 30 | G55.47 | ACAACC |
| 31 | G65.47 | GGTAAC |
| 32 | H15.47 | GTGGTC |
| 33 | H25.47 | CCGCGT |
| 34 | H35.47 | CTGACA |
| 35 | H45.47 | CCGAAT |

| | | |
|---|---|---|
| 36 | H55.47 | AGCCGC |
| 37 | I15.47 | TAGCGC |
| 38 | I25.47 | TGACCT |
| 39 | I35.47 | CTTATC |
| 40 | I45.47 | GTAGCC |
| 41 | I55.47 | CCATAG |
| 42 | J15.47 | GAGGCA |
| 43 | J25.47 | AATTGA |
| 44 | J35.47 | ACTCAC |
| 45 | J45.47 | AAGTTG |
| 46 | J55.47 | CCTGTA |
| Parental | D. simulans | TAAGGT |
| Parental | D. sechellia | CCTCCA |

*D. sechellia* majority ancestry population #

| | | |
|---|---|---|
| 1 | A4E | TACGAT |
| 2 | B3E | CACCAC |
| 3 | C3E | GCATTC |
| 4 | D3E | TTCCAT |
| 5 | F3E | AGGCGA |
| 6 | F4E | CTCTCG |
| 7 | G2E | GCCTTG |
| 8 | H2E | AGGTTC |
| 9 | H3E | TAGCAG |
| 10 | I2E | CACTGC |
| 11 | I3E | AGTGGC |
| 12 | I6E | ATGTAG |
| 13 | J1E | GGTACG |
