## SupplementaryTable2 for "Hybrid Drosophila genomes rapidly purge minority ancestry but repeatedly retain selected regions"

**Supplementary Table 2**

Validated low-density markers with associated gene name, chromosome coordinate position, and sequence. For each marker, the species-specific nucleotide variant is indicated in brackets.

| Variant_GeneID | Chr | Position | Marker | Sequence with variant in [ ] |
|---|---|---|---|---|
| FBgn0002931dsimech1 | 2L | 0.08 | 2L_0.08 | ATTCACCCAATTCCAATGACCGTGATGCGGGTTTTTGTTCCGCCAGTTCCGAAAGCGAAGGCGGAGACGACTTGGTCGTCGAGCACGCGCGCAGT<br>GGCAGTCCCGACATCAGACCAAAAGGAACAGACTCAGCGGACTCCAAGCCAATAGCCCTTGTCCGCAACAAACGCAAGAGCTCGGAACCTTTCAA<br>AGTCGTAGGTTTGACGACTC[A/C]CAAGTCTAAGMCCATGCCTGATCCTCCGTCCAGTGCGAGTATGAATGCAACTGGTCCGCTAAAGAAGCGCATA<br>CGRTACACATCTTCCGCCGATTCGGCAGTTGTCCTAACACCACCAGCCATAGACTCTCCACCRCCCAACAGTTGCATACCCTCCGCGTTGCGCAT<br>GCAGCATGAAATAATGCCTAATCCRGCACATATATATGTTCGCCATCCGGGAGTTACGACTCTTCACAGATCGCCTGCTGCACACCCAGAACAGCTG<br>GAACCACTCGCATTAGTAACCACCAAAAAGCAATGTGTMGACAACGCGGGGCCTAAAATAGAGGCCTTTTCCACATTGCTAATTGGAAAACAGCCATC<br>CGCCAAAAAAACGTTGAAGGAAAGGACGCARAAAGAATCAACATCATCTTCATTTTTAGACGCTAGTCTTAGCGATGAGGATCTGAGCAAGACGGGATT<br>GGCATCAGTTTCGCGACCTCATCAACATCAACGCAATTACAAGAATATGACTAGGGAGCGCCGCATTGAGGCAAACGCACGTGAACGGACCCGTGT<br>ACACACCATATCAGCCGCCTACGAAACGCTCCGCCAAGCTGTTCCAGCCTATGCCAGCACTCAAAAACTTTCCAAACTCTCCGTTCTACGGGTGGC<br>GTGTTCCTATATACTAACCTTAAGCAGAATGGCTGGCGAGGATTACAGCGCCGATCAATCCGTACCGTCCATTGCCACGTGCTTAGAAGCTGTAACAT<br>CAACCATCCAAACGGAGGGAAAAGTGAAACGAAAGAAAGACGAG |
| FBgn0002931dsimech3 | 2L | 0.08 | 2L_0.08 | ATTCACCCAATTCCAATGACCGTGATGCGGGTTTTTGTTCCGCCAGTTCCGAAAGCGAAGGCGGAGACGACTTGGTCGTCGAGCACGCGCGCAGT<br>GGCAGTCCCGACATCAGACCAAAAGGAACAGACTCAGCGGACTCCAAGCCAATAGCCCTTGTCCGCAACAAACGCAAGAGCTCGGAACCTTTCAA<br>AGTCGTAGGTTTGACGACTCMCAAGTCTAAGMCCATGCCTGATCCTCCGTCCAGTGCGAGTATGAATGCAACTGGTCCGCTAAAGAAGCGCATACG[<br>A/G]TACACATCTTCCGCCGATTCGGCAGTTGTCCTAACACCACCAGCCATAGACTCTCCACCRCCCAACAGTTGCATACCCTCCGCGTTGCGCAT<br>GCAGCATGAAATAATGCCTAATCCRGCACATATATATGTTCGCCATCCGGGAGTTACGACTCTTCACAGATCGCCTGCTGCACACCCAGAACAGCTG<br>GAACCACTCGCATTAGTAACCACCAAAAAGCAATGTGTMGACAACGCGGGGCCTAAAATAGAGGCCTTTTCCACATTGCTAATTGGAAAACAGCCATC<br>CGCCAAAAAAACGTTGAAGGAAAGGACGCARAAAGAATCAACATCATCTTCATTTTTAGACGCTAGTCTTAGCGATGAGGATCTGAGCAAGACGGGATT<br>GGCATCAGTTTCGCGACCTCATCAACATCAACGCAATTACAAGAATATGACTAGGGAGCGCCGCATTGAGGCAAACGCACGTGAACGGACCCGTGT<br>ACACACCATATCAGCCGCCTACGAAACGCTCCGCCAAGCTGTTCCAGCCTATGCCAGCACTCAAAAACTTTCCAAACTCTCCGTTCTACGGGTGGC<br>GTGTTCCTATATACTAACCTTAAGCAGAATGGCTGGCGAGGATTACAGCGCCGATCAATCCGTACCGTCCATTGCCACGTGCTTAGAAGCTGTAACAT<br>CAACCATCCAAACGGAGGGAAAAGTGAAACGAAAGAAAGACGAG |
| FBgn0002936sechsim1 | 2L | 0.8 | 2L_0.8 | ATGAAGTCATTGCTGAATCGGATAATCCTGTGCAGCGCCTTTCTGGCCGTGGCCAGTGGTCTGAGCTTCACGGTGAC[G/T]TCGCGGATCTACATGG<br>ATGTGAAGCACAACAAGAAGCCGGTGGGCAGGATCACGTTTGGACTCTTCGGGAAGCTGGCWCCCAAGACGGTGGCCAACTTCCGGCACATTTG<br>CTTGCGCGGCATCAACGGGACCAGCTACGTGGGCTCGCGATTCCATCGCGTKGTGGACCGCTTCCTCGTCCAAGGCGGCGACATCCTGAACGG<br>CGACGGAACTGGCTCCATTAGCATCTATGGAGACTACTTTCCGGACGAGGAYAAGGCTCTGGCGGTGGAGCACAACAGACCCGGTTACTTGGGCAT<br>GGCCAATCGGGGTCCGGACACCAATGGCTGCCAGTTTTATGTGACCACCGTG |

| | | | | |
|---|---|---|---|---|
| FBgn0040105dsechsim9 | 2L | 02 | 2L_02 | ATGCTAAAATCCGCAATTGCTTTGCTATGTGGGCTTCTTGCTTTAAATTTGTGTGGAGCTTGGGCAGAATCTATGAGTAGTCCTTTGAAGGACCCGCCTAG<br>CCAGTGCGGAGGATACTGTCTTGGTGTACTCATGCCCGTGTTAAATCACCTGAACATTCCCCACAATCTGGCGAATTCAAGCAAGGCTAACGAGGTC<br>CTGTTGAGACAGTACACTATGGAAGGCCAACTGACTGCTCTGGAGGATAAGCAACTAAGCACTGAAGTCGCACTGGATGCCCAGGGCAGAAAACTG<br>GATATTATTAACGAACAGAACTTTACTGACCGACTGAATGGCTTGGAAAGCATACTGTCGGCAATGAAGGAANNNNNNNNNACATCATTCGTGTTGCG<br>GGCTNNNNNNNNNNNNNNNNNNNNNNNNNNNNNNNNNNNNNNNNNNNNNNNNNNNNNNNNNNNNNNNNNNNNNNNNNNNNNNNN<br>NNNNNTACGCGATGAGACAGCTGCTGGTTTACGCGTTGGTCGCGGGCTTGTTTTACTTTTCCGAAGGCAAGCCGTGCCCCCGAAAAGACACGCCCC<br>CACAAAACCCGCTGCGACGCCTTCTACAAGTGCCGGGAACTGCCCTCCAACTCGCACGTTTGGATTCCGGTGAAGTGCAACGAGGGACTGGTSTT<br>YGAGTCCAGTTTGGGCAGCTGCGTTCTGCCTGGCGAGGATTGGGAGTGCATCACCCCGAGTGAGATGAGCTCCACGCAGTCGCCACCTGGGAG<br>GCCGGACGCAGACATACTGATAGTCAACGACATGGACGAGGCGGGTCTGCTGTTGAGCGAGTCYAGCCACAAGGACGATGGCACCGTGGAAATA<br>GTTCTGGACTCCACACTGAGCAGTGAGGAGAATGAGGATAGTGAGAAGGCGGCCCAGAGCAAGYCGCAGCCGGAGGAGGACATMTCTTCCAGTC<br>AYAATTTCGACTCGAGYGGCGATGGGGAGCTGGTGGAACTGGATTCCCCAGCTGGTCTRAAGCCSGAGAACCGCGAGGAGGTGACTTCCAGCAGC<br>ATGGAGCACAGAACAGAGGTGGAAAAGCTAAGAACGGAACT[G/T]AAACAGGTGGCTGGTATCAGCGAAATAGCCAAACCGGGATCACCYATCGATC<br>CCAGCCTGACTGCTCACCTGCAGAGATTGTCCCAACTGATTGATGGACTGCAGCAGACCTACCAGAARACAGATAAGCCTCAGACGGAGATGCGTC<br>CAGACCAGTTGAATGCCTTCCTGGCACACTTTGACATCAAGAACCGGTATGAGATGATGAACCCCATGGAGCAGACCTCTTCCACGATCACCACCAC<br>CACCAGTACCACAGAAGCCCCCAAAATAAAGCCAAGGCTACCGGAACCCACCTCGAATAAGACCCGCCTGCAGGAGCATCTGAGCCACCATCGC<br>CTGGAGCCGGAGACGAAACTAATGCTCAGCAATGCCGTGCCCTACTCCTCGCATGGAACCACCGGGCAGGGATATGCCAATTCCCAGATAGTGGT<br>GAATCGCCCCGAAGGCTCTGTGATGTTTGCCCTGCCGCCCAATTACGGGATGAGCCAGGAACAGGGCTCGTATGTAAACCACCATTCCGCACAGA<br>GTCACGAATACAGCGATCATGAGCCCAAGATATCGGAGGACACCTTGAAGACGGTTCTGGAGCTATCCAAGCAGATGATCGCCGCACAAAACCTGC<br>CCAAGGTGCTGCCTAATCCGGGCTTCAATGGTGCCTATTACCAACCCATTCTGCAGCCTGTTTTCATGTCCCCCCAGGGCAATCCCTTCCAGCAGTA<br>CTACGGCATGCCGCCTCCCTTGAATCCGCACTACATGCAGCACAAGAAGCACTCCGCTGGAGGCGGTTCCTCGGGCAATGGCTACAACAAGCCC<br>AGCACCACCATTATTCACAACAATGTGATACCTGTTCACCTATCCAGCACGAGTTCGGGGGAAAAGGAGGTGCTGGACACGTATGGCAATAGTTTGGG<br>GGTATATCCCAGTATGGATAAGCATGTGACCGCCAGCCAGACCAGTGGCATGGAGTACATGACGACAGCTAGGCCCGTTACGATCGCAACCACTG<br>CCACCACCTACTCCGCTCAGTACGGATCCCAGTCCCCGTTTGCCAGTGACTACACCTTGACCACTCCGCGACCCTTCGAGCACCAACCTTCAGC<br>CGCCACAGTGGCCACCTACTACACTCCAAGTCCGCATCTGGGCGAGCACAACGCGAATAGGGTATACCAACCCACCGAGTCCTATCCCACGATG<br>AATATGCCCCGACCCATGGACATGTTTCCAGGACAAATCGATGCGGACAGGGTGAGCATTCGGCCCAACTCGCAAAAAATCGAAAGCATGGAGATG<br>GACGAAGATATGATTAAGGCCAACTATCAAAGTGGCTCTGGAATTGGAAATGGGAATGGGAACACTCTTCCTCACGTACTGACCTACAGCAGCGACAT<br>GAGCCAGCAATTAGCACCTTCGAATGGCTATCTGGAGCAGAGTTATAGCCACCAGCAGCCGCAGCACCACAACCATCCATATATGCCCAGGCCCA<br>GACCCACGATGAGCAGCAGTATCTATAGTGACTCGGATGGCAATATGGAGATGAGCATGTTCACCAGCAGTCCGAATCTCAACCCATTCAAGGGCGG |

| | | | | |
|---|---|---|---|---|
| FBgn0040105dsechsim9 | 2L | 02 | 2L_02 | ATGCTAAAATCCGCAATTGCTTTGCTATGTGGGCTTCTTGCTTTAAATTTGTGTGGAGCTTGGGCAGAATCTATGAGTAGTCCTTTGAAGGACCCGCCTAG<br>CCAGTGCGGAGGATACTGTCTTGGTGTACTCATGCCCGTGTTAAATCACCTGAACATTCCCCACAATCTGGCGAATTCAAGCAAGGCTAACGAGGTC<br>CTGTTGAGACAGTACACTATGGAAGGCCAACTGACTGCTCTGGAGGATAAGCAACTAAGCACTGAAGTCGCACTGGATGCCCAGGGCAGAAAACTG<br>GATATTATTAACGAACAGAACTTTACTGACCGACTGAATGGCTTGGAAAGCATACTGTCGGCAATGAAGGAANNNNNNNNNACATCATTCGTGTTGCG<br>GGCTNNNNNNNNNNNNNNNNNNNNNNNNNNNNNNNNNNNNNNNNNNNNNNNNNNNNNNNNNNNNNNNNNNNNNNNNNNNNNNNNNNNNNN<br>NNNNNTACGCGATGAGACAGCTGCTGGTTTACGCGTTGGTCGCGGGCTTGTTTTACTTTTCCGAAGGCAAGCCGTGCCCCGAAAAGACACGCCCC<br>CACAAAACCCGCTGCGACGCCTTCTACAAGTGCCGGGAACTGCCCTCCAACTCGCACGTTTGGATTCCGGTGAAGTGCAACGAGGGACTGGTSTT<br>YGAGTCCAGTTTGGGCAGCTGCGTTCTGCCTGGCGAGGATTGGGAGTGCATCACCCCGAGTGAGATGAGCTCCACGCAGTCGCCACCTGGGAG<br>GCCGGACGCAGACATACTGATAGTCAACGACATGGACGAGGCGGGTCTGCTGTTGAGCGAGTCYAGCCACAAGGACGATGGCACCGTGGAAATA<br>GTTCTGGACTCCACACTGAGCAGTGAGGAGAATGAGGATAGTGAGAAGGCGGCCCAGAGCAAGYCGCAGCCGGAGGAGGACATMTCTTCCAGTC<br>AYAATTTCGACTCGAGYGGCGATGGGGAGCTGGTGGAACTGGATTCCCCAGCTGGTCTRAAGCCSGAGAACCGCGAGGAGGTGACTTCCAGCAGC<br>ATGGAGCACAGAACAGAGGTGGAAAAGCTAAGAACGGAACT[G/T]AAACAGGTGGCTGGTATCAGCGAAATAGCCAAACCGGGATCACCYATCGATC<br>CCAGCCTGACTGCTCACCTGCAGAGATTGTCCCAACTGATTGATGGACTGCAGCAGACCTACCAGAARACAGATAAGCCTCAGACGGAGATGCGTC<br>CAGACCAGTTGAATGCCTTCCTGGCACACTTTGACATCAAGAACCGGTATGAGATGATGAACCCCATGGAGCAGACCTCTTCCACGATCACCACCAC<br>CACCAGTACCACAGAAGCCCCCAAAATAAAGCCAAGGCTACCGGAACCCACCTCGAATAAGACCCGCCTGCAGGAGCATCTGAGCCACCATCGC<br>CTGGAGCCGGAGACGAAACTAATGCTCAGCAATGCCGTGCCCTACTCCTCGCATGGAACCACCGGGCAGGGATATGCCAATTCCCAGATAGTGGT<br>GAATCGCCCCGAAGGCTCTGTGATGTTTGCCCTGCCGCCCAATTACGGGATGAGCCAGGAACAGGGCTCGTATGTAAACCACCATTCCGCACAGA<br>GTCACGAATACAGCGATCATGAGCCCAAGATATCGGAGGACACCTTGAAGACGGTTCTGGAGCTATCCAAGCAGATGATCGCCGCACAAAACCTGC<br>CCAAGGTGCTGCCTAATCCGGGCTTCAATGGTGCCTATTACCAACCCATTCTGCAGCCTGTTTTCATGTCCCCCCAGGGCAATCCCTTCCAGCAGTA<br>CTACGGCATGCCGCCTCCCTTGAATCCGCACTACATGCAGCACAAGAAGCACTCCGCTGGAGGCGGTTCCTCGGGCAATGGCTACAACAAGCCC<br>AGCACCACCATTATTCACAACAATGTGATACCTGTTCACCTATCCAGCACGAGTTCGGGGGAAAAGGAGGTGCTGGACACGTATGGCAATAGTTTGGG<br>GGTATATCCCAGTATGGATAAGCATGTGACCGCCAGCCAGACCAGTGGCATGGAGTACATGACGACAGCTAGGCCCGTTACGATCGCAACCACTG<br>CCACCACCTACTCCGCTCAGTACGGATCCCAGTCCCCGTTTGCCAGTGACTACACCTTGACCACTCCGCGACCCTTCGAGCACCAACCTTCAGC<br>CGCCACAGTGGCCACCTACTACACTCCAAGTCCGCATCTGGGCGAGCACAACGCGAATAGGGTATACCAACCCACCGAGTCCTATCCCACGATG<br>AATATGCCCCGACCCATGGACATGTTTCCAGGACAAATCGATGCGGACAGGGTGAGCATTCGGCCCAACTCGCAAAAAATCGAAAGCATGGAGATG<br>GACGAAGATATGATTAAGGCCAACTATCAAAGTGGCTCTGGAATTGGAAATGGGAATGGGAACACTCTTCCTCACGTACTGACCTACAGCAGCGACAT<br>GAGCCAGCAATTAGCACCTTCGAATGGCTATCTGGAGCAGAGTTATAGCCACCAGCAGCCGCAGCACCACAACCATCCATATATGCCCAGGCCCA<br>GACCCACGATGAGCAGCAGTATCTATAGTGACTCGGATGGCAATATGGAGATGAGCATGTTCACCAGCAGTCCGAATCTCAACCCATTCAAGGGCGG |
| FBgn0002989sechsim4 | 2L | 0'2.9 | 2L_0'2.9 | TCTCCATCGTCGCTGGTCAAGAACTGGGAGAAGGAGTTCACCAAGTGGCTTCAGGGTCGACTCCTTTGCCTRCCCATGGAGGGCGGGACCAAGGA<br>GAACACCATCCGAGCACTCGAGCAGTTCTCCATGACTTCTGCCCGKCTGGGGACACCTGTTCTGCTCATCAGCTACGAGACATTTCGCATCTATGC<br>GGAGATCCTGTGCMAGTAC[C/G]AGGTGGGAATGGTGATATGCGACGAGGGCCATCGTCTCAAGAACAGCGACAATCTCACGTATCAGGCTCTCAT<br>GGGTCTGAAGACAAAGCGACG |
| FBgn0000256sechsim1 | 2L | 03.8 | 2L_03.8 | ATGCGCAAGAGTGCAGTTAACCCGCCGAAGCCAATGCGTCCTTTATATTGGACACGGATAGTGACGAGTGCGCCGCCTGCGCCACGCCCCCC[G/<br>A]TCGGTGGCCAATTCCACGGACAGCACGGAGAACAGCGGGAGCTCACCCGATGAGCCTCCGGCTGCGAATGGTGSAGATGCTCCGCCCACAG<br>CGCCACCGGCCACCAAGGAGATCTGGACGGAGATCGAGGAAACGCCACTGGATAATATCGATGAGTTCACGGAGCTCTTTTCCCGCCAAGCCATT<br>GCSCCCGTTAGCAAGCCCAAGGAGCTGAAGGTCAAGCGAGCCAAGTCCATCAAGGTACTCGATCCCGAGAGATCGCGAAATGTGGGCATTATCTG<br>GCGGAGTTTGCATGTGCCGTCCAGCGAAATCGAGCATGCTATCTACCACATAGACACATCGGTGGTCAGTTTGGAGGCCCTG |
| FBgn0031562dsimech2 | 2L | 04.9 | 2L_04.9 | ATGTGCTTCAGATATAGCTTYATAGTATTCGCGGCTGTTCTGCTCCTGGCWGCCGGAATTCGGGCAGCTCCSTCGGATGTCGTCGTGGAGAAGGCG<br>GATGATCCCGCGGTGACCAATCCCCTGCCGGATGCCAGTGAGCTCACTGTKCCGGAGGACTGCCATCAGCCCAAGGAAACCGGTCGCTGTTTCG<br>CTCTGTTCTACCGCTA[C/T]GCYTACAAYGTGGATACYCAAWCCTGCGAGGAATTCGTCTACGGAGGATGTGCYGGCAACAAGAACAACTTCGARTCC<br>AAGGAGCAGTGCGAACAGGCCTGTCTGGGAAAGTCKGCGGGTTCCTCCRCGGACTCAACCACCGAACAAARCAGCGAAGTGGCCACAG |

| | | | | |
|---|---|---|---|---|
| FBgn0025472sechsim2 | 2L | 06 | 2L_06 | AAAGTGGCAGACGGCTATTGACAACACACTGTGGCTGAAGCACATTCGGTGCATTCTGGCCGGTGCCGTGAGGATCGTTGACAAGGTGGAGACCAT<br>GAGCACCTCCGTCGTAGT[G/T]CACTGCTCCGACGGCTGGGATCGCACAGCCCAGCTGACGGCMCTTTCCATGCTGCTGCTCGATCCGCACTAT<br>CGCACAGTGCGYGGTTTCGARGTTCTCATCGAGAAGGAGTGGCTCTCYTTTGGCCACAAGTTCCAGCAACGCATTGGTCACGGCGACAACAAGCAC<br>TCGGATGCAGACAGATCGCCGGTGTTYCTGCAATTCATCGATAGCGTCTGGCAGGTTAGCCAGCAGTTCAACAACGCGTTTGAGTTCAACGAGCACT<br>TCCTGATAA |
| FBgn0031878sechsim2 | 2L | 06.9 | 2L_06.9 | GTGAAAATGGACGAACCGAAGGATATGCGCTACAGTTTGGAGATCGACGAGGATTTGAGGAGACCCTTGGAGGCGCGGGCCAAGAAAGTTCTGTCC<br>GTTAATCCTGCCAGTCAAAGGACTGTTAATCTCGGCGAGGATACTATGTACGAAACAGGTRACTTGGAGCACTTGGACGATGGCGATGATAGCTTCAGT<br>GCCTTTGA[A/G]CGCATGTGCGACAAAACGGGCTCCGATCCGGATAGCACTCTTTTCCAGTACCTGCAYAGCGAGGAYCGGAGCCAGGTGATGGC<br>CAGGGCTAGGGATCACAGCACMGACGATCAGAAAGAGATCGATGCACTGCGTCGCATGCTAGAGTCAGTGGGTACAGAAGACGAYCTATTGGAGAA<br>CGAAAGTCCCGTCAAGGGGGCGGAATGCACTCACCTGGAGGACATTGAGGCTCCGTCGCGCTTTTGGGACAACACCCTGGCCGGAATCGACGA<br>GACCTCGTCCGAC |
| FBgn0031969sechsim3 | 2L | 07.9 | 2L_07.9 | AGACATTGCGCTTGTTCCGCTAATTGTCCTGCTGGGACAGATCGTAACCGGCATACTCTTGGCGGGCGGTCTCATATGCACCTGCTGGTACCCGAC<br>GCGRCAAGTGACGCAC[A/T]TCTGTCACAGCGATCCAAAGGCGAAGGCCAGTGTCCTGCGTCCACTGAACGCATTTGGTGTAAATTCGAMGGCAGC<br>CACAGCGCCGGTGGCCCAACTATTCCGGAACAACATTAGTTCATCGGGCAACGAGCGTGTGGGAGTGCGCCTGCTGGACTATAATCGCGATTCTG<br>GCATAAGACAGGAGAATGGTACCATGGAAAGCATGCACAG |
| FBgn0032170dsimech2 | 2L | 09.9 | 2L_09.9 | TCACAAGTCTTTCCGCAGTACKCCGACCTGGGACAGCGKCTGTTCAACCTGATGCACTCCAACTCCAAGGCCACAACGAAACATCTGGGCGCCGT<br>TGCTTTCCGACAGCAGTGCGAACGCTTTTTGGGCATCATGGATGATTCGAAGACGCTGGAGTGTTACATTAAGATGTACGCCCAGGAGGAGAATCCG<br>GACCAAAT[C/T]GACAAGACGGGAGTGACGCGGTTGCTCCACATCTGCTACACCATTGCCATGCAGCACTCGGGCAACGCMGTGCTGTGCCCGG<br>CCATCAACCGCACGTTCGGTTCGGTGACCAAATCCATTTTCCTGTGCCACGACAGTATGAGTCTGGGCTACGTTTGCCGCTGGTTCGAGCAGAACCT<br>GATGCGGCTGGTGC |
| FBgn0000018sechsim1 | 2L | 10.9 | 2L_10.9 | GCTGAATCTATGCCAGGGCGACTTTGGCCTCTACTATCTGCATCGGGAGTTCAGTGTGTTCCTGGAAGACGGTCGTTATGCCATGCTGGCTGCCATG<br>ACCGTTGTGCGCGG[C/T]GCTCTGCCGGTCGAAGACTACGTACGGTACCCAGATCTGTTCGACAAAGTGGATGCCTTCTCGTATGTGTTCTTTCTGGT<br>GGATTTGAAGCTMGGCGTCGTCACAGATAGACTAATCCTGCCCAACGACTCCATCGTCATTGCCCACAATCATGGGATTTCCGTGTTTGGCTCAACA<br>GTGATGATGATGTCCCGCCTGCATCAGTGCGTTTACGTCTATTGGGTGAATGACGGTAAGTTCCACCAGCAGGAGACGATTGGTCCCCGGCCGAGG<br>GACTTTA |
| FBgn0028542sechsim2 | 2L | 13.9 | 2L_13.9 | GACCTGCGAGAATGGCTTCTGTGCGAACGTAACGACGTGCGTRTGTCAGAACGGATATCGGTACGATCAGAATACGACGACCTGTCTGCCGGATTG<br>CGGGGACAACTGCGATAATGGGGTGTGCATTTCGCCGGGCAACTGCCGCTGCTTCAAGGGCTACGTCCGGAATCGGGAGCGATGCGAGGCKGT<br>CTGCGTGGGTGGATGTGGCTTCTACGGCAAGTGCATCGCACCGAATGTGTGCGGATGCGCCAT[A/T]GTCCCGGGTCCGGAACGCACYTACCAGC<br>GATGCGAGTATGGATTGTGCAACGCCATGGGAAGATGCCGCTGCCAGGTGGGCATGACGCGATTCATCGATCGGTGCATGTCGCCGGATACGGTC<br>ACCACATACGCCTCCATGAATCCCGTCAAGGTGAACGCATCGCTCATCCAGGAGTTTAATCTGCTGCTGGGCAGGCACTTCAACTTGACCACACTCA<br>GCGATATGTGGTGGTTG |
| FBgn0001974dsimech7 | 2L | 15 | 2L_15 | ATGAGCCTTAATTACGAACAAAATGCGGCAGACGAGCAATTCGCCCGCGCGCACAAGGCGGTCAGCCTGTCGGACGACGAGGAGAGTGAGGGC<br>CAAGCGGAGACCACGTCTKCGGGAAATCAGGAGCCGCATGTCAGTAAATCACCGCGGGAGTACTACGATGAGCCTGGCGGGAAGGGCAATGGAA<br>GTGGAGCAGATGATCAGGAGGAGCCGGAAACGGAGGCTGCCAGCGGAGCGGCCAAYACACATGTSGTGGCACACCACTACAACGAGCTGAAGG<br>AGGSGGGCCGCAAGGATCGCCAGAAGTCAAAGATCTTCTTCATGAGGAACTTTAACAACTGGATCAAGAGCCAGCTGATCAACGAGTACATGTCGAA<br>GATCAAGCAGAACAAACGCATGGGYGATGCTCTCAGAGTTCTGGACATGTGCTGCGGCAAGGGCGGTGACCTYCTCAAATGGGAGAAGGCTGCCAT<br>CTCCCACCTCATCTGCACGGACATCGC[A/C]GAGGTGTCGGTGGAGCAGTGCCAGCGGCGGTATCAGGACATCCTGCAGCGATCGGAGAAGTCG<br>AAGTTTGCCAATAAGTTCACCGCGGAGTTTTTTGCCTGCGACTCCACATTGGTGCGSCTGCGGGAGCGGTATAAGGATCCTTCGCTTCAGCTGAATCT<br>CGTGTCCTGTCAGTTTGCCTTTCACTATTGCTTCGAGTCAATGGCGCAGGCGGATTGCATGATGCGGAATGCAGCCGAGTGTCTGAAACCCGGTGG<br>CT |

| | | | | |
|---|---|---|---|---|
| FBgn0001974dsimech7 | 2L | 15 | 2L_15 | ATGAGCCTTAATTACGAACAAAATGCGGCAGACGAGCAATTCGCCCGCGCGCACAAGGCGGTCAGCCTGTCGGACGACGAGGAGAGTGAGGGCCAAGCGGAGACCACGTCTKCGGGAAATCAGGAGCCGCATGTCAGTAAATCACCGCGGGAGTACTACGATGAGCCTGGCGGGAAGGGCAATGGAAGTGGAGCAGATGATCAGGAGGAGCCGGAAACGGAGGCTGCCAGCGGAGCGGCCAAYACACATGTSGTGGCACACCACTACAACGAGCTGAAGGAGGSGGGCCGCAAGGATCGCCAGAAGTCAAAGATCTTCTTCATGAGGAACTTTAACAACTGGATCAAGAGCCAGCTGATCAACGAGTACATGTCGAAGATCAAGCAGAACAAACGCATGGGYGATGCTCTCAGAGTTCTGGACATGTGCTGCGGCAAGGGCGGTGACCTYCTCAAATGGGAGAAGGCTGCCATCTCCCACCTCATCTGCACGGACATCGC[A/C]GAGGTGTCGGTGGAGCAGTGCCAGCGGCGGTATCAGGACATCCTGCAGCGATCGGAGAAGTCGAAGTTTGCCAATAAGTTCACCGCGGAGTTTTTTGCCTGCGACTCCACATTGGTGCGSCTGCGGGAGCGGTATAAGGATCCTTCGCTTCAGCTGAATCTCGTGTCCTGTCAGTTTGCCTTTCACTATTGCTTCGAGTCAATGGCGCAGGCGGATTGCATGATGCGGAATGCAGCCGAGTGTCTGAAACCCGGTGGCT |
| FBgn0028858sechsim2 | 2L | 15.9 | 2L_15.9 | AAGTGGGAGGACCGCAATAAGCAGCCAGCCGCCACTGCCACGGAAATTGGACTTCAGTTCCAGTATCCACCTATCGAGAAAATGGATCCGATTCTCAATTCTCTTACCGAGTGCCAAAAGCTCAGCTTGTCCTCAAATATGATTGAGAAGATTACTGGAATTTCGGGAATGAAGAACCTGAGAGTCCTGAGCTTGGCACGTAACAATCTTAAAACGCTCAACGGCATAGAGCCACTGGCCGATACTCTGGAGGAATTGTGGGTCAGCTACAACAACATCGAGAAGATCAAGCCCCTGGAATCGATGAAGGCCCTACGTGTATTTTACATATCCTTYAATATGATCAAGGACTGGCCGGAGTT[C/T]ATGCGSATGGGTGTTCCACCAAATCTCAGCGAGATCACATTTGTGGGAAATCCTTTGAACGAGAACATGGACCAGTCTGCCTTCACTGCCGAGGCAGTCCGTCGTCTGCCCAACATGAAGAAACTCGATGGGGAGCCCGTCATTCGT |
| FBgn0032624sechsim1 | 2L | 17 | 2L_17 | AAATCGCATTGATCACGAGGAAAGGCGCCGCGAACACGAAAGAAAAGTCGCTGTGGAAGCCTTAAAGTTGGCCGAACAGCGCCGAATGAGGGATCCTGTAATTGCCCCGACGGAGGTCAATGAAAACTCGAAGAACGTGCGACCAATTATAAATATCTCACGATCTCCAATAAAATTCACGCCAGAGGAAAGGATCA[A/G]AGTGRACCGTTTRAGGGCAGCGAAGAGAGAGCGTATCGAGCGTATCCTTCGCGAAATGACAAACAGCAAGCAG |
| FBgn0032663sechsim1 | 2L | 18 | 2L_18 | ATGGATTTCCACCTGCTGATGATTGCACTCGTGCTTGCACCAGTCTTTACACCTGTCCACTGCCAGGTGGCCTGCAATGCGAGGAGCGGCCCTGTCCGCCAGAAACGCATCGTTCTCAGCAGCCCGGGAACAGGTGGGCTGGCCGAGTTGGACACTTGCCAGTACCGCGTGGCGCC[G/A]TGGAGCGGCCAGGTGTGTCAGGTGCGCGTTGATTTCGAGCGTTTGGAGCTGCCGCAGCCGCARCTGAATGCCAATTCCGGAGTGCTGGAGTGCGTCGATTTTCTGCAGATACAGCATTTCCGGCTGTGCGGTCGCAACAGTGGTCAGCACTTGTATATACCCATTCGCCGGGGTCAGGAGCTGCAGCTTCTGTTCCGACTGGCCWCGTCCCTCGGCAGGCAGTCCACCTGGCAGTTGACTCTCACCCAACTGGAGTGYCCCAGGGACTCCGTCAGRGCTGTAA |
| FBgn0032838dsimech7 | 2L | 19.9 | 2L_19.9 | GTCCAGCAGCACTATCGTTACCAGAAGGAGAAATCCGATCARATGAACTCGTTGACCTCGCAGCTCTTGCTCCTGGAATCCCGACTGAAGCGGAAGCAGAAGCAGATAGCSAGTCTATTGAACCATCGGGAGTTGACCATCCAGCGTCARCAGAAGATYATCGACMCACTGTCCTCCCGGCTGGTGGATCA[C/T]GGGCTGGAGACCATTGAAGCTAGCTATGCCAACGAGYTGGACTCCCTGAACGACTCGGATTCGGCGGTGGTSCTGGAGGACATCGACTCGGACAGCCCRATGACATTGGGCACGCGACGGAAGAGTAGTGGCGCTGGCGGATTGGGCGGCGTGCTGGGCAGCGATGGCATCACCATAGTGCGGTCCATATCCGATGCAATCGAGACGAATCACAACAA |
| FBgn0032971sechsim2 | 2L | 22 | 2L_22 | GAGTCCATGCGAGAAAATATGCCTTAATACAGCTAGAAAAACAGTACAGAGATGCGACAAAAACTATTCACCACCAAAGTTAAGTAGAATCAAGAATTTCTATACCTACAGTWTTGTACTGGGGAGTCTCTTC[A/G]GTATGGTCATGTGGGCCATCTATAAGCTGGGCAAGCCCGAGGAAGACCATCGGGGTCCCATAGAAGATGAGTTCAGTCAGCTTCCCTGGTTCCGCCAGTACWTAATGAGGATGTGGCACACACTTCAGTATTATGAAAAAATGATGGAAGAACCCCAGATGGCAAAGCTGCTGCCAAACGTTGTCCCACCGCCGTATATACAGCCTCCCTATTCACTAGTCCTGGAAATCAAAGACGTACTGGTGCATCCAGATTGGACTTATCAGACGGGGTGGCGGTTTAAAAAGCGACCTGGTGTGGATTACTTTCTTCAGCAGTGCAGTCGAAATTTTGAAATTGTTATATATACTTCAGAACAGGGTATGACCGCATTTCCCCTTCTGGATGCTTTAGATCCCTATGGCTACATAAGTTATCGGCTAGTAAGAGGTGCCACGGACTTGGTGGAGGGTCAGCATACCAAGAACCTCGATTATCTTAACAGGGATCTGAGCCGTGTGATAGTGGTGGACTGTGATCCGTATACCACTCCCATGCATCCCGATAATAGCTTGGTGCTCACAAAGTGGCTGGGTAACGATGATGATGTCCAGCTCTTTGATCTCACGGCTTTCCTGCAGTTGGTTGCCGAACACCAAGTAAATGATGTAAGGGAGGTGCTGCGATACTATCGCCAGTTCGAGGACCCCATTGAG |
| FBgn0040849dsimech1 | 2R | 0.8 | 2R_0.8 | GCAGATATATGCATTGGAGCTATGTACTCGTGGTACGAAGATTACACATACTTAGACCTTTCCATGTATCTTGTACGTTCTGGAATAACGTGTCTTGTACCAGCGCCATTGCGACTGACTAGTTGGTACCTTCCCTTAGAGCCTTTCAAAGAAACTTTATGGGCTGCAATTCTRTTATGTCTATGTGCGGAAGCTACAGGRTTGGTTTTAGCATTTAAAAGTGAGCAGGCGCTGTATGTACTGCCTAGCTACCGTGAGGGCTGGTGGACTTGTATAAGCTTTGG[T/A]GTATGTACCACCTTTAARCTTTTCATATCGCAATCAGGAAACAGCAAGGCATATTCACTGACAGTTCGTGTACTACTATTTGCCTGCTTCCTTAATGATTTAATAATAACCAGCATATATGGTGGCGGCCTTGCAAGTATATTGACAATTCC |

| | | | | |
|---|---|---|---|---|
| FBgn0033005sechsim1 | 2R | 01 | 2R_01 | TATAGAACCAAATTTTTCCGGTGGGTACAATCAAACCACTGGATATTCTTCGGATACTAAGGACACAACTTTCGTCGTTGGATTACAAGACCTTCGAGTTGAAGACTTCAAAAAATGCATTGAAATTTTTGACAAAACCATTATTAATTCAATGAATGTCGGGTTTGATTCCCAGCATGTTGAAAGTGTTCTGCACAATCTTGAGTTGTCTTTAAAGCACCAGAATCCGAACTTTGGAAATAC[T/C]TTGCTTTTTAATTCCACAGCATTATGGAATCATGATGGCGATGTAGTCAGTAATCTCCGCGTATCAGATATGATATCAGCACTTCGGGGAAGCATAAGCCAGAATAAAAACTATTTCCAAGAAAAARTAGAGAAATATTTCGCCAATAACAATCATAGACTAACATTAACTATGTCACCTGATGAGGCATATGAGGATAAATTCAAACAGGCCGAGTTGGAGTTGGTGGAACAAAAAGTAAAACTTTTGG |
| FBgn0033056sechsim1 | 2R | 01.9 | 2R_01.9 | ACCTCGGGTCAGCGGAACAACACCAGCACGGCCAGCAACAGTTCCCTAAACCAACGAAATAAGGAGGACTCCGCGGAGAGGTTGTCAGTGA[A/G]CGAAGATACGGCAGCCGTGGACAGCAGCGCCTTGCTTCAAAGTGGATCCTCAAACTCTCGTGATACAGTTAACTTCGATTCGACCTCCACCTCGATCAGGACGTTCCTTAAGAACACAGTGGCCCTGGTCAGGGCAGGAGAGTCCTTCACATCTGTAAAACATCTCTTTCACAAGTTCATCAAACGCAGCGGCTCTGTGTCAAATTCCAAGCCTTCGCTGACATCCTCCTGCTCCCAGGCGGTRGGCGAGGAGAATCCCTACATGAGCAAGTCCAATGACAMGCTTCTGACGGACTCGCCCATCCGCACTACTGTGCCCAGCACCTCCCAGATGACGCCCGCCTCACATTTCTCCAA |
| FBgn0028642sechsim1 | 2R | 03 | 2R_03 | CAACGAGCCGGTGCGGGAGCTGCCCCACGATGGCTACGAGCAGCTCTTCGCTAACAATCGCAACCAGGAGCACCCGGCCGAACAGGATTTGGATGACGAGCAGCTGGACAACCGACCGCTGAAGGAGGTGCGCTTTCACAGCGTCCAGGACACTATGTCGCGCTCCAAGAGCTACACRGATAACTCCAATGCCCGGCGGCGCAGGAGGCGTCGCAACCAGTCGCGCAGCTCCTCGGAGATGCAGATCAACCAGACGAACCTCCGCCTGCA[T/C]AACGCACAAACGCAGGTGGGAACGACGCCCCTCAATCTGCTGAACAACCTGGACAACTGCGACGTGGCCAGCATCTGCTCCACCTGCTCCTCCAGCTCCTCGAGCGACATGGACGACTACGTGTACCGCCTGCCTGCGCGGAAGCACTACGGCGGAGTGAGAGTGGCCTATGTGCCCAATGATGCGCTGGCCTACGAGCGCAAGAAGAAGATGGCCCAGGACTCGTCCCTGGCTCCCGGGGGAGGCAATGCATCCGTGGGCGGGGCACCGGCGATCATGCACGAGAGCAAGAACTGCACGATTTCC |
| FBgn0033259sechsim1 | 2R | 03.9 | 2R_03.9 | TGATGGCTGTAATGGCATTGGTTACTATCGTCTCCCTGGTCATCATTCTGCCTGTTAACTTCCTGAACGGGCCCAA[A/G]GAAAATCCGTACGATGTGAATGCCTTCGGCCGCACTACAATGGCGAATCTTTCGCCGGATTCGCCTTGGCTGTGGGTGCACACGATCATCACGATTCTGTACATTCCGCTCGTAGTGCTCATCATGCGGCGCGCCTCGGGACGGAATGCCTTCAAAAAGGCAGCCACCCGCACGATTATGATTTCAAACATATCCTCCAGCGATCGTAACAAGACTGTGGTACGGAACTATATGCAGGAGCTCTTTCCRGACGTCACCATCGAGAATGTGTCAATTGCGTACAACATTTCACGACTCTACGTAAGGAACGCCGAGTTCGAGCGAGTTCACGAAGCACGATTGTACTGCGAGCAGCATAGGAACAGGGATACTCTCATG |
| FBgn0033369dsimech2 | 2R | 04.9 | 2R_04.9 | CGAGCACCTGGCCCGGACCAACTCGAAATTCCGGCGCCTGCGAATCGCCTGCAACAAGATCGGTCCGGATGGAGCCTTCTTCCTGCTRCGTGGAATGTCCATGGGTGGCAATCTGGAGYTGCTGGATCTGAGCTACTGCTCGATTGGCACGCAYGGCGGYGAGTGGGTGGCCAAGTACCTGGCCTCGTGCCGCAGGCTGCAGATCCTGCACTTGAACTACAACGACATG[A/G]GCCCGACGGCAGTGAATCTCATCCTTCTAACCATGAAGAAGCAGTGCAAGCTGGAAAGACTAACCTTGTATGGCAACCASTTTGACTCGCGTACTGCGATGATAGTGCGTCGCTTGCTGGATGCYGAGGTWGTGCTCCACTCCGAGCTGGACATCAGTTACACCTACGACGAGGCACTGCAGGATTACAGGGTGGTGCCTTGGCG |
| FBgn0033369dsimech2 | 2R | 04.9 | 2R_04.9 | CGAGCACCTGGCCCGGACCAACTCGAAATTCCGGCGCCTGCGAATCGCCTGCAACAAGATCGGTCCGGATGGAGCCTTCTTCCTGCTRCGTGGAATGTCCATGGGTGGCAATCTGGAGYTGCTGGATCTGAGCTACTGCTCGATTGGCACGCAYGGCGGYGAGTGGGTGGCCAAGTACCTGGCCTCGTGCCGCAGGCTGCAGATCCTGCACTTGAACTACAACGACATG[A/G]GCCCGACGGCAGTGAATCTCATCCTTCTAACCATGAAGAAGCAGTGCAAGCTGGAAAGACTAACCTTGTATGGCAACCASTTTGACTCGCGTACTGCGATGATAGTGCGTCGCTTGCTGGATGCYGAGGTWGTGCTCCACTCCGAGCTGGACATCAGTTACACCTACGACGAGGCACTGCAGGATTACAGGGTGGTGCCTTGGCG |
| FBgn0001291sechsim1 | 2R | 05.9 | 2R_05.9 | GGAGAAGAAGGCCCCATGTCGCTGGACTTTCAGTCGCCGAACCYGARCACATCCACCCCGAATCCTAACAAGCGTCCCGGCTCGCT[G/T]GATCTGAACAGCAAGGGTGCCAAGAACAAGCGCATCTTCGCACCACTGGTCATCAACTCACCGGATCTGTCATCCAAGACGGTRAACACACCTGATTTGGAGAAGATCCTGCTGTCCAACAATCTGATGCAAACACCGCAGCCGGGCAAGGTGTTCCCCACCAAGGCGGGGCCCGTCACCGTGGAGCAGTTGGACTTCGGCAGGGGATTCGAGGAGGCCCTACACAATCTCCACACTAACTCCCAGGCATTTCCG |
| FBgn0033581sechsim1 | 2R | 07 | 2R_07 | ATGGAAGCCGACATAACGCCCCTGGCGGAAACCAACGGAGAGCAGCGCCAAAATCCCCAGGAAAGCGCTCAAAACCCAGAGCAACCGCAACCGCG[G/A]CAGGATTTGGCGGGCAAGTCGGTGGGTTCCGCTGATGATGGCGTCTCCGGAACTGTGGGACAGATTATGGATTACCTCGAGGACGATGCCGGTGCAACTGCCGACCAGGACGCGGAGATGGGCGAGGCGGAGTATCTGGATGGCGAGATGCCGCAGGAGGATGAGAGMGAGACTCCGCAAACGGAGAAAGCCAATGAAGGTTGCCAGGAATCAAGCAATAAGAAGGATTTGCAAGAATCTGCAGCTAAAGATAATGTTTCCAAGGAAAAAACTGGCGAGGATGGTGAAAAAGACACCGAA |

| | | | | |
|---|---|---|---|---|
| FBgn0027495sechsim1 | 2R | 08 | 2R_08 | GCTCTAGAGCTGTACAACAAAAGCATCTGCTATGCCGAACCGAACTCGGAGCACCTGTCCATCGGTTATGCTAATCGGTCGGCGGTACTATTCGAGTGGAAGCGCTACCGGCAATGCCTGGACAATATCGAGCTGGCAAGGCAGGCCAACTATCCGGCGAGACTGAGCCACAAGCTGGACAAGCGGGAAAGGGACTGCCAACAGCTACTCGATCAGCAACCGCCGGATGTGGTGCCCTATGAGTTTAAGCTCAGTTTCGAAC[C/A]GCATGCCCAGGTGCCCTTCATCGCCRACTGTTTGGAGTTGCGCGAAACAKCCGCCGAGGGTCGCTTTGTGGTGACCAATCGGGATTTGGTCGTCGGGGATCTTGTGTGCGTGGAGGAGCCCTTCTGCTCCACACTGTTGACGCCCATGCGATACATTCGGTGTGCCACCTGCAAGCGGGAGAACTACCTGACCCTCATACCCTGCGACAGCTGCTGTTCCACGATGTTCTGCTCCGAGGAGTGCAAGTCGATTGCCATGCAAACCTATCACCGCTTCGAGTGTCCCAT |
| FBgn0027495sechsim2 | 2R | 08 | 2R_08 | GCTCTAGAGCTGTACAACAAAAGCATCTGCTATGCCGAACCGAACTCGGAGCACCTGTCCATCGGTTATGCTAATCGGTCGGCGGTACTATTCGAGTGGAAGCGCTACCGGCAATGCCTGGACAATATCGAGCTGGCAAGGCAGGCCAACTATCCGGCGAGACTGAGCCACAAGCTGGACAAGCGGGAAAGGGACTGCCAACAGCTACTCGATCAGCAACCGCCGGATGTGGTGCCCTATGAGTTTAAGCTCAGTTTCGAACMGCATGCCCAGGTGCCCTTCATCGCC[A/G]ACTGTTTGGAGTTGCGCGAAACAKCCGCCGAGGGTCGCTTTGTGGTGACCAATCGGGATTTGGTCGTCGGGGATCTTGTGTGCGTGGAGGAGCCCTTCTGCTCCACACTGTTGACGCCCATGCGATACATTCGGTGTGCCACCTGCAAGCGGGAGAACTACCTGACCCTCATACCCTGCGACAGCTGCTGTTCCACGATGTTCTGCTCCGAGGAGTGCAAGTCGATTGCCATGCAAACCTATCACCGCTTCGAGTGTCCCAT |
| FBgn0020391sechsim1 | 2R | 09 | 2R_09 | GGACTTGTTCCCGATCAGGAAGATCTCCTAGTCGCTGTTAAGATGCTAAAGGACGACGCCAGCGACCAGATGCAGATGGATTTCGAGCGTGAGGCCTGTTTGCTGGCCGAGTTCGATCATCC[C/A]AATATCGTGAGGCTSCTGGGGGTGTGCGCCTTGGGCAGACCCATGTGCCTGCTCTTCGAGTAYATGGCTCCTGGCGATCTSAGCGAGTTCTTRCGCGCCTGCTCCCCATATGCCACCCACCAGGCGCCGACTCGGGATCGTCTGCAGTTGAACGAGCTACATCTGCTGCAGATGGCGGCCAACATTGCAGCGGGMATGCTGTATCTTTCGG |
| FBgn0002643sechsim3 | 2R | 09.9 | 2R_09.9 | ATGAATATTTGGCAACAAAAATTCCTGAAACGACCCGCGGAGGATGTTGACAATGGCCCTGACAGCTTTGAACCGCCGCACAAGTTGCCTAATAATAACAACAATAGCAATAGCAACAACAACAACGGCAATGCGAATGCCAACAACGGCGGMAACAACAATGGCTCCAATACCGGCAACAACACGAACAACAACGGCAACAGCACCAACAACAACGGCGGCAGCAACAACAACGGCTCTGAGAATCTAACCAARTTCTCGGTGGAAATTGTTCAGCAACTGGAGTTCACCACATCGGCGGCCAATTCGCAACCGCAACAGATTAGCACTAATGTCACTGTAAAGGCGCTGACCAACACATCGGTGAAGAGTGAACCAGGTGTGGGCGGTGGCGGAGGGGGCGGCGGTGGTGGCGGTAGCGGTAACAACAA[C/T]AACAACGGAGGCGGGGGAGGCGGCGGAAACGGTAACAACAACAACGGAGGTGATCATCACC |
| FBgn0001187sechsim1 | 2R | 11.1 | 2R_11.1 | AAAAGTCTCTCAACCGATCGGAGCAGCTGTTCGAAAAGATGACGGCGGGCATGTACTTGGGCAACCTGGTGCGCCTTGTCCTACTTCGTGCCCTCGAGCGCAAGCTGATCTTTAAACAGAGCAGTCGACGTCCAGAATTCGCCAGCGTGCTGCAGAGAAACGAAGAGGTTTTCGAGACGCGCTACATATCGGAAATCGAGGACGACTCCTTTCCGGAGTTTGCCAGCACAAGGAAGATTGTGAAAAATCTGTTTGG[C/A]TTGGAGAAGGCCTCCGTGGAGGACTGCCAGACTCTGAGATACATCTGCGAGTGCGTAGCTAAGCGAGCGGCCACATTAGTAGCCATCGGAGTGAGTGGRCTGGTGAACAGGACCTCCAACCGCCGCGTGATAGTCGGCATGGATGGCTCCGTCTACAGATACCATCCAAAGTTYGATGCCTACATGAGGCAGACACTCCAAAAGCTGGTGAAAGCAGACAAGGAGTGGGACATCATGCTATCTGAGGACGGATCCGGACGTGGAGCTGCGCTGGTGGCCGCGGTGGCCAGTAAAACAAAG |
| FBgn0029175sechsim3 | 2R | 12 | 2R_12 | TCGCTACTACACGTCCAATCCGAACGAAGCGTGCCTGTTTCTGCCCAGTTTGGATCTGCTGAACCAGAATGTCTTCGACAAGCACTTGGCGGGCGCKGCCTTGGCTTCCCTGGACTTCTGGGATCGTGGTGCCAACCACMTCATCTTTAACATGCTGCCCGGCGG[A/T]GCTCCATCCTATAACACTGTGCTGGACGTGAATACGGATAATGCCATCATTTTTGGCGGTGGCTTTGACTCTTGGTCTTATCGCCCGGGCTTTGATGTCGCCATTCCAGTCTGGAGTCCACGATTGGTGCGGCAACATGCACATGCGACGGC |
| FBgn0010591sechsim1 | 2R | 12.9 | 2R_12.9 | GCCAAGAAGATTCCAGCTGTGCGTCGTCAGGTGGAGACTGAATTGGCCAAGGCCAAAAACGACTTCGAGACYGAAATCAAGAAGAGCAACGCCCATCTTACCTACTCGGAAACTCTGCCCGAGAAGGGACTCAGCAAGGAGGAGATCCTCCGACTGGTGGATGAGCACCTGAAGACCGGTCACTACAACTGGCGTGATGGTCGTGT[T/A]TCTGGCGCAGTCTACGGCTACAAGCCCGATCTGGTGGAGCTCGTCACTGAAGTGTACGGCAAGGCCTCCTACACCAATCCCTTGCACGCAGATCTTTTCCCGGGAGTTTGCAAAATGGAGGCGGAGGTAGTGCGCATGGCATGCAACCTGTTCCATGGAAACTCAGCCAGCTGTGGAACCATGACCACCGGSGGTACCGAATCTATTGTAATGGCCATGAAGGCGTACAGGGATTACGCTAGAGAGCACAAGGGAATCACCAGGCCGAACATCGTGGTGCCTAAGACGGTCCACGCGGCCTTCGACAAGGGCGGTCAGTACTTTAATATCC |

| | | | | |
|---|---|---|---|---|
| FBgn0000057sechsim1 | 2R | 14 | 2R_14 | TCGCTGCCAACCGAAATAGAGGTGCATAAGAAGGAGGGCAATAACTTTCTTGAGAATGGCAAACTTGTGGATGCAATTGATGCGTACTCAGCGGCTTTG<br>GCGAAATATCCACAAGGAGAGGTCCTCTATCTGAACAGAGCGACGGCTCTAAT[G/A]CGACGCGGCTGGTTTGGTGACATATATGCGGCACTTAGGG<br>ATTGTCAYGAGGCCCTGCGTCTGGATCCCAGTTATGTGAAGGCCCACTTTCGGCTGGCCAGGGCATTGCTTGAGCTGCATCGACCCCAGGATGCG<br>GATCAGTGCTTGCAGGCACTCATTCAGCGTTTCCCCAACTTCGCAAACAATCACGGCGTGCTCATGCTGAACAAGGACATCAAGGAGAATCGACGAC<br>AGACACAGAGTCCCGATGCACCCGAGCCACAGCCAGTGGAAAYGGATGATGGCTTACACTACATGCGTCTGAAGGACGCCGAATATGATCTCCGTT<br>CCAMGGCGCGCGACTATATGCAACGCTATGTGGGCCACTGCAACATCACGACGGACATAAAGGAGGCCAACTATTTGGGCAGCCAGGGCGAGTT<br>CATAGCCGCTGGTTCGGATGACGGCAACATGTACATTTGGGAGGGAGACACGGGCAAGATACGCGCCGTTTATCGGGCGGACAGTGCCATCGTGA<br>ACTGCGTGCAACCGCATCCGAGCATTTGCATGCTGGCCACCAGTGGCATTGACCATAGCATTAAGATCTGGTCGCCGTGCGCCGCCAGCGCCGA<br>GGAGCGACCGAATCTCGTGGCGGATGTGACGCGCTTCGTTGAGGACAACCAGGAGAAGATGCGCCTCGATCCATTCGAACTGCACACGCGCAAT<br>GCGTACTGTTTCAATAAT |
| FBgn0027535sechsim3 | 2R | 15 | 2R_15 | TTTTACCCAACACACTGCCCAGAAAGAATAATCCCATCACTGGAGGGATGGCGGCCAGCTGCGAAATGCACAACTGCTTCAATCACTCCCGCTGCA<br>GYCTAACCTCCGGATTYCCTGTTTATCTGTACGATCCGGATGAGCATAGCGTACAGCGAAAGGGTTACGATATCGATGGTTTCCTGAAGACGACTCTAA<br>AGCAGACTTTGGGATACAATGCTCATATAGTTAAGGATCCCAAGCATGCCTGCATATATCTAGTGCTGGTGGGAGAGGCTCTGCTTGAGCAGGATCTAT<br>TGCGAAACAATCGCTATGCGGCTCAGGAGGCGGAGCATCAGCAGCCCTCAACGCCCACTCT[G/T]GAAAATGACTGCCCCGTGGACATGGAGAAG<br>CTTTACAGCCTGCCATATTGGGGCGGCGATGGACGCAATCATGTGCTGCTCAAC |
| FBgn0034483sechsim3 | 2R | 16 | 2R_16 | CGCCGAGTACAATCTGGTGAAGGAGGAGCTGAAGAACATCTATGCCATTCCCAGTTACGCATGTGGATTGCACTGGTTTGGCGTGATCTTTGTGCACA<br>GTGGAATCTATGCGGGCAGCGTGTTTCGCTTCTCCATTCTGCTGCCGGATAATTTTCCGGYGGACATCAGCCTGCCCACGGTGGTCTTCTCCWCGG<br>CAGTCCTACATCCGCACATCTGTCCGCAGAA[C/T]AAGACCCTCGATCTGGCRCATTTCCTGAACGAGTGGCGCAAGGATGAGCACCACA |
| FBgn0020312sechsim1 | 2R | 17 | 2R_17 | GTCTTCCTGAACGACTTCTGGGGGACACCGATGCGCAAGGAGCAGTCCCACAAGTCTTATCGCCCGCTCACCGTGCTGACGTTCCGCTTCAACTA<br>CTTGCTGCATGCACTTGAGCCGTTTGGATACCACCTCGTCAACCTGCTGCTGCACTTGTCGGTCTGCCT[C/T]TTGTGGCGTCGGGTCTGCCGGTTG<br>CTGCTGCGTCARTGTGCCGCGTCGGGCAGTAATGCCATCTCGGCCTCATCCTCGTCATCCGTCTCCCAGCTCAACACATGCGCTTTCGTGGCCTC<br>GCTGCTCTTYGCCGTGCATCCAGTTCACACGGAGGCCGTTACTGGCGTCGTCGGGCGTGCTGAATTGCTCTCTTCCATTTGCTTCCTGGCCGCCTT<br>TCTTAGCTACGCGAAGTCCGTCGGCGACTCGGGTTGTCCGCGACGCACCAATTGGCTGACGCTTTTTGGCTGCTTTGGCAGTTGCCTGTTGGC |
| FBgn0029147sechsim1 | 2R | 18 | 2R_18 | ATGACCCAGCGGAATCGGCACAACTGTCGACCAGATTCAACGTGAGCTTCGCCCAGCTGGATTCCTGCGAAACTTACTCGGTATCGAATACGGGCT<br>ATGGCTACCG[G/T]AGGTTCTACATGCTGCGCGAGCTGAGCAAYAACAACAGGAAGGCCGGAGAGCGACTGCACTTGAAATTCTACGTCCTGACCG<br>CCATGGATGCGCACATCCTGCTCTCGGTGACGAATCACCCACGCCCAAATGATCGCGTTTACGAGATCG |
| FBgn0034834sechsim1 | 2R | 19 | 2R_19 | ATGAGACGTTTTCGGCCCACGGATCGTTCCTGGAATGCTCTGCGTTCGCGTTCACGTTCGCGTTCGCCATTCCG[G/T]TCGCAATCCATTTCTCGATC<br>CCGGCCCTCCCGTTCGGATACCATTTTCACAGAYAGGCCGCGWCTGCAGCGACGTTTCGGCCGGCCCGCCACCTCCTGGGACATTYCACCGG<br>AGGGTTATAGCCACCTGACTCCGCAGCAGTAYAAGGCCATGCAGGCCAGCGGGCAGATTGCCTCCAGGATTCTTTCGGATGGCATACACGCGGG<br>GGAGTCCGCCGCGATTTCGATGATCACTCGACAAGCTCGCCGCCTGTATGTGGGTAATATTCCATTCGGAGTTACGGAGGAGGAGATGATGCAGTTC<br>TTCAATCATCGGATCA |
| FBgn0023081dsimech2 | 2R | 19.9 | 2R_19.9 | GGAGACAACTATGCCACTCAGCATTTCCTCCGAGATGTCGTCGTACGAGATCGAACGCCTTGAACTCCAGTTCTCGGARAAGCTAAGTCATCAGCAG<br>ACGCGCCACAATATGGAGYTGGAAGCACTGCGCGAGCAGTTCAGCGAGCTAGAGAACGCGAATTTGGCACT[G/C]ACCAAGGAGTTGCAACAAACA<br>CAGGAAAGGCTGAAGTACACTCAAATGGAGTCGATTACCGATTCCGCGGAAACGCTTTTGGAATTGAAGAAGCAGCATGACTTGGAGAAGAGCTCGTG<br>GTTCGAGGAGAAGCAGAGGTTAAGCTCCGAGGTGAATCTCAAGTCGAAGAGTCTAAAGGAACTTCAAGCGGAGGACGACGAGATCTTCAAGGAGTTG<br>CGAATGAAACGGGAGGCCATCACCTTGTGGGAGCGGCAAATGGCGGAGATTATTCAATGGGTGTCYGATGAGAAAGACGCGCGTGGCTACTTGCAG<br>GCCTTGGCCACCAAAATGACGGAGGAGTTGGAGTACCTGAAGCATGTGGGTACCTTCAACAACAACGGTGTGGACAATAAGAACTGGAGGAACCGG<br>CGCTCCCAGAAGCTGGACAAGATGGAGCTGCTCAACCTGCAGAGCGCGCTGCAAAGGGAGATTCAGGCCAAGAACATGATATCGGATGAGCTGAG<br>TCAAACCAGATCGGACCTCATATCCACACAGAAAGAGGTTCGGGACTACAAGAA |

| | | | | |
|---|---|---|---|---|
| FBgn0001148sechsim1 | 2R | 20.9 | 2R_20.9 | CAGTTGGGCGGCGTCTTCATCAACGGTCGTCCGTTGCCCAATCACATCCGTCGCCAAATCGTGGAGATGGCAGCCGCTGGGGTTCGTCCCTGTGTCATCTCCCGCCAGCTGCGCGTCTCTCATGGCTGCGTCTCAAAGATTCTTAACCGCTTCCAGGAGACTGGCTCCATTCGGCCCGGAGTGATCGGTGGCAGCAAGCCCCGAGTAGCCAC[A/G]CCAGACATTGAGTCCAGAATCGAGGAACTTAAACAGTCGCAGCCCGGTATTTTTAGTTGGGAAATCCGCGCMAAGCTAATYGAAGCTGGAGTCTGCGACAAGCAAAATGCTCCGTCGGTGAGCTCTATTTCGCGTCTTCTGCGAGGATCCTCYGGATCGGGGACCTCCC |
| FBgn0020386dsimech2 | 3L | 0.1 | 3L_0.1 | CCAAATCGAACCTATTACCTGGATGATCCCGAAGGCTATGCTATCCACTGGTCGGAAGCCATTGAGAGCATGCGCAAGTTGACCTACGGAGATCCCTCCTCCACGTCTGCAGTGTCTTGCTCCAGCGGCAGCAGTAATAGCCTGGCTGTCATCTCAAATTCATTCGCCGCCTCCTCAAGCAAYTCGCCCACGGTGAGACGCAGTTCACC[A/T]GTAAACGCTCCTCAAGCTTCGACGGCGTCCGACAACCGGACGTTGGGTAGCACCAGAACGGGGACGTCACCGAGCAAGAAGACGGCGTCCAAG |
| FBgn0035179sechsim1 | 3L | 0.9 | 3L_0.9 | GGCCTCCAATCGCGTGCAGCGGYTGAAGCACGCTCTCAATATTATGGAGTACATTTTCCGACTGGACGAAATGAAGCCGGAGCCGYTGAAGATGTGCAAGATGCAGGTGAATCGCTTCGGAACTGCCTCTGAGCGAAGGCTGAAGTACCAGAGTGCCGAGCACGTCTATGTGGAGAGCCAGCTGGACAAGCGATTGTCCAGGGGCGTCAAGTGCCACGCCCACCAGGACTGCCACTTTCACTCCTGCCGAGGATTGTGCGACGAGGAGAAGCAATCCTGCACGCACATCCAGCAGAACAACAACTTCCA[G/A]ATTTTCTGCGAGCACATCCTCCTGGGCGGAGGCACCTTTCAGCCCGGCCTCCTCAGCGGAGTGCGCCTCTCCAAGGCGCTCCAAAAGCTGGTCAAGATGTGCGTCCAGCCGCCCAAGGAGCATCAAGTGCCAGG |
| FBgn0035294sechsim1 | 3L | 01.9 | 3L_01.9 | CGTCGAACGCGAGAAGCTGGACATTGATCGCATGCGCAACATGACCGAAGAGGAGCGACGCCAGGAGCTGCGCCAGAACCCCAAGGTGGTCACCAACAAGGCCACCAAGGGCAAGTACAAGTTCCTGCAGAAGTACTATCATCGTGGTGCCTTCTACCTGGATGAGGAGAACGACGTCCTCAAGCGCGACTTCGCCCAGGCCACGCTGGAGGATCACTTCGACAAGACCATCCTGCCCAAGGTGATGCAGGTGAAGAACTTCGGCCGCTGCGG[C/G]CGCACCAAGTACACGCATTTAGTGGACCAGGACACCACCAAGTTCGACTCGCCCTGGTACGCCGAGTCATCCAGCAACATCAAGTTCCACAACGAACACGCCGGCGGCATGAGGCAGCAGTTCGACAAGCCCACCGGCTCGAAGCGAAAAAAGATGGAG |
| FBgn0004372sechsim5 | 3L | 03 | 3L_03 | TTGGTACGCCGGAATATGGGGAAGGCGCGTCGATTTTCGCCCGCCTTCATCGARCTGGAACGCGAAGAATTGGAGTGCCARCGCCGCATTGTGCGCCAGTTGCAGCAGCATAAGTTCAATCCCAAGGAGAACGTGGGCTACTTGGACCAGATACCCAAGCGTGTGCCCCTGCCACTGKCCAAGGATGCCAGGGTCAGCAGTTTTCTGCACGGACRCTCCTTCGAGGGCATCGTCAATGGCATTGTCATGGGCTACGATCC[G/T]CAGGACTACACCTATCTGGTTCGATTCAGTAGAAATGACAATGCAGTCGTGCTCTGTCTTTCGGACTCACAGCTCTATTCCGACGAGGAAACCGC |
| FBgn0024179sechsim2 | 3L | 04 | 3L_04 | GGAATAAATATCATAGCCACAACGACAACGGCGGCTGCAGTCCATCACCAGATGAGCTCCGACACCACGGGACTTATACAGCCACCTCCCAATCAGCAGATACCTCTAGCTGCCTTGGAACGGGAGAAGAATCATCTGAGCTACCCGCAACAGCAATTGCAACCKTATCAGGGCAGGAATCCCTGCCAGGAGCGCAATCTGGCACCTCTGACAATGCGAACTCCGCCGGTTCTTGTGGAGCGGAGCAAGAAGCACAGTTTCCAGAC[A/G]CAGCCTCAGGAGAACTCACTKTCTTGCTTGGAGCATGATGTGAGTGTGGAGGAGCTGATAGCCAGCCATCATCACCAGCAGCAAAAGAACACGATTGTCAGCACAGGTGGAAATGGCAATTCCWGCTTGGGTCAGGGATTTCCCAAGCAGCAGAACACGGATCAAAAACTCCGGGGATGGCATGGAGTAAGAGCCCTGATCCACAAGAAACTTTTCCGCAAGGAGCACGCCGAGGAGCTGTGTCGCCAGTTGCAGTTGGG |
| FBgn0052232dsimech1 | 3L | 04.9 | 3L_04.9 | GGGTCATTCGATCAACGAGTTGCTGAGGAACTTCCTCACCTGTGCTGACTACACTCCCATTACTGAAATCMTTAGCCAGGAGCTTCAAAGGTTACTGAATCGAAGGACTAGATTGGTTAACTCCTATATCTCTGGCCTGTTGCAGGAGTTTCAAGGATACAAAGTAGTAAGACACTTAAGATCAGT[T/G]TTTCTACTCTGGAGCTATGACCACTTTCAAAGTGAGTTTGAGTTGTTCTTYGAGTTTCTCAGAAGGAATCAACTGGTAGAGGCCACTGACAAATTGAAGCACATCGTGGTATTCCAAGATTACACCCTGGGTCATATGTTTCAGGTTCGCTTGACGGAAGCYCATCCTGAATTTATCTTCCTCAGGGTCAGGAATGATTCCCTATTGAAC |
| FBgn0052232dsimech1 | 3L | 04.9 | 3L_04.9 | GTCGTCGTGGCAAACGCGACTTGCTGCGGCCCACCCAGCTCATCATCACGCGGGCCAAACCCTTTGAGGCCATCTGGGCCACCAACTACTGCACATTCATGCGCGAGTCACAGACGGAGGTCATCTGGGCCACAGGGCTTAACAACTTCAAGCAGCTGGCCCACGAGACAAAGGGCAAGGAGTTCGCACTGACGCCCATCAAGACGGAACTTAAAGATATCCGACATATTGCCGGCGGTCAGCACCATACTGTGATCCTGACAACAGACCTTAAGTGTTCCGTGGTGGGTCGTCCGGAGTACGGTCGGCTTGGACTGGGCGATGTCAAGGATGTTGTGGAGAAGCCAACGATTGT[G/A]AAGAAGCTGACCGAAAAGATTGTGTCCGTTGGATGTGGCGAAGTGTGTTCCTATGCCGTTACAATTGATGGCAAACTGTATTCCTGGGGCTCCGGCGTCAACAATCAACTGGGCGTTGGCGACGGCGATGATGAGCTCGAACCGATCGTGGTTGTCAGCAAGAACACGCAGGGCAAGCACATGCTGCTTGCATCTGGCGGTGGTCAGCACGCAATCTTTTTGGTCAAGGCGGAC |

| | | | | |
|---|---|---|---|---|
| FBgn0002638sechsim1 | 3L | 05.9 | 3L_05.9 | ATGTCCAGTGCCCAAGTGTCGCAACAGCCGCCTCCACCGCCGCTGACCGCCAGCAGTAGCACCTCAATGCCGTCGCAATTGCTACTTAGCCACATTCAGCACCACAGCAGCAGTCTGCACCATAAGCGCAGTCATCAGCGATCTTCGCCACCTGCCCAGGACTACGATCAGGGCAGCAATCAATCCACACTCTCGCCCAGATCCCAGATAACACCCAGTCCGCCAGCAGCCAGTCCCACGCATTCGGCGGCCACCACAGGATCACCAGTTAGCACCTATACGCCGGCCTTGAGTCCCGGTGGTGCACAGGATAGCAGCCAAACGGAGAGTGGTCAGAGTTCACAGATGCCTCTGCTAGTGGCTCCCATGCCCGTGGGCACTGGTCCGCCACATTCACATCCACCGCCGCCGCATCCTCGTCTCTATGTGGAGGGTTTCCCGCCACCGCATCACGTGCCACATCCACATCCCGCCCTGGGCGCCTATCCCCTGCCACGACTGCCTCTTATGCTACCGGCAAATCCACCAGGTGTGCCATCTGGAGCAGCTGCACCACCGCCCGTCTCACCGCAGCCCGCTGCCAATCACCTGAGCCACAGTGGAGCCACCATGGCCACAACCACACTCCTGCCACCCGCCCAAAGTCAGCCCAA[A/G]AAATCCTTCTGCATCGATGCCCTGCTGGCCAAGAGCCAACAYCAGTCTGGTGAGCCACAGCCYATCATTGTGGATGATCGTCTGGCCGCCCTGCACTATGCCCGSGATCAGGCGGAACTYAATCATGCCTTCGTGACCGCYTCCAATGCGGCGGCAGCTGCTCAGGCGGCTGCTTCGGCCGCCGGCGGCATCAGCGAGCAGGAGGCACTGCAGCGCATACGGGACTCCAGGGAGTACGACTCACCWAGTCCCGACGGCATGTCAAGATCGGAGTCGCCCACTTCGTCGCACCGCTCCAGTCCGCCYATCAGTCCCGGTTGTGAGGATCAGCAGCAGCCCCACGGTGCACTGCATCCCCAGCATCTGTCCATGCGCATGTCCGACTTCCACGACGAGTTCAAGAAGCCAATWCCACCMCACAGTCCCATCAGACCMCAGGATTTCCCGCTCTACGCGGGCGGACATCCCTACCAGCTGCTGGCCCAGGGTGGATCMGCCTTCCACAGACCWYTGGATCCCTCAGGGAAACCCATTCCGATACCCATGGGTCATAATTTCATGCCGTCRCAGCTGCAATTYGAGTTCCTGGCCCGCGCCGGAATGYTGCACCATCGAATTCCCGAACTGGCCGCCTATCCGCATCATGCGATTCTGGGCAAGACCAGGAGACCACGAACCGCCTTTACATCGCAGCAGCTCCTCGARCTGGARAAGCAATTCAAGCAGAACAAGTATCTGAGTCGGCCCAARCGATTCGAGGTGGCCAGCGGTCTGATGCTGTCCGAGACGCAGGTGAAGATCTGGTTCCAGAATCGTCGGATGAAGTGGAAGCGCTCCAAGAAGGCGCAGCAGGAGGCCAAGGAGCGGGCAAAGGCCAACCAGCAGCAGCAGCARCAGCAGACGCCCAGTGCCGCCAGT |
| FBgn0041156dsechsim2 | 3L | 07.9 | 3L_07.9 | TATGGTATTTGCATACGCAAGGCAGCGGGAATGTGTTCCATCACCTACAGCCAGGTTGGCTCCGATACGTACTCCTTTACGCTGACCAACGATGTGGGTGCAGTGGATCCCACTCTGCTGGCCACCTCATCTGTTCAGAGTCAGGAATGCACAACGGACTACATCAT[C/A]ATTCCSTCGCCAACGCAGGGCGGCGTTGCGATGCCCAGCGATCGCTTTTGTGGATTGGGTCTGGTATCCACGACCACATCGGCGAAGCCATTCGTTGTCTATACAGTCACCGATGGYAATGAGGATATGGACATATCGAATCGCGGCTTCTACTTGTCGTATTCGCAGAACGCATGTCCAATTCTG |
| FBgn0035941sechsim1 | 3L | 08.9 | 3L_08.9 | CACATGCGGACCCACACGCGGTACGATTCCAGGGGAGGATCTCGAGAGGGCAGGGAAGGACGCGAGGGACGTGAGGGCAAGGAGAGCCGCGATTCGCGCAGGAGCAACGAACGACTCGAGGAGCCCAACTC[A/G]CCCGGTTCCCATTCGCTGCTGGAYATGAAGATGAACATGAACATGGGCATGGGAATGGGCCTCAATATGGGCCTGCCCATGAATCCGATGAGTCTGCTGCAGGAGGAGCTGCTGCAGAAGTCCCAGCCGGGCCTGACCCTGGGYGGACCCATGCCACTGGTGGTCAAGACGGAGAGCGCC |
| FBgn0013469sechsim1 | 3L | 10.9 | 3L_10.9 | ATGCAGGACCCAAACGAAGATACCGAGTGGAATGATGTGCTCCGTGCCAAGGGAATCATTGGGCCCAAGGCAAAGGAGGCGGAGATCACAGAGGATCAGATCCAGAAGCTGATGGACGATGCCATCCAGCGGCGCACAGATCTGCCACTGAATGAAGGCCAGCGGGACAAGAAGATCGACGACATGTCGCTGGACGAACTCGACGAACTGGAGGACTCCGAGGACGAGGCTGTTCTAGAGCAGTATCGCCAGCGACGCATCGCSGAGATGAGGGCCACMGCCGAAAAGGCGCGATTTGGRTCCGTGCGCGAGAT[A/C]TCAGGGCAGGATTATGTGAACGAGGTGACCAAGGCCGGCGAGGGMATCTGGGTGGTACTCCACC |
| FBgn0036237sechsim2 | 3L | 11.9 | 3L_11.9 | CGGCTGTAGCGGGCTATGCTGTCATGTATGTGACCATGAGGAAATCTTTGTTTAAACTGGAGGACCAACAACTTGAGCTGGCAGCTGAGCGGGAGAACACTAGACGGCGATCGAGTGCGGATATTACGGCTCGGGCAGAAGAAATGCGCAGACAACAGCCGACGGGACGTCAGTTACTGAARCCGGCCGAAAACTATTTTCAGACCCTGGCCCATTTGGAGGCACATCATCGCAT[C/T]CGGGACAAGACGCTGCTGATCAAGTGCCTCTTCACGCTGTGYTTTGTGATCATCTTTTTCCTGTTGCAYTCGCTGCCCTTTATGCACGGTGCCACATTGGGTTGGGTGGCAATTTTGGCCGCCTTCCTGCTC |
| FBgn0036329sechsim2 | 3L | 12.9 | 3L_12.9 | GTGCACCGACAGGGACTACGAAGCCGCCGCAGCCTTCCCAATCTCCGTCAGCAAAACAACCGCCTCAGCGAAATTGTCCTGCACAATCTTCGCCAGCTGGTCGCACGCCTTAACCAGGAAAGCATCGCCCAGGGACAGGTGATCGAACAGCAACAGCAGTGGATCAACAATCCCCGCCTGACGCAATCCGAGCGRTATGCCCTCCGCTTGAACCAAATCCGAATCGATTCCCAGCGCAGCCGCCAAGTCCTCGCCCAGATCGGCCAGATTGAGCAGCGTTTGCAGGAGGTTATTGGTCAGGTGCTGAGCCAGGTGAATGTC[A/G]ACTCGCTGCGTGGCCAGATGATCGATCAACGCCAAGTGGAATCCCTGATCGCCGATGTACTGTTGGGTCGYCTTGGTCAGGTGGGCATCATGACCATTATTCGCCAGGTGGTCCAGGACAACAATGGTGAGCAAATCGATCGTACTGGCTTGGGCATCCGCCTCAGCGA |

| | | | | |
|---|---|---|---|---|
| FBgn0000639sechsim2 | 3L | 14 | 3L_14 | GTGCACCGACAGGGACTACGAAGCCGCCGCAGCCTTCCCAATCTCCGTCAGCAAAACAACCGCCTCAGCGAAATTGTCCTGCACAATCTTCGCCAGCTGGTCGCACGCCTTAACCAGGAAAGCATCGCCCAGGGACAGGTGATCGAACAGCAACAGCAGTGGATCAACAATCCCCGCCTGACGCAATCCGAGCGRTATGCCCTCCGCTTGAACCAAATCCGAATCGATTCCCAGCGCAGCCGCCAAGTCCTCGCCCAGATCGGCCAGATTGAGCAGCGTTTGCAGGAGGTTATTGGTCAGGTGCTGAGCCAGGTGAATGTC[A/G]ACTCGCTGCGTGGCCAGATGATCGATCAACGCCAAGTGGAATCCCTGATCGCCGATGTACTGTTGGGTCGYCTTGGTCAGGTGGGCATCATGACCATTATTCGCCAGGTGGTCCAGGACAACAATGGTGAGCAAATCGATCGTACTGGCTTGGGCATCCGCCTCAGCGA |
| FBgn0000639sechsim2 | 3L | 14 | 3L_14 | ATGTCGGCCATCAGCAAGCAGAAGACCCTCAACACCATGTACAAGTCGAAAGTGTCGCCTTCKCGTCGCCTGAAGAATCTGYTGAAGCCRCTGCTCGGTCAATTCTTCAACATGAAGAAGGAGCAGCCCAAGAGGAGCTCCTACATYGCCTCCGAGGAGAAGGAGGCCGAGTTCGAGTGGCTCAGCAACGATATGGACAACATGGCCAACGAGCATTTGGAGCATCGCCTCATCGAGGAAAT[A/C]CGCCAGTGCGCCAAGGAAGATGCCGTCATTGTTTACAATGAGAAYGGATCCGGKGAGCTGCACACCATCGACCAGGAGGATTACTATGTACCAGTGCATTTTGCCCGCACCAATGCCGGCACCTTCTTCTGGACATCGGTGCAACCCAAGGCCTGCGAACCCTATGCCATCGAATGGAACTTCCTGGACCGTTGGGCTCAAGCT |
| FBgn0040296sechsim3 | 3L | 14.9 | 3L_14.9 | GGGCTGGGAACTGCATCGACGAGACGAAGTACCAGCGGGTGATCAAGGAGGCCTGCCTGCGCGCCACAGAGATCCTACGCAACGGCGGATCCGCCGTCGATGCCTGCGAGGCGGCCATTGTGCGGCTGGAGAACTGCGGCTACACAAACGCCGGCTATGG[C/G]TCAAATCTCTGCATGGACGGCTCTGTGCAGTGCGATGCGGCTATAATGGATGGCTCCACGCTTAACTTTGGCGCCTGCACGAACGTTAGTCGGGTGAAGAAYCCCATTCAGTTGGCRAGACGCATCTGCGATGCCCAGTCCAGTCCGCAGCTTCTGGAGCGCATTCCACCGATGATCCTCGCCGGCACTGGAGCGGAACACTATGCCGACGAGGTGGGTTGCTCCATGGTGGAGCCCGGCGTACTGATCTCATCTAAGGCCAAGTTCCAGTTTAATCACTACAAAAGCAAGTACGACCTCGTGGTCAAC |
| FBgn0036564sechsim1 | 3L | 16 | 3L_16 | AAAGCAATTGAAACGCTCAACGCCTAAGGATCGCCTGGCGGATGAGCAACTGGAGCCGTATACACAAACC[C/T]TGCTCTACCAGGAACATGGACCATTCCAAGTCCGGCAGTCAGCCCTTCTTCTAAACTGCCTTCAGGAGTCGAATCAAAGAAGGACAGTCGAACGCAGCTGGAAACAATGCGAGGAGTGTGTCAAGCTGCTTGACACCTCCGAGGAGGAGTTATCGCTGCGATCCCGMCTCTCCTACGGTTTCGCCGCCCACTTGAGTCCCAAGTGGCAGGTGCAGYTGCAGCTTGTGGACTTGCTGCGATCCTTGGGTATGACCAAAACAGCTCTGGACATCTGTCTTCGCATACACGCCTGGCAGCAGGTGATCGAGTGCTACACGAGCCTGGAACTTCGTCACAAGGCAGCGGAGATCATCAGGCAGGAGTT |
| FBgn0036772sechsim3 | 3L | 17.9 | 3L_17.9 | CATCAGACGCCCAGCGTGGAGCCGCCCTCCTATACGGATCATGCGGCCGGCGGAGGACCAGCTGGTGGCCAGAGCCTTGGWACGGGTCCAGGTAGTGTGCCCACCCAGCAACCGGCTACGCCCACAGCMGCAACGTCAGCTGGTGGCGCCGGATCTKGCGGACCTTCAAATGCAATTGGAGGGAATGCGGTTGGCACTATAAGTGTCAA[A/G]AAGCTTGAGATGCAGCARCAGACTCCATCGGCAGCTATGGCCATCAAGCAGGAGCCTGGTGCTCAAGGTCGGGGSGTCGGAGGTGTTACCTCCACCACGGAGGCCTTGAACAACTTTAAGCGACTGTACAACCCGCCAAAGCTGACGTTAAAGGATCCGGATAGCTTCTACGACGGGGAGTGGCTCAAGGAGGTCATCTATGACTTTCAGTACCAGGAATACTG |
| FBgn0003415sechsim3 | 3L | 20.9 | 3L_20.9 | TTCTCCCGATGAAAATCTTCTGCTGGCCGTGACTGCGTGCCGTATGTCCAACTATTTGCTGGCTCAGGAGGTAGTTTCAAAATCCACGAGACGCAACCTAGCCACAGCTTTAATACGGCACAAAAACGTTYTGTTCGACGACAGTACTGTGAT[C/T]TATCAGCGACGCAATCAACAGCAAGCTGACTCCCAGGCATTAAGTTTGCGAGCTAGAAAAAGTCGACGGATGGCGGCGAAGAGCAAACCAGCTCCAGTCGGAACGCCAGCTCCGAGTACCATAGTATCCTTCTCGGACTTTTGCGAGACGATTCTGTTGGCCAACGAGCAGCCAGCACTGCTTGAAGCTATCAGTGATGCTTTTATTCATGCGCTTTGCATCGAGCACAGCATG |
| FBgn0037142sechsim2 | 3L | 21.9 | 3L_21.9 | ATGGGCCGACGACCAGCTAGATGCTACCGCTATTGCAAAAACAAGCCGTACCCTAAATCTCGGTTTTGTCGTGGTGTSCCTGACCCCAAAATTCGTATATTTGATTTGGGAAGGAAAAAAGCTACAGTTGAGGATTTTCCCCTGTGCGTTCATTTGGTTTCTGATGAATACGAACAGCTGAGTAG[T/C]GAAGCTYTAGAAGCTGGACGCATTTGTTGCAACAAGTACTTGGTAAAGTACTGCGGTAAGGACCAGTTCCACATCAGAATGCGCCTGCATCCATTCCACGTCATTCGCATCAACAAAATGTTGTCGTGCGCTGGAGCTGATAGGCTTCAAACTGGAATGCGAGGAGCGTTTGGAAAACCTCAGGGCACGGTTGCTCGAGTTCGTATTGGTCAACCCATTATGTCWGTCCGCTCTAGCGATCGTTACAAGGCTCAAGTTATTGAAGCTTTGCGGCGTGCTAAGTTTAAGTTCCCTGGACGTCAGAAGATCTACGTTTCAAAGAAGTGGGGATTCACCAAATACGAACGTGAGCGTTATGAAGAACTTCGCGATGACAACCGCCT |
| FBgn0024733dsimech1 | 3L | 23 | 3L_23 | ATCGTCTCCAAGCGCTTCTTCATCCACTCCGCCCCGGAGGAGGCTGAGGATTACAAGGAACGCCACATCACCGTSGGCGTGCCCAAGCGCAACTACAACGTGGTGTTCATCAAGTCGCCGCAGCGCAACAACAGGAAGACCATCAAGATCAGCCCCGCCGCCAACGAGGAGAAGACCGTCATCTACGTGCTGAGCAA[A/G]AAGGGCGAGAGCGACTTGAACGCCGAGGTAGTGGAGCAGGCCAGCTCCACCAGCAAGCCGGAGGTCTTCTTCATCAAGTACAAGACCAACGAAGAGGCCGCCCACGCCCAGCAGCAGATCCAAGCCCAGTACGACGCTCTGGGCGGCAGTAGCCAGCTGACCGACGAGGGAGTGGCCCCCGTCACCTCCGTGATTGGAGCGCTGGGCGGCAGTGACGGTCACATCGACGCCGGATCCGCTGTGGGCGCCACAGGAGCCGGTCAGATCGTCTCCACCGGCGCTGTCGGATCGCACACGGGCAATGCATACCTGCCACCCAATTTC |

| | | | | |
|---|---|---|---|---|
| FBgn0037227dsimech2 | 3R | 0.1 | 3R_0.1 | ATGCACGGACGCGTAAAAGTGAGGACSACGGAGGAGGAGCGCGAGCGGAAGAAGAAGGARCAGGCTCTGAAGGTSCGGGCCTACCGTGCAGCC<br>ATGGGAAGRATTCAGAAGAAGAGRGAGGCGGGCGAGCTGGACAACGAAATGCTSAGCCTGACGGTTCAGATCTTGCTGCGGAATCCGGATGTCAG<br>CACGCTGTGGAACATWCGCAGGGAGTGCGTGCTGGAGAAGCTGTCCAGGCTAAAGGAAGAGGAGACCRCGTGCGAGACGCCCAGCGAGGAGA<br>AGCCCGAGGAGGAAA[A/T]CCAAACCGKGGARGATAAGMCGRGTGAAAAGAAGGCCGCTGCCGAGGACAAGKYCCATTCCATTTTCACCTGCGAA<br>CTGGACCTCACCGAGCAGTGCCTTATGGTCAACCCGAAGTCCTACAACGCCTGGCACCACCGCTGCTGGACCCTRGAACAGAATCCCAGGGCC<br>GACTGGCAGCGGGAGGTGCAACTGTGCAACAAGTACCTCAAATTCGACGAGCGCAACTTCCACACCTGGGACTACAGGCGCTATGTRACCGGCAA<br>GGCCATGGTGCCGGCYGCCCAGGAGCTGGACTTYTGCACMGAGAAGATCARGGTTAACTTCTCCAACTACTCCAGCTGGCACCACCGYAGTCTYC<br>TGCTTCCGGAGCTCTACCCCAACCAGCAGAGAGACCGGCCCATRAGTGAGGAGAAGTTGCAGAAGGAGCTTGAAATGGTGCTGACGGCTGCTTTC<br>ACGGATCCCAACGACAGCAGTGCTTGGTTCTACCAGCGCTGGTTRCTGGGCAGCGGCGCCCAATTGGATCGGGCTCCTAGAATAGCTGCTTTTCG<br>ACTRGAGTCACATGGAGCAGTCCTCGCCCTMGACAARCCCTACCCAGATATCAMACAAYTGAAGACTGAGYTAGTTGGGGGAGGACAGACGATTTTA<br>CTGGAGTCCTGGCWACCAGTGGACAAGCGTGGAACCAGTTGGAAATGCCAGCAGAACTTCGATTACCAAAGAAACTCARCTTACTCCYTTAGKCTCT<br>CYGAWCAGAAGGTTGACCTCTTGCCTCTGTCTGGCYCGGAAAACGGAGCCTATTACTTTGCGCCGCCACATGGCACYGCGAGTTGCAGCAAGGAT<br>CTCCTGGCGGAGCTGCAAACGCAGCTGCAATCCTGCCTAGATCTTCTGGAGTTTGAGCCGGACAGCAAGTGGACCCTACTMACAAGTGCCCTTCTT<br>ATGCGTGCTATCGATTTCTCAGCCAATCACGAAACGAGCTTGGCGCACTTGGCCAAGTTGGAAAAGGTGGATTCCTTGAGAGAGGGTTACTACAARGA<br>CCTCGCTGCTAGGTGGACCTTGGAGTGGGAGCTTGCCAAGTGGCCTCWATCTGCCGACTTTCCCAAGAGATTCGACAWAGCGGATGACGTACTG<br>CTGGAATCGCTTCCCTATGATCAATACTTAGTCATCGCTGATGAAATAACCTTGCCCAASAAACTACGARACAAATTGGGGRAATCAGYGCCCAAAACGT<br>TTGYAGACCTGAAGTTGCAGTCA |
| FBgn0037293dsechsim9 | 3R | 0.9 | 3R_0.9 | ATGGCCGGGAGAAGGGTGATTGGTCTCTGTTTTGGAATACTCTTACAAGTGATGCTGGCGACCGCTGGCCAGGATCGGATGAGCATGCTCTGTTCCG<br>ACGACGACACACCAGCCTGTGCCCAGAGCCGCGAAAATGGAATGTACTTTTTGTTCGCCAACGAGTGTGACCTTCGGAAGGCCCAGCGGGGCAA<br>CCTCATGAATGGCCCYCTATATGACGTTACCCTCCGCTTCTGCTTTCCCAGCTGCGAATTTGAGTGCAGCTCAAGGTATCAGCCAGTTTGTGGGATCA<br>GCTCRAAGTCGGGGGAACGRAAGACCTTCAGGAGTCGCTGCGAAATSCTGCGAACCGCTTGCATCTCCCGGTC[A/G]GACTGGATGGTCCATCAAT<br>GGGGAGTGTGYCCAAAAGCYAACGCGGTGCCCCAGAGCAGYGAAAAGCCGCCCGTGCCCGTGCCCTGCACCAGGATYTATCGACCTGTATGCG<br>CCATGTACGCGGGTGTCAAGTCAACCTTYTCCAACGAGTGTTTGGTGAACGCGGAAAACATCAAGACCCAAAGAAACTGGCGCATCGTTTCCGAGGG<br>TCTTTGTGGCGAGGACAGCACCAAGATGAAGCACAGTCGCAAGCAGAAGCCGAAGGCCAAGCCCAAGTCGGATGCGGAGCGCACCAAGCGGAG<br>CCACAGAGGCACGAGGCTGTCCACTCAGGCGGAAGACTTCCAGATACCGGAGGACGCCGTGGAGATCTACGCTCCCTCGACCTTTCACACGCA<br>GTTCATCAGCCAGTCTGGCGCCATGGAGAAGAGCTACTCGCTRCCCGCCAGGAAGCCCTATGTSGTGCTCCCCCCCAAAWCGAAGCGGCGAAT<br>CACCACAAAGTCTTGTGTATTCGGCAATCAGCCCGTTTGCGGGAACTTAAGAGGCCAGAGTCGCACCTTTTCCARCGTGTGTGAACTCATGGAGTTTA<br>GCCAAAAAGTCGGAAATGCATGGACCATTGCCCACGATGGAGCCTGTCGCCGCTGCGATAAGACGTGCCCTACGGTGTATCAACCTATTTGTGCCA<br>CCAGAAACGGCATCAATCACACAATCGTCAACGAGTGCTATTTGGAGCGAGTGCGCTGCAAGGATCCAAAAAGCATATGGAAGCTATCCCACARAGG<br>TGAGTGCGCCAAACCAGTGAGCAACCCACGGCACATTWMCTCCACTGGTAAAAGTYGCCCAACGTCGTTGGTGCCAAGTGTTTTATACGGAAAGAG<br>CACCACCACACCACGAAGCCGCTTCAGGAAACCAAGCACCACCACACCACGAAGCCGCTTCAGGAAACCAGTACGTAGGTTCACAACCACAAGG<br>ACTCCAACCACACCAYACAAGMCACGTCAGCTGGCGATGACAAATTTCACRGCCACTTCGCCTTCCCTGGAAGCGAATAATCGCAAAATACGCAAAA<br>TTGAGCTAGCCGCTGCGGGATTTGYGGGCTTGGGTGCGTCAAGCGGGTCAAATTACCTGAGCTCTGAGGAAGATGCYTTCTGGTCCTCTGATGACAG<br>TTGGCTGGTGCAGAAAACTCTCGAWAATGTCAAGGGCTTACTTGGCAATAAGCCTACTTCCTTTAAGAAGGCGCACGGTGRGAAGTACCCACTGTCC<br>YATTGGATTGTCCCACCAAGGCAAACAAGTACTACCAGCACCACCACAGCTGCACCACCAGCTAAACTACCAGCTGTTCTCAGCATGKCATCCACG<br>GAGCTATTGAATTTRGATTTTGGAAACCYTGCCGGCGCTTACGAAGAAGCTGATCACGAATCGATGCTCATRCCGACCACTAAAGAGGACACCACCTC<br>AACCTCAACCACGCCTGTGCCCAGCACCACTACTCCGCCCAACATCACCGAACCTTCAACCACTGAATTGCCCACSAGCGTTGACGTTGCCTCTG<br>ACTCGGCGGAAACGGTCAGTGAAGACTCCACCACTGATTTTGGAGAAACGACTKATCCAACGGGCTCAACGACYAGCAACGATTCCAGCACCTCAG<br>ATGCACCACAACCTTCGACCACAGAGGCGGAGGAGTTGAACAGCCAAACCACGGGGAGCGAGTCCACAACCACCGTGGCCTCCGACGAGCTGT<br>CCACGACCACCTTGAATGCGGACTACGCGGCGGATGAGTCATTTACCGAGTCCAGTGGTCAGACCTCCATCTATGGGCTCGACAAGAACTCCTTGA<br>TCATGAGGTTGCTACGTGCCAGAAGTAACCAAAACGTGCTCATC |

| | | | | |
|---|---|---|---|---|
| FBgn0037405dsechsim1 | 3R | 01.9 | 3R_01.9 | GGAAGTCCTGTTGGAATGCCTCTGAGCCAGGAGGCCATCGCGCAGCGGCTGGCGGCGCTTAGTCGTTGGCAGGACGAGCAGAAGCGGCTGCT<br>GCAGGAGCGRCAGTCCAATCAGCGCGTTTTGCTCGGCTTGGAGCAACGCAATATGTACAAAATGCTGGGACTGCTACACCAGGAGACGGAAAGCC<br>GCGAAAACAGTGTGTTGGAGGAGTGGGACGAGGAGCACTCTATACAAATGCCGCGACTTGCGGCGGATCCGGACGACCACGAGCCAGAGATTCC<br>AATGGCTGATCCCCAGCCGGCCAAGCCCAAACGYCCMTTTCTGCGTCGCGGCGAAGGCCT[A/T]AAGCAGCGGTTCAAGATCAATCCCGACCAG<br>CTGCGCCTGGAAAACCTGCCGCGATACAAGTTCGCTAATGCCCACCCTCAGTTCCGCACGCCTCAAACGAAAAAGGGTATT |
| FBgn0001020sechsim4 | 3R | 02.9 | 3R_02.9 | ATGCCTTCCTTCCAACCGATCGAGGCCCCCAAGTGTCCGCGCTGCGGCAAGAGTGTCTACGCTGCCGAGGAGCGTCTGGCTGGCGGCTATGTAT<br>TCCACAAGAACTGCTTCAAGTGCGGAATGTGCAACAAATCCCTGGACTCCACCAACTGCACAGAGCACGAGCGCGAGCTCTACTGCAAGACGTGC<br>CACGGTCGCAAGTTCGGGCCGAAAGGTTACGGCTTCGG[T/C]ACTGGAGCGGGCACCCTCTCCATGGACAACGGGTCACAGTTCCTGCGCGAGA<br>ACGGCGATGTGCCGTCCGTAAGGAATGGAGCCCGCCTGGAACCCAGGGCCATTGCTCGTGCCCCCGAAGGTGAGGGCTGCCCTCGTTGCGGT<br>GGCTATGTGTACGCCGCCGAACAGATGCTGGCCCGCGGACGCAGCTGGCACAAGGAGTGCTTCAAGTGCGGTACCTGCAAGAAGGGTCTGGAC<br>TCGATCCTGTGCTGCGAGGCTCCGGACAAGAACATCTACTGCAAGGGCTGCTATGCCAAGAAGTTTGGRCCCAAGGGCTATGGTTATGGCCAGGG<br>CGGTGGTGCTCTCCAGTCCGACTGCTATGC |
| FBgn0014863sechsim1 | 3R | 02.9 | 3R_02.9 | ATGGAGGAGGTTCTCGAGACGATGCAGCTGGAGAACATGCACTGCGATGTGTGCGAGGGTACGACCTTCCAGGAGCGCGAGGGATTCTACTACTG<br>CGTGGAGTGCGGCACACAGAAAGATCAGATTCGTGCAGTGGACATTACCGCCGAGGATAACTTTGATGATACGGCAGCAGGCCGGTACACAGCCA<br>GGACGATCCGACAGAAGAAGGATACGGAGAAGGAAGATGAGGACGACATCACCTCGTGGGAGTTTTACAATTACGTGCTGCGCGGCTTCCTTCAGG<br>AGCTGCTCAACATGGGCGCCAAGCCGGAGCTTAAGCTGATGACCCTGCAGGTTTGGACGGCTTACTTGGGCAGCATGGAGGTGGCCTTCAGCAAG<br>AGCAACAAGACGGGACTGCCCAAACTTAATGTGCGGGCGCTGCCCATAGATGCCCGCATCATCTACAACCATAAGCCATTGAAGAAAGGCAAGAAG<br>GGCAAGAAGAGCACTCTAACGGGTGACCCAAATGATGAGCGCGCCAAGTTCAGATTGTGGAACAGAACTAAGCGGAATCTAGACGCCTCCGGCTAC<br>CGCTCGCACGGAGGAGCTTCAGAGTCCGAAGGGGAGCAGAGTCTCCACCTGCAATGGTCAATGCGTGCACGCAAGTCCCTGAARCGCCATATGC<br>CGCTGAAGCACCTGGACAAGCACAGCCGCGACAGCAAGGGTAGTATGTCGTGCCACAGCYTKAGACCGAGGGTCAAGCAGCTGCAAAATTTTGAC<br>CGTAACATCTATTGCCTGAATATTATCAAGTTGTATGTGGTGCTGGGCATAGCCCTTAATATGGTGGAGGATGATATACAGCTATCRGACCTGCTKCGCTT<br>CATTGATGAGGAGCACTTGACCAAGCGGTGCATGCTTAACTATCTGCCSGGAAATGTGGCKGCCAAGGGCAAGGCACTGCTCAAAGATATGGAGCTC<br>AGCAAAATCAAAGACAARGTTACCAACAAGCTTCTGCGTTCAAACATTGCATSTATGTCTCGATTCATAAACTTGTCTGAATATCAAAAACCGAACCTG[A/C<br>]ACTCCCTAGCGGAGCGCTATATTCTGGAGTTGGCACTGCCACCTCGGCTACTGAAGTACGTGAATAGCCTGCTTGACCTGTATCCGCCCACTTTTTT<br>CAACGCSATGACCGTACACTCGTATCCGCGCTATGAAGCCAGGACTATGGCCTACATCCTCTACGCCATGAAGTTACTCTTCGGCTTGGACGATATC<br>AAGGAGCGAAAAATATCYGAATCGKCTGCTAAAATCAACGAACAACTGCTGGAAGTCGGCGGAGATGAGGCTCCCTTGCTTTTTGTTTTTACCRAGTGG<br>ATGGAGTTTGTGGAGATGAGGAAAGTGATTGTTTCGCATTACAACCAAAGTTTTGCCCGTCGYTTCGGGGTAGCCACCCGGACTGGTTGTCAGGTGGA<br>CGACATCCTAGCAAAAGAGTGGAAAGAAAAGGAGCAAGGCGAAACCTTCGACTGGATGCAGGGATCTGCGGCAATGAGGCGTCAGCATGAAAACCT<br>TACACACATTATTGAAACGATGCTGAAGGACCATTTTGGCGAGTCCAGCAAAGAAAGTWTGGAGAARGAGCACATTGAATTCCAGCCCAGCTTGACYC<br>CGGCTCACTCCTACTTCAACAGAATCCTACTCCAAGAATCGCGGTCGGATGGTGCAAAAATGAACATCAAAATTCCAGAAAAAATGAAAGTGGATCACA<br>CGGCCCGCAACCTGGATCCCTTTGTGCTGGAGACGACCGAGCTGACGCAGTATTTGTCCCAGCATGGCCTTAAACTACGGGTGGAGGAGCTKGC<br>CTGTCAGGAGGACATCCAAAAAGTGGGCATATTCCGSCCAATAACAATAATMCGAGGTGATGGTCGAGAATACCGMGCCAACACKGAAATTAAAACC<br>GAGACGTGGATCAACGAGCTWAAGAGGAAGGAGAAACGGCCGGACTTTCGGTTCACACAGCCCAYTGGAACMTATGGAGCCCGCTATTTGAAACG<br>TATAACAATGCGCGATGCAAGRCGTGTGAAACTTGAGATAAATAATCCSTTTTGGGATGTTACCGAAACGCCCAGCTTTCTTCTCAAACTCAACGATAACG<br>AGATACYACTTGACAGCCTTAGCTCTCTGCAAACCTTCGAGGAAGGGACCATGGATCCATTGAACATCCCTCTAGATCTGCCCCGTCGCAATCTGGA<br>AAAAAATGTGAATCCGGATGSTTCTGATAGAGCATCAGACCAAGTAGCGGGCGATATCAATGATGAGCCGCCAAGCCCTGAKACRCTGCTRTTGCAAG<br>TGTCCAATTTTGAYTGCTGGCTTCTACACGGGTATATGAAAAGAATTCGGAAGCATGACAAGCATGAACTGCGTCAGTTGTTCCCCTGTTCCTTCCGTTG<br>GCTCTTGGAGACCTGTGCCAGCACAATCGGCATAGTCTGGGAGGAGCTCTACGAGGAACTGCTCATCGTGGAAGTGATGTACCATCACAGCATCAG<br>GGACTGGAGCAACCATAGGAACTATCTGTGTATTCAACACAATACTCAAGAGAAGGACATCCGAGCCYTAGCCAGGACGTACAAGGAGTTATGG |
| FBgn0037792dsechsim3 | 3R | 05.9 | 3R_05.9 | ATGAAGATCTACAAGGATATCATCACCGGGGACGAGATGTTCGCCGACACCTACAAGATGAAGCTGGTGGATGATGTGATCTACGAGGTGTACGGCA<br>AGCTGATCACCCGCCAGGGTGACGATATCA[A/G]GCTAGAGGGCGCCAACGCATCCGCCGAGGAGGCCGACGAGGGCACCGATATCACCTCG<br>GAGTCTGGCGTGGATGTTGTGCTTAACCACCGACTGACGGAGTGCTTCGCCTTCGGSGACAAGAAGTCGTACACCCTCTACCTGAAGGACTACATG<br>AAGAAGGTGCTGGCCAAGCTGGAGGAGAAGTCGCCCGACCAGGTCGACATCTTCAAGACCAACATGAACAAGGCCATGAAGGACATCCTCGGCC<br>GCTTCAAGGAGCTGCAGTTCTTCACCGGCGAATCCATGGACTGCGACGGCATGGTGGCCTTGGTGGAATACCGCGAAATCAACGGCGACAGCGT<br>GCCCGTGCTCATGTTCTTCAAACACGGTCTGGAGGAGGAGAAATGC |

| | | | | |
|---|---|---|---|---|
| FBgn0037874dsechsim1 | 3R | 07 | 3R_07 | GGCGAGAAGTTCACTGCCAAGAGACCCTGCCACGTGGTCTCCTTTCCATTCTGGCCCGCCGGCGAGAGGCCCAACTACAGGGTGAACGAGGAG<br>AACCGCTGGTTCGCCACTTAYAGGTTCCAGCCGGTGAAGCGGATGCTCTTCAAGATGGTGGGCGACATCATGGCCGACTACTGGACAAAGATAGAT<br>GGTACTCAGCCGGCGGAGAGCGAGRCGATGCCCATGTGCGAGTTCCACAAGGAGGCACTGCGGCAGGAGCAGCTGCTGAAGGATGCAGCTGT<br>GGTCAAGGCCCACAAACAGTGTTTGGCTCTCGTGGATATCACTTCACGCCAGGATGCCCTCCTTAAAAAGCGTATYGTGGCCCAGTTGGAAAGGGAC<br>ATTGAGGAGGTCACGCWRCGTTGGAAACGACTTCGTGAGCGCCACCACACGGATGTCATGTT[A/G]TTGGAGGATCACGATCAGATGTGTTCGGTGG<br>GCAAAGTGCCYACGGCTGTGCAGGATATCAAGATAAAGTACCTATCCCAGGAAGCGGACATCGGGCGTCTGAGTGCCAARCCCACCTTGGGTATC<br>CATGTGATCACCGGCAGGGGTAACGTGAGTCTGCTAAGTAAAGTGCGGGTCTCGGATCCCGATGATGACATCCCATGCCCCACGGTACCCGTGGA<br>GCAAAAGCTRGCCAAATGTCCACCCCCCTTACTAGGTCACCTCAAACGTCCGCGRAAGGTGTCCATCACGAAATCSAAATTCGGCGACGATAAGCA<br>ATCTTCCTGCAAGAAATCWTCGAAGCCAGGACGCTTTTTCGAGGCTCCTCGCCAGCACAGYCCCTTCGTGTTCAAATACCATGAGGTTGCACCGCA<br>GGATCAGARTCCAGATCCCAAGGATGTGATRCGTGCCCAGGCCATCCGCTCYTTAAAGGAKCAAAGTTGCCCCTCAAGCGAGGATAACTACAACGG<br>ACGTATCAATCCCCGAGACCAGGTGGTGGCGGCCATCRTTGAGTGCGCCCAAAATATGTGGTTCGGCTCCCTGGAGGCTCATCAACGGCGCCTG<br>ACATCTGGCCAATCCGARAGACCAACGACTATTGAAGAGCCAACCAAGAATGAGCCGGAAGAAATCTGCAAAACAAATGAAGCCCTAGATTGGACAAA<br>GACAATCAAGAAGTTCGACCCCAACAACCAACACTTAATGGAGCGCCTGCTKCGGGATGGATTTTCCGTACTGAGACGGGATCCACGCTGCGTSTTC<br>GCCGCTTTTCCCAACTCGCAYAAATCCTTTGTGATGCTCGAGTGGATTAAGCGTCGATATGGGAAGACCTACTGCCACGAAGAGATTGTCGATACGATT<br>GAAAAGGGACTGGCCACCTTTATGCATGTTGAGAGCCARCTAAACGAAGCACCCAGCGCCAAGCGCGAAGGATTCTCGGCAAAGGACACATTCGA<br>CGACATTAAGCGTCTGGAGGCGCTGTGCAAGAAGTTAAAAACCGAATACMGAATGCCGCTCAACGAYAAGATCCTGACCCTGACAAGGATCTGCTG<br>GAGTGCCCTGAGTCCCCACTTGGCCAGAAGCTCATCCATGCTCAAAACCTTCTTCGCCTATCTGCCAGTTCGATACGCAGACATGCTKAAA |
| FBgn0037985dsechsim6 | 3R | 07.8 | 3R_07.8 | ATGTCAAACCTCCAGTTAACTCAGGAAAAACTAGATGCCCTACGCTCCGAATACAGGCCACGCCGGCCGTGCGCCCAGCTGAAATATCCTCGACC<br>CAAGACAAAGCCGGAGTACCARGCGATATATCCGTGGTTGCTCCCGGATGAGACGGATCCGCACTTCGTGTTCTGCGCGGTGTGCGAGTGCAGG<br>CTCAGCTGCAAGAGATCCGACCTGGGAAAGCACGAGGGTAGCATTAAGCACTCGGAAAATGCACAAAGAAAATCGTTGCCCGCCAAGGGCAACAAT<br>CCGAACT[A/T]TGATGGACCCAGCTCCGGTGCGATGAAGTGGGAKGATACCGAGGACGAGGAGGGTCTGCTGCAATACGGAACAGATCCCCAAGA<br>CGACGATACAGARGAGGAGGAKGGGGAACTGGACGAAACAGAGGGCGACTGCGAGGACGATGAGGAGCGGGCTSAAGAACTGGAAGCACAGAAT<br>GATCTCGAAACGGAGCCCGTTATTAAGCAGCAACGCCGCACCTCGGAGACCGACTCACAGAATTCCGGCATGCTGGATTATCTRCCACTACACGTG<br>ACCATCAACGAGTTGCCTCAAAGTCTTCCGTCCACTGTCTCCGCCTATCCCACATCCTCCACAACAGCAACAGTAGCTCCGCCWTGTAGGATCAC<br>CATTAAGAAAGTTCCAGTGCACAGTACTCCACTCAGTGGRATGTGCCAAGCTCAGGAGATCAATTCCAAGGCCACCATAACGCCTATACACACGACG<br>CCYTCCTACGCTGCCCGCCCGCTGCAATCTTATCAACTGCAACCATTGTCACCTATAACGCCAGCGCGCGACTCTTTGGAGCTGTTCTTCGACAGC<br>ATTTGCGCCACCGTCAAGAACTTGCCGCCCAAACTGGCCACCGAGGGCAAGATCCGGGTGATGCAGCTCATCGGGGAGCTCGAGTTGCGYGCT<br>CTTTGCGAACAGGAGGCGCTCTCTCAGGTCGCCGGACTTCCGGATCTCGCGGGAAACTCCATGTCCGGAGCATCTGAGGCCTCAGGCAGTGGTC<br>AGCAGAATGCGACAGTGGCCTCCACCACAAAA |

| | | | | |
|---|---|---|---|---|
| FBgn0038128dsechsim5 | 3R | 09 | 3R_09 | ATGACACAATCWGCGGGTTGGCTGCCAGCTTGCGAGGATTGGAGCAAACACGAGGAGCTCTCGTATGGCTGGGAGCGTGCCATAGACTCCAAGG<br>GTCGTTCCTATTACATAAACCACCTGAATAAAACGACCACSTACGAGGCACCCGAGTGCATACGCTGGGATGAGCATCCTCCGGAGCCRCGACAGG<br>TTCAACTCCAGCGGAATGCCAGCCTGGGTTTTGGATTCGTGGCYGG[C/G]AGTGAACGACCCGTAATCGTGCGATTTGTGACGGARGGTGGTCCTA<br>GCATCGGAAAACTGCAGCCGGGCGATCARATCYTGGCGGTGAATGGCGAGGATGTGAAGGATGCGCCGCGCGATCATGTCATCCAGCTGGTGCG<br>AGCCTGTGAGTCGCAAGTYAATCTCCTGGTGTGCCAACCCATGGCGCACTGCGTCCTGCCTGGCAGAAAATCCACACTTCTTTCAGCGGGCAAGC<br>GAGCCAAGCTGAGATCGCGTCCTAGTCGAGTTCGATTCGCCGAGAGTGTYTGCGTCAACGGAGCTCCACTGTTTCCACCGTCGGCCTTCTCCCTT<br>GGTGACATCTGCGTACCCCCGATGGCCAATGTGCTGAAGGTGTTCCTCGARAACGGGCAGACCAAGTCCTTCAAGTACGATGCCACCACCACGGT<br>GCAGGACGTGGTCAGCTCGCTGCTGGACAAGCTGTGCCTGTGCTGTGGCGAGCTCTTCAGCCTGGTGCTGGAGCACGTGAAGAGCCTCAAGCG<br>CAACAAGCTGACCCTCCTGGATCCGCAGGAATCGCTGGCCAGRATTGCTGCGCGTCCGGGAGCGCACAAATTGAGGTGCCTATTCCGCATCACC<br>TTCGTGCCCATCTCGGCCGCGGAACTGGCGCAGCGGGATCTGCATGCMCTGGACTATCTGTACTTGCAGTGCTGCAACGACGTCAACCAGGAGA<br>GATTCGCCCCAGAGCTGCAGCCGGAATTGGCCCTGCGCTTGGCCGCCCTCCACATGCACCAGCACGCYCTGGCCAACAACTTTGCGCCCGCC<br>AAGCTCACCGTCAAACTAGTCGAACGCGAGTTCGGACTGGAGCGCTTTGTGCCAGTGAGTTTGTTCGAGGGCATGAAGCGGAAGGAACTGCGCCG<br>GCTGATTTCTCACTTCCTCAAGCTCAACGCGGAGATGACTGGCTSCTCCAGCAAGGTTCTCACACARCTGCAGGCCAAACTCCATTATCTAGATATAAT<br>CGCTAGCTTACCAAGCTATGGAGCCAAATGTTTCAGCACAAACCAAAGAGAGGGCGTCGAACGCGTCCTGCTGGTGAGTCCACGCTTTGGACTCAG<br>CCAGATATCCAGTGCCCGCAATTCGGTGCCCCAGCCTATTTCCGCCATCGAGGACTTCACCCATGTGGTGGTGAATCGCGAGGACGATGTCACCT<br>GCAGCGTATCCATATTCATGCTCGGAGATCGAGCWGTAAAGTTCATAATGGAGGATCGAGATGGCTGCGAGTTTAGCCTTGTCCTAGGCGGCTACTA<br>CCGACTGCTGACAGGTAATGTKTTAAATATTCTGCGAGAACGGGAACCGAACGACGAGGACAATGCGCCGGGCTTCCTATCCCAGCATACAGTGCT<br>CCCAGCGGGATGGAGCTACCTGCTCCCCTTCCACACTCGAGCGCACAGCGTTAACTTCCTGATGGCGCCACCCTACCATCCGCTAGTCCAAAGG<br>GGTCAGCTACAGCAGCAGTCCTCCATTCAAGCGCCTTCCTCACTGCCCTACGTCATGGATCTGGATTTGCACAGCGTGATGGCCACCGAGCTGCT<br>GGAGGAGGCAGCCAAGGATTCGGGACTGAGTGACAGCAGTGGCAGCAACTAYTGCGAAGGACGCAGCCATTTGGGCAGCCATCAACTGGAGGCT<br>AAGAACGAGCAGGTTCTGAGGAGGGTTCAGGAACTGCAGCAYCTAGTTCAGACCTCAGAGCAATATCTGAGTGAACAGGAGCAGGGCGGCGGCGG<br>WGGAGGCGTGGCCATGCTGGAATTCGATTCGGATTGCGATAGCCTCAACAGCAGTAAGGTTTCATCGACGGAAGGGTCGCAGGGTGGAACGATTCT<br>GGGACCTGGCGCTATCATTCCGTCAGGACATCCTTTGAAACATAGCGATTCCCTGACGCTATTGGCGGAGACCATTAACCACGAGTTGAGTGGYATAA<br>CCCAGGGACTGAACGCCATACCCGAGTCYGCTCCAGTCTCCACATCTGCTCCTGCGACTCCAGCTACGCCCAAGCGAAAGCCAAGTCTGTGCAG<br>CACTTCCAGCCCGAGACCGCAAAGGAAACTGAATGGTTTTAGCCAGCTATTGAGTGACTTGCAAGCCCTGGGCACCGATTTCTCCCAGAGCGAAAG<br>TGATTCGGAATCTGTGGCCTCACCGGCGCAATCTCCTACAACCMGAAGACCGCAGATGATGCTGCTTCAGGCAKCGGGATCTTCCRCTGGATCCG<br>GAGCACCAGCCAAACAGCAACTSTCGCAAAGGAGCAGCTTTGGCTTGCACAGTCCCGATGGCACMCATTTCGGTGCCGAAACMAAGGACTACAAT<br>CTAAGGGAGTATTTGCAGCAACTRAAGGAGATTAGCAATGCGGCATCGGCGGACACAGATGTTGCTGCCCGTCAGCTCTCCGAGATTTATGGATTTGA |

| | | | | |
|---|---|---|---|---|
| FBgn0051304dsechsim8 | 3R | 10.9 | 3R_10.9 | ATGACACAATCWGCGGGTTGGCTGCCAGCTTGCGAGGATTGGAGCAAACACGAGGAGCTCTCGTATGGCTGGGAGCGTGCCATAGACTCCAAGG<br>GTCGTTCCTATTACATAAACCACCTGAATAAAACGACCACSTACGAGGCACCCGAGTGCATACGCTGGGATGAGCATCCTCCGGAGCCRCGACAGG<br>TTCAACTCCAGCGGAATGCCAGCCTGGGTTTTGGATTCGTGGCYGG[C/G]AGTGAACGACCCGTAATCGTGCGATTTGTGACGGARGGTGGTCCTA<br>GCATCGGAAAACTGCAGCCGGGCGATCARATCYTGGCGGTGAATGGCGAGGATGTGAAGGATGCGCCGCGCGATCATGTCATCCAGCTGGTGCG<br>AGCCTGTGAGTCGCAAGTYAATCTCCTGGTGTGCCAACCCATGGCGCACTGCGTCCTGCCTGGCAGAAAATCCACACTTCTTTCAGCGGGCAAGC<br>GAGCCAAGCTGAGATCGCGTCCTAGTCGAGTTCGATTCGCCGAGAGTGTYTGCGTCAACGGAGCTCCACTGTTTCCACCGTCGGCCTTCTCCCTT<br>GGTGACATCTGCGTACCCCCGATGGCCAATGTGCTGAAGGTGTTCCTCGARAACGGGCAGACCAAGTCCTTCAAGTACGATGCCACCACCACGGT<br>GCAGGACGTGGTCAGCTCGCTGCTGGACAAGCTGTGCCTGTGCTGTGGCGAGCTCTTCAGCCTGGTGCTGGAGCACGTGAAGAGCCTCAAGCG<br>CAACAAGCTGACCCTCCTGGATCCGCAGGAATCGCTGGCCAGRATTGCTGCGCGTCCGGGAGCGCACAAATTGAGGTGCCTATTCCGCATCACC<br>TTCGTGCCCATCTCGGCCGCGGAACTGGCGCAGCGGGATCTGCATGCMCTGGACTATCTGTACTTGCAGTGCTGCAACGACGTCAACCAGGAGA<br>GATTCGCCCCAGAGCTGCAGCCGGAATTGGCCCTGCGCTTGGCCGCCCTCCACATGCACCAGCACGCYCTGGCCAACAACTTTGCGCCCGCC<br>AAGCTCACCGTCAAACTAGTCGAACGCGAGTTCGGACTGGAGCGCTTTGTGCCAGTGAGTTTGTTCGAGGGCATGAAGCGGAAGGAACTGCGCCG<br>GCTGATTTCTCACTTCCTCAAGCTCAACGCGGAGATGACTGGCTSCTCCAGCAAGGTTCTCACACARCTGCAGGCCAAACTCCATTATCTAGATATAAT<br>CGCTAGCTTACCAAGCTATGGAGCCAAATGTTTCAGCACAAACCAAAGAGAGGGCGTCGAACGCGTCCTGCTGGTGAGTCCACGCTTTGGACTCAG<br>CCAGATATCCAGTGCCCGCAATTCGGTGCCCCAGCCTATTTCCGCCATCGAGGACTTCACCCATGTGGTGGTGAATCGCGAGGACGATGTCACCT<br>GCAGCGTATCCATATTCATGCTCGGAGATCGAGCWGTAAAGTTCATAATGGAGGATCGAGATGGCTGCGAGTTTAGCCTTGTCCTAGGCGGCTACTA<br>CCGACTGCTGACAGGTAATGTKTTAAATATTCTGCGAGAACGGGAACCGAACGACGAGGACAATGCGCCGGGCTTCCTATCCCAGCATACAGTGCT<br>CCCAGCGGGATGGAGCTACCTGCTCCCCTTCCACACTCGAGCGCACAGCGTTAACTTCCTGATGGCGCCACCCTACCATCCGCTAGTCCAAAGG<br>GGTCAGCTACAGCAGCAGTCCTCCATTCAAGCGCCTTCCTCACTGCCCTACGTCATGGATCTGGATTTGCACAGCGTGATGGCCACCGAGCTGCT<br>GGAGGAGGCAGCCAAGGATTCGGGACTGAGTGACAGCAGTGGCAGCAACTAYTGCGAAGGACGCAGCCATTTGGGCAGCCATCAACTGGAGGCT<br>AAGAACGAGCAGGTTCTGAGGAGGGTTCAGGAACTGCAGCAYCTAGTTCAGACCTCAGAGCAATATCTGAGTGAACAGGAGCAGGGCGGCGGCGG<br>WGGAGGCGTGGCCATGCTGGAATTCGATTCGGATTGCGATAGCCTCAACAGCAGTAAGGTTTCATCGACGGAAGGGTCGCAGGGTGGAACGATTCT<br>GGGACCTGGCGCTATCATTCCGTCAGGACATCCTTTGAAACATAGCGATTCCCTGACGCTATTGGCGGAGACCATTAACCACGAGTTGAGTGGYATAA<br>CCCAGGGACTGAACGCCATACCCGAGTCYGCTCCAGTCTCCACATCTGCTCCTGCGACTCCAGCTACGCCCAAGCGAAAGCCAAGTCTGTGCAG<br>CACTTCCAGCCCGAGACCGCAAAGGAAACTGAATGGTTTTAGCCAGCTATTGAGTGACTTGCAAGCCCTGGGCACCGATTTCTCCCAGAGCGAAAG<br>TGATTCGGAATCTGTGGCCTCACCGGCGCAATCTCCTACAACCMGAAGACCGCAGATGATGCTGCTTCAGGCAKCGGGATCTTCCRCTGGATCCG<br>GAGCACCAGCCAAACAGCAACTSTCGCAAAGGAGCAGCTTTGGCTTGCACAGTCCCGATGGCACMCATTTCGGTGCCGAAACMAAGGACTACAAT<br>CTAAGGGAGTATTTGCAGCAACTRAAGGAGATTAGCAATGCGGCATCGGCGGACACAGATGTTGCTGCCCGTCAGCTCTCCGAGATTTATGGATTTGA |

| | | | | |
|---|---|---|---|---|
| FBgn0051304dsechsim8 | 3R | 10.9 | 3R_10.9 | ATGTCGTGGCTTTTGGGCAGGAACCGCCAACAGCCGCAGCCGGATCAGACCGCCGGCTTTTCGGATGGAGGAGGAKCTGCCGATCCGGAGGGC<br>AGGACGGCCGGCGARAAGTCCGG[G/T]GATTCGCAACTGAGCCGGGCGGAGCGCAAAGCCATGGAAGCGTACCGCTTCGATTCGTCGGCGCTG<br>GAACGTGCAGCGGATGCTGCCAAAACCCTGGAGCGTTCAAAACACGCCCGGGAGGCCCTCGAGCTKTCCAAGATGCAGGAGGCCACCCGGCA<br>AACGGAGTACAACACCAAGGTCAAGGAGTACGAGGCYCATATMGAGCAGGCCAAGGTCGAGCAGAAGCGCATCGACCACGAGGAGCGCCGCAA<br>AACTCTCATCGAGGAGACCAAGCAGCAGCAGCAGCGTGCCCAGTACCAGGATCAGCTGTCCCGCAAGCGGTATGAGGATCAGCTGYTGCAGCAG<br>CAGCGRGTACAGGAGGAAAACCTGCGCAAGCAGGAGGAGAGCGTCCAGCGCCAAGAGGCCATGCGGCGCCAGACCATCGAGCACGAGATCGA<br>GATGAAGGAGAAGAACCGGCTTAAGCTTCTGGAACACGAGTTGCGGGCCAAGGCACGAGTGGATCGCGAGAACCGGGATATCAATTTGGAGAAGAT<br>CCGTCTCAAGGCGCAGGAGCATCGAACAACCGTACTGGAGGGCATCAAAACCGCCGGTACTGTCATCGGTGCCGGTGCTGAGGCTATGCTCACC<br>GACTGGGACAAGGTGCTGACCGCCGCCGGAGGACTTTCGTTGTTGGCCCTGGGTGTGTACACGGCCAAGGGCGCCACYGGAGTGGTCTCTCGA<br>TATGTGGAGGCGCGCATTGGTAAACCTACGTTAGTTGGSGAGACKTCCCGCTTYGCYTTCCTGGACGCCCTTAAGAACCCGCTGCACTACCTGAASA<br>GGCTGCGCGCCAAGCCGACCGATGCCCTGCAGGGCGTTGTGCTAAATCCGAAGCTGGAGGAGCGGCTTCGTGACATTGCCATCGCCACAAAGA<br>ACACACGTATCAACAGGGGCWTGTACAGGAACGTTTTGATGCACGGCCCACCAGGAACGGGCAAGACCATGTTCGCCAAGAAGCTGGCCGAGCA<br>CTCTGGCATGGACTTTGCCATCATGACCGGTGGCGATGTGGCGCCCATGGGCAAGGAGGGCGTAACCGCCATCCACAARGTGTTCGACTGGTCG<br>CACACCTCACGCCGCGGACTGCTGCTCTTCGTGGACGAAGCGGATGCATTCTTACGCAAACGTTCTTCTGAGAAAATYTCGGAGGACCTGCGTGCT<br>GCCCTGAACGCCTTCCTGTACCGYACTTCCGAGCAGAATCCCAAGTTCATGCTTGTGTTGGCCTCCAACACTCCCGAGCAGTTTGATTATGCCATCA<br>ACGATCGTCTGGACGAGATGGTGGAGTTCACGCTGCCTGGCCTGGAGGAACGGGAACGCCTCTTGCGCCTTTACTTCGACAAATATGTGCTGCAG<br>CCTGCCGCTGCGGGTGCCAAGCGGTTCAAGCTGGACACCTTTGATTACGGAAAGACGTGTTCGAAGATGGCAGCTCTGTGCGAGGGTATGTCGGG<br>TCGAGAAATCTCCAAGCTGGGCGTGTCCTGGCAGGCGGCAGTCTATGCCTCCGAGGATGGTCTGCTCACCGAGAAGATGGTCCTGGACAGGTGC<br>TACTCCGCTGCCCAGCAGCATAAGCAGAAGATGGCCTGGCTTTCGGATCAGGAGCGTGCTGATCACAAATCCATCACAGGCACAGCTGCCCCAC<br>CCCTCACSCTAACTGCCAACAAACYRTCCGGCGCGGAGGAGCCGATGCTGAAA |
| FBgn0040237dsechsim4 | 3R | 12 | 3R_12 | ATGTCGTGGCTTTTGGGCAGGAACCGCCAACAGCCGCAGCCGGATCAGACCGCCGGCTTTTCGGATGGAGGAGGAKCTGCCGATCCGGAGGGC<br>AGGACGGCCGGCGARAAGTCCGG[G/T]GATTCGCAACTGAGCCGGGCGGAGCGCAAAGCCATGGAAGCGTACCGCTTCGATTCGTCGGCGCTG<br>GAACGTGCAGCGGATGCTGCCAAAACCCTGGAGCGTTCAAAACACGCCCGGGAGGCCCTCGAGCTKTCCAAGATGCAGGAGGCCACCCGGCA<br>AACGGAGTACAACACCAAGGTCAAGGAGTACGAGGCYCATATMGAGCAGGCCAAGGTCGAGCAGAAGCGCATCGACCACGAGGAGCGCCGCAA<br>AACTCTCATCGAGGAGACCAAGCAGCAGCAGCAGCGTGCCCAGTACCAGGATCAGCTGTCCCGCAAGCGGTATGAGGATCAGCTGYTGCAGCAG<br>CAGCGRGTACAGGAGGAAAACCTGCGCAAGCAGGAGGAGAGCGTCCAGCGCCAAGAGGCCATGCGGCGCCAGACCATCGAGCACGAGATCGA<br>GATGAAGGAGAAGAACCGGCTTAAGCTTCTGGAACACGAGTTGCGGGCCAAGGCACGAGTGGATCGCGAGAACCGGGATATCAATTTGGAGAAGAT<br>CCGTCTCAAGGCGCAGGAGCATCGAACAACCGTACTGGAGGGCATCAAAACCGCCGGTACTGTCATCGGTGCCGGTGCTGAGGCTATGCTCACC<br>GACTGGGACAAGGTGCTGACCGCCGCCGGAGGACTTTCGTTGTTGGCCCTGGGTGTGTACACGGCCAAGGGCGCCACYGGAGTGGTCTCTCGA<br>TATGTGGAGGCGCGCATTGGTAAACCTACGTTAGTTGGSGAGACKTCCCGCTTYGCYTTCCTGGACGCCCTTAAGAACCCGCTGCACTACCTGAASA<br>GGCTGCGCGCCAAGCCGACCGATGCCCTGCAGGGCGTTGTGCTAAATCCGAAGCTGGAGGAGCGGCTTCGTGACATTGCCATCGCCACAAAGA<br>ACACACGTATCAACAGGGGCWTGTACAGGAACGTTTTGATGCACGGCCCACCAGGAACGGGCAAGACCATGTTCGCCAAGAAGCTGGCCGAGCA<br>CTCTGGCATGGACTTTGCCATCATGACCGGTGGCGATGTGGCGCCCATGGGCAAGGAGGGCGTAACCGCCATCCACAARGTGTTCGACTGGTCG<br>CACACCTCACGCCGCGGACTGCTGCTCTTCGTGGACGAAGCGGATGCATTCTTACGCAAACGTTCTTCTGAGAAAATYTCGGAGGACCTGCGTGCT<br>GCCCTGAACGCCTTCCTGTACCGYACTTCCGAGCAGAATCCCAAGTTCATGCTTGTGTTGGCCTCCAACACTCCCGAGCAGTTTGATTATGCCATCA<br>ACGATCGTCTGGACGAGATGGTGGAGTTCACGCTGCCTGGCCTGGAGGAACGGGAACGCCTCTTGCGCCTTTACTTCGACAAATATGTGCTGCAG<br>CCTGCCGCTGCGGGTGCCAAGCGGTTCAAGCTGGACACCTTTGATTACGGAAAGACGTGTTCGAAGATGGCAGCTCTGTGCGAGGGTATGTCGGG<br>TCGAGAAATCTCCAAGCTGGGCGTGTCCTGGCAGGCGGCAGTCTATGCCTCCGAGGATGGTCTGCTCACCGAGAAGATGGTCCTGGACAGGTGC<br>TACTCCGCTGCCCAGCAGCATAAGCAGAAGATGGCCTGGCTTTCGGATCAGGAGCGTGCTGATCACAAATCCATCACAGGCACAGCTGCCCCAC<br>CCCTCACSCTAACTGCCAACAAACYRTCCGGCGCGGAGGAGCCGATGCTGAAA |

| | | | | |
|---|---|---|---|---|
| FBgn0040237dsechsim4 | 3R | 12 | 3R_12 | ATGAAGATRTTTGTGACGGTTCTGATGGGCGCCCTGTGCCTGAGCCTCAGCCARGCTGACATCAGCTTGACCCAGGCGGGTTACAACTACCGCATC<br>CAGCAGCAGCAGCAGCAGCAGCAACCGCCCATTGTGGCCCACCTGTTGAACCAGCAGCTGCAGCAGCAGCAGCAGCTCCAGMCCCAGCAWT<br>CCTCGCACCCGGTGCTTCCACCTCTCCCACTGCCCTACCTCCCGCCCGCTGGCCAGTTCCATTCTGCCCAGCCTGCCGTTCGTCCCACMACY<br>GGATCGTTCCCCATGCCCGCCGGCTTYCCCGTCAACTTCCAGACCGCTCCCATCCAGCAGCAGCGACGYGCTCGTCCGCGTGTGAGGATTCCC<br>AAGCGWCCCATTGTGACCAAGAACTTTTTCATMCACTCGGCGCC[C/T]GAGGAGTCCGAGGATGAGGTCCAGGACGAGCTGAACCAGTTGGCCCA<br>GCAGCCCCGTAACCACTACAATGTGCTGTTCGTCAAGACGCCGGCTCAGACCAATCGCGCGGCGGCTCTCAACTTGGCCAAGACCCTCAAGCAG<br>GAGAAGACCGTGGTCTATGTGCTGGCCAAGAAGACGACCGCCTCCGATCTGCAGGATGCCATTGCCGARGCGCCACAGCACATCAACAAGCCG<br>GAGGTGTTCTTCATCAARTACCGCACTCCAGAGGAGGCCCTGAATGCCCAGCGGCAGATTCAGTCRCAGTACGATACTYTGGGAGGCAGTTCCACC<br>ATYACCGACGAGGGTGTGGCYCCCATCACCTCGGTGGTTGGCTCCCTGGATCCCCCAGAGGAGGAGGAGGAGGAGCAGCAACAGCAGCAACAT<br>TCRGAGGGACACTTTGCGGAGAACGGTGCCGGAAACTCTGGCGTCGGTCCCATCGGCAATCATTATCTGCCMSCGAACCAGTTY |
| FBgn0038487dsechsim1 | 3R | 12.9 | 3R_12.9 | ATGTCGAATTCCGTGAATATATCAGTGGAAACGACGTGCGAGAACCAGATACGTGAGATTGGCTACGATGGCACCGAATTGTATCAGCCGCCGCCGA<br>CTTTGTCGGAAAGTCTGGCGAAATGTGCGGCACGCATAGATTTCAGCAAGACGTCGCTGGACGACCTGAAAAAGGAGGAGAAGTCGGCGGCGGCC<br>GCTGCCGACGAGGACAAGGACGCCACTCAGTTCCAAGAGAGCCTGTGGCCTTGGGATGCGGTGCGCAACAAGCTGAAGGATGCGCTCACGGAG<br>ATATGTGTTCTGTCCGATGTGATCTCCATTGCCAAGGACAAGCGCTATCTAGTGCTCGATCCTCTGCTGGAGGAGGCTGACGACACCAAGCCCATTG<br>TGCAGGTGTACAGCCGCAAGAAAGCCATCTCTCAGGCAGCCCAAGTGCTCCTCAGTGGCGCTGAACGCCTGCGAAACGCGCACAGTGAGCAGC<br>GCAATCGCAATGTCTCAGATTTCCACATAGAACTACTGCGCCTGCGACAGAACTGGCGCCTGAAGAAGGTGTCCAATGCGATTATCGGGGATCTCAG<br>CTACCGCACCGCGGGCTCAAAGTTTGGCATGAGCGGCACCTTCGAGGTGACCAAAGCTGAAGAGACCGGGGATGAGGACACCCCCAGCTCATC<br>CAACAGCAGCAGTAGCACCTCCGGCAAYAATGGMATGCAACTTAAAGCCAGCTCCGCATTGCGAGTGATTGTTCCAGCGGAGCTGCAGGGTGTCG<br>CCTACATAAAGGTGATCACACAGAAGGATCAGGAGGATTTGTGCACYGCTCAGCTTAATCTGATGGGCCACGGACCCAATATAACAGCMCAGGTGGG<br>TGTCTGGCAGAAGACMCTGGAGTTCGCCCAGAATGTGCTCTTCTGCAAGGAGCTATT[C/T]GCYCAGTTGGCACGTGAAGCTATCCAACTGCAGGCA<br>CCAATTCCACATGTGGTCATCGGCAACCAAATYCGCGCCACTCTGCTGCCTAACATCCAACTTATTATCTCGTTRTGCCACTCAACCACCTTCGATTC<br>CAGCCAGCCGGCGCCAATTAACGATCATGACCATGTKCTGGAACACTCGCTGCATCAGCTTCTGCGCGAGGTGCACTACAAGAACTCACACCATC<br>CCTTTCCCCATCCGGCCAGTGCACCAYTGGGACCCACTAAGAAGCGCATGCTAGCMGGTCCGATGGCCGCYGATCGGGAAACKTTGTTGGACAT<br>GACCAAGTCGCAGACCATACTCGAACAGATCATCGCCCAGGCGCAGCACATTTTCATGCGCAAGCGAACACAATATGTGCTCGATACGCTGGCCA<br>GGGATGTGAAGGATCCGCAGATAGTGTCGCATTGGAACGCCATGAACAGTCCCACCATGTCGTGTGTGAAGATCAACATTGTAACGCACGGCTACGA<br>TGCCATTGGACGCACCTCGCTGGTCATTCACGTCAAGGAGCGCTCTCTTAAATGCATTTGYCGCGATGGACGCGTTATGCGGCTGTCCTACGARCC<br>GCAGGAACTCCGCGACCTGATCCTCTGCCAGATCAACTCTCACCAGATCTCCTGCCTGATCAGCCTAGCTCGCTGCATGTCCTGGACCGTGCTGT<br>CGAACAGCAACCACTTGGGTGTCGGCAAGGTGGAGCCGCTGGGAAATGCCAGTTCCTGCCTTCTGGCCTCTCCCAATAGCGATCGCATGATTGCC<br>GTTCAGATTCGCTGCGATCCGCAGATAGATGTTAAGGTTTATATAGCGAGAAGTCCACGACAGGACTTCTTTCCCAGTCCACTGGTGCCGGAAAAGCT<br>GTGGGAGAATCTGGGWGGCACATTCAAGGAGGTGAGGTTCGACAAAATCGAGGGCAAGAGCTTTCTCAACAAAATGGAGTTTCTGATGGCCTCRCTG<br>ACCAGYAACACRGCC |
| FBgn0038578dsechsim7 | 3R | 14 | 3R_14 | GAATGTCCAGTTGCGCGCCTGTGTGCGTGGCGTTCAGATGACTGGCGGCGCGCACATCACGGGACAGTTGATCTGGCGAGAGTACTTCTACACCA<br>TGTCGGTGAACAATCCAAACTACGATCGCATGGAGGGCAATGAGATCTGCCTGAGCAT[C/T]CCGTGGGCYAAGCCGAACGAGGATCTYCTGCAGC<br>GCTGGCGCTTAGGCCAAACGGGATTCCCGCTCATCGAYGGCGCCATGCGACAACTCCTGGCCGAGGGATGGCTCCATCACACGCTGCGCAACA<br>CGGTGGCCACCTTYCTCACGCGCGGCGGATTGTGGCAGAGCTGGGAGCATGGACTGCAGCACTTCCTGAAGTATCTGC |
| FBgn0025680dsimech1 | 3R | 15 | 3R_15 | GAATGTCCAGTTGCGCGCCTGTGTGCGTGGCGTTCAGATGACTGGCGGCGCGCACATCACGGGACAGTTGATCTGGCGAGAGTACTTCTACACCA<br>TGTCGGTGAACAATCCAAACTACGATCGCATGGAGGGCAATGAGATCTGCCTGAGCAT[C/T]CCGTGGGCYAAGCCGAACGAGGATCTYCTGCAGC<br>GCTGGCGCTTAGGCCAAACGGGATTCCCGCTCATCGAYGGCGCCATGCGACAACTCCTGGCCGAGGGATGGCTCCATCACACGCTGCGCAACA<br>CGGTGGCCACCTTYCTCACGCGCGGCGGATTGTGGCAGAGCTGGGAGCATGGACTGCAGCACTTCCTGAAGTATCTGC |

| | | | | |
|---|---|---|---|---|
| FBgn0025680dsimech1 | 3R | 15 | 3R_15 | ATGAGGGTAGTGGCCAACTTTTTGGATTCCCTTCGAAAGGTGCGGCTCAAGTTGTCCACGGAAAAGCTTGTCAATAGTAATCAAAMGCCAAACNNNNN<br>NNNNNNNNNNNNNNNNNNNNNNNNNNNNNNNNNNNNNNNNNNNNNNNNNNATTGGGATAGACCATACGGAGGTSCCTCAAAATACACCAGCTGGATC<br>CGATCAGATGTGGCACAATAT[C/T]AARAGGGTGCTGCTGATGTGTGGCGGGGAGARGCAAGCAGGGGGATTTGAGGATGCGGAACGGMCGAAGG<br>ATTCCAGCGGGCTGGTTGATGTCCTGGAGGCCCAGAGATTCGTTAGCCAGGTGTTCTCAGCCATGCAGGTGCCCAAACAGGCGGCCAGTGAGATG<br>GCCGATGCATTGATAGCCGCGGATTATATGGGACAGCGTTCCATGGGCATCCATAGATTGCCCTCAATTGCGGCAGATCTACTGAATTGCACTGTTGC<br>TGGTGAYGCCACRCCGGGAATYGTGTCCGAAAAGAAAGCCATCGCCTTAGTGGATGGCCACAATGCACCTGGACCCGTSGTGGCCAACTTTTGCAT<br>GGATCTGGCCCTGCAAAAGGCCCGGGAAGTGGGCATCGGCTGGGTTTCGGCTCGCAGTTCRAACTGCATTGGATTTGCCTCATGGTATGCCTGTCA<br>AGCGCTGGAACAGCGAATGATTGGTCTGTGCATGACGAATGCTGCTCCAACTCTGGTGCCTGCTGGTGGAATCGAACCCATTYTRGGGGAAAACMC<br>AATTGCCTGTGCCGCTTCCGGTGTCCATGAACAATTTGTGGCTGATTTTGGTATGGCGGCCTGTTCGGTGGATGAACTGGAGCTATCCTATTGCAATGG<br>CTGGTCAAAGGAAGTGCCCAAATTAGTGGCTTTGGATARAAATGGCAAGGAGACCACATCAACGGAGGAGGCCTTGAGGGCCCAAAGAATCCGAGC<br>CTTTCAGCCGGAGTACAAGGGTTTTGGCCTGGCTGCCGCAGTGGATATTCTGTGCGGCGTGATGACAGGTGCTCGATATGCAAATCAAATCCAGCGT<br>CGTGGCGTCTATAGTACGGAAAATGCTCCGGCGAATCTGGGCCAAGTTTATGTGGCCATCGATCCGATGCGTTTTTGTCCCTCCTTCGAGGACAGATT<br>GGCCGATTTCCACCGGCTGTTGAGACAAGCGATGCCCAGCAAGGTGGGAAAACCACCGATGGTGCCGGGTGACAAGGAGCTGCAACACATGAAG<br>ATRGTCGACGAACAGGGAGGACTCACAATGTCTCCCWGTACTCTGTCAGTTCTGGACGAACTGGCCACAGAGTTTGACATCGAACCTTTGAAGATGA<br>AGGCCAATAAAYCARWARRTSMMAGTATGCCTGAAGAATTG |
| FBgn0038775dsechsim6 | 3R | 16 | 3R_16 | CTGGAAAACGTTCTCGATCGGGAGCAGGCGGTTACTGTGTGTCAGGCTTTGCTATATGTGCTTCGCGAGGCAGTGGCCATGAGTGTTATAACAAACAA<br>TGGGCAGGAAGACTGTCACAGCGTGAGCAGCGCGGGCAGCTCCTACCTAGA[G/A]TTGTATCCCCGGGGCACCGAACAGGTMGTCAACGATCCA<br>CAAGTTTTGCATTCCGTCATGGTGGGCTTGCGCACCTTGATCGAACGCTTCAAGATCACKTGGAAGGAGTCGGTGGAGACCATTGGCCTGGTCAATT<br>GTCTGGCATTTGTCCTGGAGAACACAAGCATGGATGCCAGG |
| FBgn0038554sechsim1 | 3R | 16.9 | 3R_16.9 | ATGTCCACTGCAATGGCAATGGCATCAGCGGCATCCTCGAGTGCCGCGGCAGCAACAACAACGGCCATTAGCAACTCCACAAACACTACAGCGTC<br>ACCGGCAACAACCAACACCACGAAGACCATAACGACATCAGAGGTGCCACTGGAACCAGCTTCAAATATGGCTCCCATAGCATTAGAAATCGAAGAT<br>TCTGGAAGGAATGAKGCCGMAGACTTGGAGCATGGTGCTGCGAAAYCACTSGAGCARCGYAAAACGACAGTGGTGCAYACCTGCCCCCGTCCACC<br>GGGTGGCTACAAGTACTCCATGGAGTTTCTCTACGGGATCGGTAGTGGCATGGCCGGCATACCACTGAATATACCAACGCCCTCATCGATAACACCT<br>CGCACCGTACGATCCACGCCCCCCTTACTA[A/T]CMACCCACATGCCTTTGCTGACCAACATGGGCGTGTCACCACCACGTCCGTCAGGCATTCG<br>TTACGCAGGACCTGCTGGAGGCTCAAACGGTACCACGGCAGCAGCATCCACGCCGCCATCCACTACAGTTACAACATCAAGTGGCTCTCCGGGA<br>TCGGGAGCCGAGGCAATAACAAGTACAGTTTCATCTAGTGGCTTGCCAGCAGCGGGCTTGCCAGCTGCTCAGTTCATATGCCAGGGATATATGCCC<br>ACTGGACCGCAGCGYCGCTTGTGGCACGCGGAAAACGCRGTGTGGCAGTTTGATAGAAATTATCCTTACAATCAGGCRTACTCGCCGCCATATGGA<br>ATACCGATGATGCCGGTGGGCTTTGAGCAYCCGTATGGGCAGAGAATAATATACCCGGGCTATTACAACCAGACTCCTCCGGGAATTAATCCTGCTG<br>CRGTGGCGGGACTGACAAGAACTAGTCGTCCGGTGACCCAACAGCCACATATACTGACTCAGCCAGCCGTGGGAACAACGGAAAGTAGCGAGGA<br>AGCTCCTGCGACACTTGGAAACGCTCCGCAGGTSAGCAACACGTGGCGTTACGGAAGACCAGGAACACACCGCGGACGACATGCGGTGSCCGC<br>TGCAGCARCACCAGCTGTGTTGCACAGAGATTCAAAGACTTTTTACAACAGCATCGGCAGTGGTGYGGCTAACGGTCCTCGATTTAAAGCACCCTTTG<br>TGGCAAATGTCCGTAATTTTCAAGCAGGAGCTGTAGCAACTGTCGCAGCCGGAGGAGCCGCAACCACAGCAGTAGTGGCCACAAGTGCACCTGCA<br>ACTGGAGCAGCTAGCTCCAGCGATCAGAACGTGGCAAACAAACGCAACCATCAAGGAGCTGCGACCCAAAACCATCACAGAAATCGTCACAATGC<br>CAAAAAAGGAGGCAAGAATTCGGTTGCAAAAGAGCTGACATCCAACTCCTCAGAATCGCTTTCCAATTCATCCTCAAAGTCCCAGCTRAACAAGCGTC<br>CMAGCTCTAGCTCCTCTATTTCMCCAATCAAGCATCCACATCGCAATTATAGAAACCGGATGCGTTATACTGCAACGGAGCCATCTGAGCAAAAGGC<br>GGTCCCCACGACCGTTCCAGTAAGTAACTACCAGCCACCAACAGTGCGAAGAACTTCTAAGTTCCAAGGATCAAATGCCTATCAAGCACATRCTGCT<br>CCCGGTAGGCAACAGTCTCGTTAYTACCAATCGCGACACATGGATGGCTATGTYTTTCAATCTGGACACTATATGGTATATGCAGCGGGAGCACCGCC<br>TGTGGGGTTAGGAGCAGGAAAATCACCAGTGTCAGAAGCCACCGGGGCACCTCCTGGAGCAGCAGCAGCAGCASCASCAGMAGMAGCAGCAGT<br>AGGKGGAGGAGCAGCAGCAGCTCCAGTCACAGCCAGCAGCGCCACTTTGGAGGGCGAGCAGCCGCTGGACTCGGACTTTGATCAGCGCCAGG<br>AATTTGCTGACTTGGGTCTGGATCCAGCTAATGGCGGCTTCTCCTCCGACCTGGAGCCRACCGGTATCAAGCAGGACACCTCGGAGCTGGACACT<br>CGCTCKTGCCTGTTATCGCATCCAAATTCCGAGGTCGATGGCGATGATAATCAATCCGTCGCCAGTTTTGCTCCTGGTGTGGAGAGCGATGATTCCGA<br>TAGTGGACTAAGCGATGCCTCCGTGGAGTCGRTGGCGCGCGATATATTGGTTAGTTGTCTAGCARTGGCCACCGGGGCTCAGAAGCCCGATCTTAC<br>AGGCCCCAATCTCGTGCCCTACGGCGATATGCATTATCTCAAGGAGCTGGACAGAAAGAACCCCCAAACCAATGGCTACAGATCRCACAGGCCATA<br>TCATCACTCGCATTATGCTTACCATTCGCAAATGAGTCCAAGAGGATTGAGCTGCTGCGGCGATATGTTGAATCAACACTCTGRAGATTTGGTGTTTAAGT<br>TAGACCAGAATCAGCCGGATGGCATTGAAAGTGGGCAGAAGATTTTCTTGAGGGAGATAACCGAGCATCCTGACAACATCTCGATGGCCTCAAATCTC<br>TCCTGCAGTCCATCTGCYAGTTCCAGCAAGAGTGTATTGGCTCCCATGGCATCCAAGTCCAACATTGCAATGCCAGAAGAAAACAATGATCAGGATGA<br>ACTGCCCTTGGTGGTTCACAATCGTTATTGGCGCGAATTTTTTGGTTACACGCCAGCCGAYCGCTTTTTACTCCGAGCCAAGTTTGTGGAAATGAGACG |

| | | | | |
|---|---|---|---|---|
| FBgn0038934dsechsim1 | 3R | 18 | 3R_18 | ATGTGGAAGTGCCACAAGTGCGGCAAACCCGTTTACTTCGCGGAACGTAAACAATCGATCGGCTACGATTGGCATCCGGAATGTTTGCGCTGCGAG<br>GAATGCGGCAAACGCTTGAATCCCGGCCAGCATGCTGAGCACAAAAGTGTGCCCTACTGCCATGTRCCCTGCTATGGCGCCCTGTTTGGACCGCA<br>GTTGTTTGGACACGGCACCCGGGTGGAATCGCACAAAAGTTACGGGGTGAAGGGAGCCCAGAAGCCAACCGGAGCTCAGGCCAATGGCCCGCC<br>ACTCCCACGTGATCACCTGGAGTCCAAGCTTAAGGTCTACAACCAATTCTACGACAACAAAAGCCTGGAGATACGYAGCCGCGAGGTCAACAACCG<br>GTTGGTGCTGGAAGGCGCGCTTCGAGTTTACTGGGGMGTGCAGGGAGTGATTCATCTCAAGGAGGACGATGATCAGCGCATCTTAGTGCGCAAGAG<br>GAACTCTTGCCGTGTGTCCAAGGCTGCCAATGAAAG[C/T]AGTTCGGATAAGGAAAACGAAGCTAGCGAATCGGTGGCTCCGCCCACCASCACCAC<br>AGCYGAGGTCGATCAACTATCCACGGATGTTTCCCTTTCCGAGAGCATGACCTTCGACTCCTGTAGCCTGAACGAGATCTCCGAGCTGCCCACAAC<br>CCCAGAAGATGCCTCGGCCAACACCACAGCCAACGGCAAAGAGCAGACCAATGGTAATGTTTGCAACGACGACGAGGATACGACCACGACGGAC<br>TCATCCGGCACCCTGGTGGAGGCTCCTACGGCCAGCACAAGTTGTGTGTCCAGCACACTGCCTTCGAAGTTGGACCGYCTGGAGAAACTGGATTG<br>GGACGACATAGACGATCTGCTGCAGGTAGAGAGACGTCACAACGACAAGGATCGCATTTATGAAACYATGCCTGTCAAACTGCCATCGTCGCAATCC<br>TCCAGCGAYAGYTCGCCYAGYAAAACATCCACGGAGACMACTACGACCACTGAATCAAGTTCGACTCAATCGGCATCMACAAACACCAGCAGCAC<br>GGATGATTTTATGACCGCCACGGGMTCGYTGACCGCCAACACCAATACGCAGAAYACCACAACGGTGTCTACTACTGAGACCTCCCTGGATAATTTC<br>GAGACTTGCGATGACGCTACYCTGAAGCCCATTGATTTCGAGGACTTTAAGCGTAGTGTCCACCAGGACTATGTAAATGGGGCCAACTCCTTTACGGA<br>GCCCAATGAGGGCACCCTTAAGCGTAATCAACCCATTGATCCCTCTCGCATTCACGACTCCCTTAAGCTCTACGGCGARAATTCAGCKATGAGCAAA<br>AGCTTTAATTGTGAGCATGCGTTGCGCTCAATAGATCCCACTTTAATCAACGACACCATGAATCTGCGCAGTAGTGTGGGCTCTCCCCACTCCGCACA<br>GCGTCAGTACGCCTTACAAAAGAGTGGATCGGCCACAGTRACCAGCAGGGATCAAAAGAAGCCATATCAGCAGGGGCGSCAGTTGTTCGAAAAGGG<br>CATCAACCGCTCCAAATCGGGACCCAGCTGTTTTGTGTACTCCGACAGCGATGACGACGACGAGGCSACGCTGCGTCCCCAACGTTTGGCCACC<br>ATTAGRCGCAGCGACATACCWCCGCGGTACATACAAATCCAGATGGACTGTTATCCCAAAGARAATGTGGCTGCTGCCAGCGAAGGTGAGTCAAGT<br>CGSGCCGATGCTCCATCCATAACMAGTGGAGCAGCGGCCGGCGATGAGCTTACGCAAACCGAGGATTTGTATACTGCAAGCGAGGGCGTTGATG<br>GTCCAGACRKAGATGGATCGGCTGGGTTGCATGTGACGGAAGATGGAGTCGTGCTGAGAAGGCCCCCACGCACTGGAGCTTCTGCCATCAAGCG<br>ACGMAGTGGCAATCGTCGTTCTCGTACCAAGCTGAAACGTCGGTGCTCTATCAATGGTCACTACTACAATCGGGAGACRTCGTTTTTCACGCCRCCC<br>TACGGTTCTCAAATGTCAGTSTGGGTCTCCTCGATGGTGACCACCACTGAGGTCATCAACTTGGTGCTGGAGAAATACAAGGTGGACAGTTCGCCGG<br>GMAACTTTTCACTCTTCATTGTTCGGGACAATGGTGAACAGAAGCGYCTCAAGGACGACGAATATCCTTTGATAACGCGCGTCACTTTGGGACCTCATG<br>AGGATGTGGCTCGTATATTTTTAGTGGATTCTCGCAAAACTGATGAAATCAGTAATGAAGTTGCTCAATTTTTGAATTTATCTTTGCCCGAGTGCCGTGCTATT<br>CTGGAGCGCTACGATCAGGAAGTGGCYCGCGAAGTGGCCAAAATAAAGGAAAGATATGCCGAATTGCGTCGYCGTATTGTCAGTCGCATGGAATCTC<br>TYAAAGTGCATTTG |
| FBgn0039055dsechsim7 | 3R | 18.9 | 3R_18.9 | GAGGAGGACTTGGCGCGCCCATACACCGAGGACGAGGTAAAGGAGCTGTGTCTACGCACCAAAGTGAAGGTTGAYATGACCGCTTGGAATTGTCTG<br>TGGGAAGCCAAAAAGAGGTTTGAAGCAAAAGGACGTTTTGTAAACAAGTCTGAGAGATTYATCAACCGAATGTAT[A/G]TGAAAGCGGTGCGCAAAAAGAT<br>GGTCCAACCGTATCCGGAGGAGTTTGTCGCCCAGMGAAGAGAAGTAGTTGCCGCCGAGACTAAGAAGCACAACATCAGCCGATTGGATAGATGGC<br>RAAGGAAAAAGACTCAAAACCTATCAGCACAAGAATCCTCTCCAGCTCGTGGCGCTCAT |

| | | | | |
|---|---|---|---|---|
| FBgn0026257sechsim3 | 3R | 20 | 3R_20 | ATGCCGTTCTCTTCGCCGCAAAATCATCGGCTCATCTCGGCGGTCGCCGTTCGCCTGTCGCCGTTCGCAGTTCGCAGCTCGCCTTTCGCGGAATC<br>GGAWTCACAATCACATCCTTCTGCGAGTTCGAGGAGTGCACACCGGTGGTCGCATCCGCATCCGCAAAGTCCGCGCTTCGAAACCGCTATCGTAA<br>TCGCAATCGCCATCGCCAGCGCAACCAGAAATAGAACCATGGATATCTCCGCCTACCAGCACATGAACATCCGGATGAGCACGCGSGGCAAGCGT<br>CCTCTGAGGAACCTCACGCCCAACGACAAGGTGCGTGCCATCCAACGCATCCACAATGGTGAAACCAAGGCCAGTGTGTCGCGGGACATCGGAG<br>TGCCGGAGTCCACGCTGCGTGGGTGGTGCAAGAACGAGCAGRAGCTGCGCTTCATGTGCCGCCAGCTGGGCCCCGATCACCTGGGACTCGAC<br>ACGCCACCGGAAAAGCGCGCCAAGTTCGAGCTGCAGCTCCAGCTGCCACCAAAGTTCGTGGCACTGCCTCCAAACTACGAGGAGCTGGGCTTC<br>GGAGCACTGCCCTACAGTCCAGCTGATTACCCAGTGCAAAATGAATCCCTGCTGGAGAAGCTGAGCCTTGTGGAATTTGTGAAAAAGAACGGCGGAT<br>TGCATCCGGAGGGTGCRCTGCATCCGGGTCAGGCCGGCGTGATGGACTACTCCAACAACATGCTGCACCAGCTGAACCTCCTGGCCCTGCTCA<br>ACTCCAARCTCACCCCACAGACTGCCGATGTCCTGGTGGACGCACAACCCAAATCAGAGGACAACAAAGTGATTGATTCGCCCGCTGCCTCCGAG<br>GATTACAGCAAAAACAAWTACCCCTTGYTGAGTGTTAAGAACTGGGCCAAGGATCCGGCCAAGCGGGTGAACTCCGCCAATCAGCCGGAGCA[A/T]<br>GTGAACGATAAGAACAACAATGCCCTGGAGGCCAACAGTTCCGCACCACAAGCTCTGGCCACGGAGCAGGCGAAGACTCCCATGTTTCCGGACA<br>CCACAATGCCCCTACTGCCAGCACCCTTCCCGAATCCTGCGGATCTAGCGCCTGTGATAGCACCAGTGACTCCGGCCAACGCCCCGCCCACTG<br>GARCCGATAACCAGGGAGCGGCTCTGCTCGACTGGTGCAAGCTCTTCAATGCCAGCTTGAACTTCCTGGCCTTCGCCGCGGCCGCAGCCTCCAT<br>GCAGCCCGGCGGCGGAGGACCTGGTGCACCCAGCTACAATCCCAACCAAATGGCCAGCGGCGGWTCGGAGCCSGATCTCGAAACATATCCTTT<br>YMACGAGGCGCTACTGAAGCGCCYSWRYYSRCTGGCGCACAKTGAGGCCTCYAACGAAAGCTACYKYRAMAGCGAGCCGGAGGATCTGTCCGTG<br>CGCTCCTGCGCCTCCAAGGCGTCCAGCAGATCCCACACCCCCGACWAKARCGTCGGGTCCAGCGACGCCGAACAG |
| FBgn0039283dsechsim8 | 3R | 20.9 | 3R_20.9 | GGTGGCTCAAAGACCGCCGACGACTTCGACTTCCGGATGCAGCGGGTGCGGAATGTCCTGAAGGAAGTGCCCCTCATCGACGGGCACAACGAT<br>CT[A/G]CCGTGGAATATCCGCAAGTTTCTGAAGAACCAACTGAAGGACTTCCACTTTGGCGGCGATCTGCGGGAACTGGCTCCCTGGTCRTCGAGTG<br>CCTGGAGTCACACAGACCTGCGCCGCCTGAAGGAGGGCATGGTCTCCGCTCAGTTCTGGTCCGCCTACGCCCCCTGCTCATCCCAGCATCTGG<br>ATGCAGTGCAGCTGACGCTGGAGCAGATCGACCTGATTCGTCGACTTGTTCAGCTCTATCCRCATCACATGGCGCTGGTCACGACTGCTTCTGGAAT<br>CGAGCAGACGCACCGGACTGGCAAGATCGCCAGTCTCATTGGCGTGGAG |
| FBgn0039420sechsim1 | 3R | 22 | 3R_22 | ATGCCCGCCGCCGATGAACTGCAGTCGCCGTCCAGCGAAATGGTCCTSCTGACCCCCAGTTCSGGCAATCCTCCGTCCAGCATTGCCCAACCA<br>GCGGGYAATCCTGCSGAGAAAGCTCAGGGTCGCGAGGGATCCGCCGAGTCGGACAGCAGCAGAGTTGTGATCAAGAAGCAGCTGGGCCTCCTC<br>GAAGGAGTGGCCATCATCCTGGGCATCATCTTCGGATCCGGTATCTTTGTGTCCCCGAAAGGCGTCATCCAGAATGTRGACTCCGTGGGCGCTTCG<br>CTGGTCATCTGGGTGCTCTGCGGACTGCTGTCCATGATTGGTGCCCTTTGCTACGCGGAACTTGGTACTGCCATACCAAAATCTGGAGGGGATTACG<br>CGTATATCTTCGARGCCTACGGMTCACTGCCGGCCTTCCTGTATCTCTGGGATGCCATGATGATATTCGTACCAACGACCAATGCCATAATGGGCCT<br>GACCTTTGCCTCCTACGTGCTGGAACCCTTTTTCGGCGGAGCCTGCGATATTCCCAAAATAGCGCTGCAGCTCCTGGCGGCCATCACAATCTGCTT<br>CCTAACCTACCTGAATTCGTACTACATGAAGGTGACCACCAAGATGCAAAACATCATCATGTTCACCAAGATCGCCGCCCTGGTCTTGATCATCSTGGT<br>CGGCCTGGTCTGGATGCTGATGGGCAACGTGGAGAACTTCACRAGACCCTTTGATAACACTGAGACTGATCCCGGCAAGATGTCGGTAGC[C/T]TTTT<br>ACTCCGGTATCTTTTCGTAYGCCGGCTGGAATTACTTGAACTTCATGACGGAGGAGCTKCGCGATCCATATCGCAATCTGCCGCGCGCCATTTACATM<br>TCACTGCCTCTGGTCACCGGCATTTATGTGCTGGCCAACGTGGCTTATCTGGCCGTGCTGTCGCCCTCGGAGATGATCGCATCCAATGCCATCGCT<br>GTGACCTTTGGCGATAAGATTCTCGGTGTCTTTTCACTGATCATACCCCTCATGGTGGCCATCTCTGCTTTTGGCGGYCTTTCCGTCCACATTATGACCT<br>CCTCCCGCATCTGCTTTGTGGGTGCCCGCAATGGACACATGCCGGCGATTCTGTCGCACATCAGCGTRAAGAGCTACACCCCACTGCCGTCGCT<br>YGTTTTTCTGTGCTTTCTCTCCATTGTGATGCTGCTGGTGAGCGATGTGTATGTGCTGATCACCTACGCCAGCATTGTGGAATCGTTCTTTATAATGTTGTC<br>GGTGTCGGCGGTTCTWTATTTCCGCTACACMCGTCCCTGCATGGAACGACCAATTAAAGTCGCCATGTGGATTCCCGCCGTATTCGTCATCGTGTG<br>CGCSTTCTTAGTCGTCGTGCCCATCTATGTGGCGCCCTATGAGGTCGGAATGGGGGTATTAATCACCATAATTGGCATACCTTTCTACTACGTGGGCG<br>TCGTGTGGAAGAATAAGCCGAAATGGGTGCAGAGCACGATTGACTCGCTGACCTTCACTTGCCAGAAGCTCTTCATGTCCGCCAAAGAGGAGAAGGA<br>GGAC |

| | | | | |
|---|---|---|---|---|
| FBgn0039487dsechsim9 | 3R | 22.9 | 3R_22.9 | ATGACGGTAAACAAGCGCGACGCGGGTGCCACGCGACCGCGGTGCTTCTTCGACATTTCCCTGGGCGGCCTGGGCGTGGGCCGCATTGTCTTC<br>GAGCTCTTCAACGATGTGGCGCCCAAGACGGC[C/T]GAGAATTTCCGCGCACTCTGCACGGGCGAAAARGGATTTGGCCTGRTCACCGGCAARAA<br>ACTGCAGTACAAGGGCGTGATATTCCACCGGGTGGTCAAGGACTTCATGGTCCAGGCGGGCGACTTCTCCGCGGGCAACGGCACCGGCGGCGA<br>GTCCATCTACGGCGGGACSTTCGAGGATGAGAGCTTCGAGAACAAGCACGATCGTCCCTTCCTGCTKTCGATGGCCAACCGGGGCAAGAACACCA<br>ATGGCTCGCAGTTCTTTATTACAACACAGCCTGCGCCACATTTGGACAATATCCATGTTGTATTCGGCCAAGTTATATCCGGTCAGGAGTTGGTGCGCC<br>AATTGGAGGGTTTGCCTGTCGATCGGAACTCACGCCCGCTGCAGGATGCAGCCATTGCCAACTGCGGTGAACTAGTTAGRCAGACAAAGGCCAAAA<br>AGGARAAGAAGCACAAGAGGCGYTCCACGRCCACCGAAGACTCAAACAGCGAGGAAAGTGAGACTGAAGCTAAGKCCGTGCGCAAGGCCAAGAA<br>GAAAAAACGCAGCCGCAAAGAGTCAAAGTCGCAAAGCGAAAGCGAAGACAATGAAACAAGCCGCGGCAATGGAAAACATGACCAGACCCCGCAGG<br>ATGATGACGAGGACCGCGAGGAAGGCGAACTGCATCCGCTGGTCACAATTACAAAGATAGATCCTAACGAGATACCAGAGGTATCGAACAAATTCCTA<br>ATGCGCGSGGAGCGATCAAAGTCGCTGGATCGTGAGGAGCGTGGAGCACGCGGGCAGGATCGCGATCGAGGCAAAGATCAGGATCGAGACCGC<br>AGCCGTAACAACTTCGGTTGGTCAAAGAAACAGGCACCGCCCACATCGCGTAGCGGTCGCATTGTCAAAGGACGTGGAGTGTTTCGCTTTCGCACS<br>CCGTCKCGTAGTCGTTCGCGCAGCACTACGCCACCTCACTGGAAGCATGCCCAGAAGCGCACCATCAAGTTGTCCGACCTGGAGCGCATTGAGG<br>ARGAGAGCAAGATGCGCGAGGAGGAGATGAAGCGRCGCGAGCATGAGAGAAAGCGTCGCCACGAGGAGGCCACAAAGAATCCCAAGCAGTCCT<br>TTTTCGAGCTATCCCATGCCAGCACTTACGGTGGCAAAATAACTCCGGATCGATTCTCTCCAGAGCACAAAGCCAAGTCGAAGGACGCCGCAGAKC<br>GCGGTAAAGCCAAGGAGAGAGAAAAGGCTGGCGAKAGGGATAAGGCAAGGGATGTGACTCCTGTWAAGCGCCGCAAGTCGATGGACATGAACGC<br>CTTGGACTACGAGCAGCAGACAGAATCCGAAGCCGAGTCCGARAACGAGGASCAGTTGAAGGTGAAACCACCWAACAAGSCGGATATCAGAGTRG<br>AGCCATCGACCAGCAAGGATCGCARCGCTAATCGCGATGAGCGTAAGCCCGAGGAGAAGGCCAAGCTAAAGCGCAGTCGTTCCCGGAGCCCTC<br>GTCGCCAATCCCCTCCTCGTCGTCASCAACGCTCGCCCGCCCGACGTCCTGGCGAAAACTACAACCAAAAYCCGAATCAGAATCAAAATCAGAAT<br>CAGTTTAATCGTGGCAATCGCMGAAATCCTTTTGCGGACAGAGATCGCCGCCGTCGCTCGCGCTCTCAGGATAATGAGTACCGCAATCGTTWCCC<br>CCTGTCACCAATCCGAGGAAGAGGCTCGCCCAGTGGAGACCGCCGKCGTCAGCAGCGCTCCCGCAGTCAGAACCGTAGGCAGGATTCACCCA<br>GCCAGAGGCGTCAGCTGTCGCCGGGCCGAAAACGTCAGGATTCGCCGAGTAAGCGTCCCAAYTCAACGGAGAGAAAGCGGCGGGACTCGCCT<br>GAAAGAAAGCGRGACGATTCACGCGACAGGAAGCGGGAGGCTTCTCCCAACTGGAAGCGTGAGGATTCACCTGGAAGGAAGCGCAAGGACTCCG<br>CCAATAGAAAACGGGAGGATTCCCAAGACAAAAAGATACARGACAGCAAAAGGCAGGRGTATTCWGCGGGTAAGAAACKAGAGGATAGCCCGAGCA<br>AAAAACTAAAGGATTCTGATAGGAAGCGTCAGGATTCTCCTGYSAAAAGGCAMGAWTCGCCGGGCAGAAAGCGTCAGGACTCGCCAGGCAGGAAA<br>CGGGATTCTCCTGACAAAAGAGCTGCGGAGTCCGCCGACAAAAAGCAACAAGACTCGCTGGCCAAAAGGCAGGATTCACCGGGCAGGCGGCGTA<br>GGGACTCGCCGGAAAGGAAACGCAGGGAMTCGCCTGACAGAAGGCGCAAGGAWTCRCCGGCCAGACGACGCCAGGAATCMTCCGGCAGAAG<br>GCGCCAGGACYCAGATGGCAGAAGACGCCGTGAATCGTCCGGCAACAGRCGCAGTCGCAGCCGTGGACGTCGGCACAGTTCCTCGCGCAGTC<br>GCAGTCCCTCAACTAGTCGCTCCCGCCGCCGCCGCTCSCGCAGCCCGCTGCCCAAGCTCACCTCCGCGCTGCCCATGAGCGAGGAGCGCG |
| FBgn0039581dsechsim2 | 3R | 24.1 | 3R_24.1 | TGGCCGTAAACCTCACAAACAACAAATCTGCAGTGGGCGTGGGCCAGTTGGCTCGCTCCAGCGATGATCTGTACATGTGCGGAGGTCACGGTGTG<br>GCCATCAAGATGCTCCACCTCTTTGGGGACAAGCTCTGGTCGCATGAGCCCAGTCTGGTGCAGCAAATCCCGCTGGTCAAGACAAAGGCCTTGTCT<br>GGCGAGGATTTCCCTGCTCTTGGGTCTGAGGCGGAAAGGAGAAAGGCCGCCAGCCCAGCTACTCCCGCTCCTGTGTCCTTCGCTWCCGTGGCC<br>CAAACAGAGGAACTGGAAACGGAAACTGCTGCTATGACCTTGGAAAG[C/T]CAGGAAGRCCCCGAGTCCGACGAAGAAGCCGCGGAGGAAGCGTC<br>ACCGGAACTTATACTCAAGAACGCCTTCCTCTCCGCCTTGAAAAACAATGGAAAGAAGCTGCCCCTGCCTCTGCTGACGAGCAATTTCTACAGACTCT<br>ATGTGGTCACGGAGGCCACGGAACAAATCGACCTAAAGAAGACGCGCTACAAAAAGCTCTCCAACTTCCTGGCCG |

| | | | | |
|---|---|---|---|---|
| FBgn0041588dsimech2 | 3R | 24.9 | 3R_24.9 | ATGTCCGCCTTTCGMTACTTCTCGATCACGGATTTCTCCTTCGAGGGACCTCTGCTSCCGCTGCACGCGGCCACCACGAGCAAGGATGCCAAGGA<br>CTCGGACAGCCCCAGCTCCGAGCTGAACATACCCTACACCGTCTTCGAGGTCCTGGTGGCCATCGTCAGCATCATCGGCAACGTGCTGGTGATC<br>ATCGTCTTCCGCCGGGAAAGAAAACTGCGACGMCGCACCAACTACTACATAGTATCCCTGGCCATGGCCGATCTGCTAGTGGGCGCGTTGGGCAT<br>TCCCTTCGCCATCCTGGCATCCATGGGACTACCCAGAAACCTCCATGCCTGCCTCTTCACCGTCTCCCTGCTCGTGGTGCTGTGTACYATCTCCAT<br>CTTCTGTCTGGTCGCCGTCTCCGTGGATCGATACTGGGCCATCCTATATCCGATGGCCTACTCAAGGAATGTCCGCACCCGCACGGCGATATTCAT<br>CATCTCCATGTGCTGGGTGGCCGGGACRATAGTGGGCTTCCTGCCGCTTTTCGGCTGGCACGCCGATGTCAACCACAACCAGGAGTGTCTCTTCG<br>TGGAGGTGATGGACTACAACTACCTCGTCTTCCTCTACTTTGCCACCATTATTACTCCRGCTCTGCTGATGCTGGCCTTCTACACGCACATCTACCGG<br>GTCATCATYAAACAGGTCCGTCAGATCGTTACGATGAACCCCGCCTCCGATCTCAGTCGTCGCTCCTCGGCGGCCGTGGTGCAGGTGACRACAC<br>CCGGAAGAGGTGGRCACACGGGAACCATGMTGCGGGTCTTGGGCGC[C/T]GCAAGGAAGCGGGACGTCAAGGCCACCCAGAATCTGTCCATTAT<br>CGTGCTGTTCTTCATGATCTGCTGGATACCGCTCTATACGATCAACTGCATCAAGGCCTTCTGTCCCGAGTGCTATGTGCATCCCAAGCTGACGCTCT<br>TCTGCATCATTCTGTCCCATCTGAACTCGGCGGTAAATCCGGTCCTGTATGCCTATCACCTCAAGGACTTCCGGGCCGCGCTGAAGAACTTACTCCT<br>GAAGATGATGGGCGTGGACATTGATCAGCAGGCGGAGGCCATCCACCGCTTCTCGGTGGCCAGTCAGCATCGCCTGCAGTCCATGGACTCCAAT<br>ATGCGCTCCACGCAGCCGCGCCTCTATGTGGRTGAGTATTCACCCATTTGGTTGAGGCAGCAGCAGGAGGCCCTGAAGAACTCCCAGCTCCTGC<br>CCAAGTGCGGAGTGGTGTCGCCCTGTTTCAACAACATCAACCAAACGGTGGCCGCCGTGGCTTCGGTGACCACAGATCTCGAGCGAGAAATGTGG<br>AACATAGTGGAGGCTTCCAGTGGCGCCGAGCTGGGTGAAACCAGCTACGAGTTCCCTTCCCCCGCTCCGGGKTCCCAGCGCAGCAGTGAACGA<br>AACAGCAGTTCCACAGTCCCACCAGCTCCTCCTGCTCCTGCGAAGCCCTCAGTTCCGTCCGCCTCTTACGACAACCACAACTACAGTTTCAGCCA<br>GGATGARGACGAAGATGATGATGATCTGGAGTTTGAGGACGTATTTGTGCCGGCCAGCTCTGTTCCGAATCCCACTCARCCTGGCATAGATCCCGTG<br>GAACTGCGTCGCTCCCTGGCTTTGGTCATGAGGGAGAAGTTGCGATCGGATGACACGGACTCCAGGCCAATGGGCAACAATCAGGATCTTCCCATA<br>GATGAACAGTCCAGGGAGAGACCGCTCTCCACTCAAACATCTCCCACTAATGGCCCGCTTCCGGCTCTTCTGCGRGCCAAACTMTTTGCTGGMAA<br>CTCCAATAGCGCTCACTGCCTGCCAGGATCCRCGGCGAGTCCTGCTCCCCAGGAGCAATCCGGTGTCTTTGTGATCGATAGTGMGGCGAGTCCC<br>GGCWCAAATGGACACAAGCCTAAGTATCGAAAGGGCACGGCATTTACTAGGAGTTCGCTAAAGAAGAGYCGATCCTGCAAYTGTAGYTCCATCGCYA<br>AGGGACGAGGGGTCCACGACGAGCCCAGCAGTAATCTCTGCAGGGATCAGGAGTCCTCTGTGCATCCACAGCATCCGCAGCCAGCCAACCATC<br>CCACAGAGAACTTCTTTAGTCCGCTGAGATCGGTGGGCAGTTTCATGCAGCATTCCAACCTGTTTCACTTYCTCCAGCCTCACGCCGCCCGTCCCA<br>CCTCATCGACGGCCTCGTCCTCGGCCTCCACGCCCACTCCWTCACCACCTCCCACGGGCCMGSCTCAGGAGGAAACKGTTCCAGTGGGTCT<br>GACCACCTCCTCGCCGTCCCTTCTGGCGACCAGTGCGGAGAGT |
| FBgn0039747dsechsim9 | 3R | 25.9 | 3R_25.9 | ATGGCCCCGTCGAGATGGAGCCGGTCATCATGCCCAGCAAGCGTCAAAAGTATATTGTCAGTGCGCCGCCTGGGTCGGCCAACTGCAGCAGCAA<br>CTCGTCCAGTTCGAGTCCCTCGAACGTCAACGTGGGCTCCTCGCTTACCATGAAGTTGACGAAGGTGCAGCCACAACAGCATCAGGCCAAGTCCT<br>CCACGCMAAACAACAAAGGCAAAGTA[G/C]CCAAGGAAATGGTTGCTCTGCAGCGCTCACACATCGAATCCCAGGTCCTCAGCAACTTTGTGACTG<br>GGGTGTCCAATCTGAAGCGTCGAAAGTCGCGCTATGTGYCCGGGAATCTCGCCACTCAGGCAGTAGGTGGAGAGCTGTCCCGCAGCTTAAATGCC<br>TCCTCCAAAAATGAAAACAAATCTCTAAAGGTTGCTCTTAAGCGGAGCACTAGCATTCCAGC |
| FBgn0039863sechsim2 | 3R | 27.5 | 3R_27.5 | GTCAATATAAGCCAAGCTGAATATTCAAGCAAATATATAGAATACTTAAGTCGCCTAAGAAGTAACCAGGAGAAGGGCAATA[G/C]CTATTTTAACAACTTCA<br>TGGGAGCTTCCKTTATAAGAGACCTTCATTTTCTAAGCATTACCACAATCGAATTCAATGACATAAGCAGCAAACGATCGCGCCATAGAAGGTCGCTTAAA<br>AAAATTAACAGGTTAAAGCAAACCAAAAAGCATCAAAGTTATGGAGATTTACTACGTGGCAAACAGGAAACAACAAAT |
| FBgn0039914sechsim1 | 4 | 0.65 | 4_0.65 | ATGCGATTTGCCTTCGATTCTAATCATTCGCAATCGGAGGCGTCATTCAAAGGCAGCAGATGCTTCTTTAATTGTCAATGCATCTGCTGTCGCCAGGGAT<br>GCTGCGTTGTGGTTGTAAAGTGCTGTTGCTGCTTTAACTTAAACTGCTGCAACA[G/A]CCTTGGCTCCCGGAAGTCGTTTCCACAATCYGCTGCAATGC<br>GTAAAAAAGTTTCTGATCTCGAAGTCCTTAGAGTATCAAGGTTTGTGGCGGTTATTTTAGTGCTGGCTCGGTGGGTTACTGCCGTAGCGACCCTCCTGA<br>CAAGCTGCATTCTCCTAGACATTTTTTCCGTGCCTGGCCAGTCTGGAGTTGCAGATAGAAGCCAAGCCAGCAGTAGGACAGTGCACGTCTCGGTTCC<br>TACCACACCTAATGAAACTCCCAGCAGCAATTCGGAAACAAAGCTAAAGTTGCTTTATGGGTATACATCGTATGATATAAATAACGACCAACAGGTAAAGT<br>CCAACAATCTATATAGAGTGCTTTGTAGAAGTCGTAATCGAAAACGACAGCGAAGGAGACGCCGCCGCAATCGCAGACGACGCAGGCACAGATATAC<br>TAAGAGACTTCATCATGTAATGCAAAGTAATATGAGCGGCTTTGAGCAAAGACTTAGTTTTAGCGATGGCAAATGCCAGTCTTTGGAGACAAATTACGGAA<br>CTAATTATGACTTAATACACGGACGTAAACTATTTAGTCAGTCAGAGAGAAGCCTACTGGTGTACCCTTTGAGGGAAATTGAAGCACCTTGGCCAGCGAT<br>TCATGGTTCAATGCGTA |

| | | | | |
|---|---|---|---|---|
| FBgn0024913sechsim1 | 4 | 01.1 | 4_01.1 | ATGCGATTTGCCTTCGATTCTAATCATTCGCAATCGGAGGCGTCATTCAAAGGCAGCAGATGCTTCTTTAATTGTCAATGCATCTGCTGTCGCCAGGGAT<br>GCTGCGTTGTGGTTGTAAAGTGCTGTTGCTGCTTTAACTTAAACTGCTGCAACARCCTTGGCTCCCGGAAGTCGTTTCCACAATC[C/T]GCTGCAATGC<br>GTAAAAAAGTTTCTGATCTCGAAGTCCTTAGAGTATCAAGGTTTGTGGCGGTTATTTTAGTGCTGGCTCGGTGGGTTACTGCCGTAGCGACCCTCCTGA<br>CAAGCTGCATTCTCCTAGACATTTTTTCCGTGCCTGGCCAGTCTGGAGTTGCAGATAGAAGCCAAGCCAGCAGTAGGACAGTGCACGTCTCGGTTCC<br>TACCACACCTAATGAAACTCCCAGCAGCAATTCGGAAACAAAGCTAAAGTTGCTTTATGGGTATACATCGTATGATATAAATAACGACCAACAGGTAAAGT<br>CCAACAATCTATATAGAGTGCTTTGTAGAAGTCGTAATCGAAAACGACAGCGAAGGAGACGCCGCCGCAATCGCAGACGACGCAGGCACAGATATAC<br>TAAGAGACTTCATCATGTAATGCAAAGTAATATGAGCGGCTTTGAGCAAAGACTTAGTTTTAGCGATGGCAAATGCCAGTCTTTGGAGACAAATTACGGAA<br>CTAATTATGACTTAATACACGGACGTAAACTATTTAGTCAGTCAGAGAGAAGCCTACTGGTGTACCCTTTGAGGGAAATTGAAGCACCTTGGCCAGCGAT<br>TCATGGTTCAATGCGTA |
| FBgn0024913sechsim2 | 4 | 01.1 | 4_01.1 | CGTAATACAGTATTGAATAAGCAGGGAGCTGTTTCCGGATCGATGCAAGGGCAGTCTCGTCGCGATAGTCAAAATTCCACTGCAAGCASTTATTACGGT<br>AGCATGCAAAGTCGTCGTAGCAGTCAATCATCCCAGGTGTCTTCTATACCTACCATGCGACCGAAACCCTCGTGTAACTCGACTACGGCTTCCTTTTAT<br>GATCCCATTTCACCAGGGTGTTCCCGGCGCTCTAGCCAAATGTCAAATGGTGCTAACTGCAATTC[A/G]TTTACTTCCACATCAGGACTACCCATATTAA<br>GCAACGAATCGAACAAAAGCTTGAATGCATGTATTAGTAAACCGAATCTCGGGGTTCAGGGTGTTGGAATTTACAATAGTAGTCTACCCCCGCCTCCAT<br>CGTCTCACCTAATTGCAACAAACTTGAAGCGCTTRCAGCGGAAAGATTCWGAAAATCACAATTTCACTAGCGGACGATTTTCTATTCCTACTTATATGCAT<br>TCTATACACATAAAAAACAACAAACCAGTCGGAGAAAACGAATTTGATAAAGCAATAGCGAATAATACACTTCGACGTCAAACTGAGCCAGTGCCAAACTT<br>AAACTTAGATCCATTAACAAATATTCCTCGGTTCAGTACAACACCGCATAGCTTTGATGTTAATGTTGGAAAGACTAATAATATAGAATCGTCAATTAATAAGG<br>ATAGCTTAAGAAAGGATCTGTGCACTGTTTCAATTAAGGCTGACATGGCAATGACTGGTGACCAACATCCAAATGAAAGAATAAATCTGGACGAAGTTGAA<br>GAACTTATACTTCCCGATGAAATGCTTCAGTACTTAAACTTGGTAAAAGACGACACAAATCATTTGGAAAAAGAACATCAAGCTCTACCTATGTGGTCAAAT<br>GTATCCGAAACTATAGCTTCAAACCACTACCGAGAACAATCAAATATATATTACTCCAA |
| FBgn0004859dsimech1 | 4 | 07 | 4_07 | CCTGTGGGCTACGATCTAAGCACCAATGATCGAATTTTTCCTGCTCTAGATCCTTCTCCCGTGGTGGTCAAATCACTGCTGGCTTCCGTGGGCAATAG<br>GTTGGTACTATCCAAGCCTAAGGTGGCTATAGTGTCCACTGGAAGTGAGCTTTGTTCACCGCGCGATCAGCTTACTCCGGGAAAGATCTTTGACTCAAA<br>TACCACCATGTTGACGGAGCTTCTGGTTTACTT[T/C]GGTTTTAACTGTATGCATACGTGTGTGCTAAGCGATAGCTTTGAAAAGACTMAGGAATCGCTATT<br>GSAGCTCTTCGAGGTGGTGGACTTTGTCATTTGCAGCGGTGGTGTYTCAATGGGCGATAAGGATTTCRTAAAGTCCGTGTTGGAAGACCTTCAATTTACG<br>ATTCACTGCGGCCGAGTAAACATAAAGCCAGG |
| FBgn0000316dsimech1 | X | 0.1 | X_0.1 | CCTGTGGGCTACGATCTAAGCACCAATGATCGAATTTTTCCTGCTCTAGATCCTTCTCCCGTGGTGGTCAAATCACTGCTGGCTTCCGTGGGCAATAG<br>GTTGGTACTATCCAAGCCTAAGGTGGCTATAGTGTCCACTGGAAGTGAGCTTTGTTCACCGCGCGATCAGCTTACTCCGGGAAAGATCTTTGACTCAAA<br>TACCACCATGTTGACGGAGCTTCTGGTTTACTTYGGTTTTAACTGTATGCATACGTGTGTGCTAAGCGATAGCTTTGAAAAGACTMAGGAATCGCTATTGS<br>AGCTCTTCGAGGTGGTGGACTTTGTCATTTGCAGCGGTGGTGTYTCAATGGGCGATAAGGATTTC[A/G]TAAAGTCCGTGTTGGAAGACCTTCAATTTAC<br>GATTCACTGCGGCCGAGTAAACATAAAGCCAGG |
| FBgn0000316dsimech5 | X | 0.1 | X_0.1 | ATGAACAAGTACGGGATGRTCGGCGTCTGCCTACTGGCTGCTCTGGGCGCTCTGCTCCTGGAGGTCACGGCCTCTCCTTCGTCCGCCGCCTCGT<br>CTAAGGTKGATCCTAGCCAGCTTGGTGGACTTTCAGCCCAGTTCTTGCCMCCCGAGTACCGCAACACGAACGTTAGCATCGAGGATATTAAAAAAATA<br>TATCGTGAAAAATGCAAGAAGGTTAACGGAGCGGACAACGCAACCTTCTACGAAGAAATCGAGCGGGCGGCAGCCAAGATGAGCACCTGCATCAGC<br>GGKGTGGTCAATCTGACGGCTCTGCAGGAGGARATGGAGGTGGCGAGGCCGAACGGCGACCTGGACACCGTKTTTAGCAAATACTGTCTCAAGGC<br>ACCAGAGGCGGAGGCCTGCGTCAAGGAGTTCAACGACAAGGCCCAGCATTGCTTGACCGCCGAGGAGAAGCGCCACCAGGAGACGGTTACCC<br>GAATTGGAGCGTCCGTTCTGGGATTCGCCTGTTCGCGTGGTGGCGATCAGATCGCCCTCTTCATTGCCGAGCAGGGACCCGAGTGCCTGGAGGC<br>CAACAAGGAGGCCATTAGCAA[C/T]TGCCTCAATCAATCCTTTCATCAGTACATTCCCAARGAYGGCCAAGTTCCGGACCTGATGAGCCGCCCAGAG<br>CTCCTATTCTCGCCCACCCACTGCGTGGACCTGCAGCGCTTCGARGCCTGTGTCATCCATCATTTGGAACAGTGCACGCAGATCACCACCGCCAA<br>TATMGTTCAGTCCGTCTTCCGATTCGTGAAGAACGAGACCGACTGYCAGGCTTGGATGCAGGCACGYGCCAACGAGAAGCCCATCCTGCTGGCCG<br>CTTCCAGCAACAACACAGCMCCTGGACTCACCTACTCCCTGGCCGGTACTCTTTTGGGCGCCACAATACTCCTGATACGCCCC |

| | | | | |
|---|---|---|---|---|
| FBgn0040364dsechsim9 | X | 01 | X_01 | ATGGTCGAATACGCGCGCATGATGCCGCACTTTGCCCAGGTGCCGCTGGACGACCAGGTGATTCTGCTAAAAGCCGCTTGGATCGAGCTGCTCAT<br>TGCGAACGTGGCCTGGTGCAGCATCGTTTCGCTGGATGACGGCGGTGCCGGCGGCGGGGGCGGTGGACTAGGCCACGACGGCTCCTTTGAGC<br>GACGATCACCGGGCCTGCAGCCRCAGCAGCTGTTCCTCAACCAGAGCTTCTCGTACCATCGCAACAGTGCGATCAAGGCCGGTGTGTCAGCCAT<br>CTTCGACCGCATCCTGTCGGAGCTGAGTGTGAAGATGAAGCGACTGAATCTCGACCGACGCGAGCTGTCCTGCTTGAAGGCCATCATACTGTACAA<br>CCCAGACATACGCGGGATCAAGAGCCGGGCGGAGATCGAGATGTGCCGCGAGAAGGTGTACGCTTGCTTGGACGAGCACTG[C/T]CGCCTAGAA<br>CATCCGGGCGACGATGGACGCTTTGCGCAACTGCTGCTGCGTCTGCCCGCTCTGCGATCGATCAGCCTGAAGTGCCAGGATCATCTGTTCCTCTT<br>CCGCATTACCAGCGACCGGCCGCTGGAGGAGCTCTTTCTCGAGCAGCTGGAGGCGCCGCCGCCGCCCGGCCTGGCGATGAAACTGGAG |
| FBgn0003964sechsim2 | X | 01.9 | X_01.9 | GAGGCGTGCAAGGCGCTGCTGGACAACTTTGCCAATCGCGAGATTACCGATCACATGGACCGACTGCCRCGCGACGTGGCCAGCGAGCGACTG<br>CATCACGATATCGTCAGGCTGCTGGACGAGCATGTGCCGCGATCTCCGCAAATGCTGAGCATGACRCCGCAGGCCATGATCGGATCCCCGCCG<br>CCGGGTCAGCAGCAGCCGCAGCTGATCACACAGCCGACGGTCATTTCCGCCGGAAACGGCGGCAACAATGGCAACGGCAACGCCAGCGGAAA<br>GCAGAGCAGCCAGACGGCCAAGCAGAAGGCGGCCAAGAAGGCCAAGCTGATCGAGGGCAGTCCCGACAA[C/T]GGACTGGACGCCACGGGCA<br>GCTTGCGACGCAAGGCCAGTTCAAAAAAGACAAGTGCGGCCTCAAAAAAGGCCGCCAACCTGAATGGCCTGAATCCGGGCCAGCTGACGGGYGT<br>GKCGGGCGTGCCGGGTGTACCGCCCTCGAATTCAGCGGCACAAGCAGCTGCAGCGGCAGCGGCAGCCGTGGCGGCCATGTCCCACGAACTG<br>GAGGGATCGCCGGTGGGCGTGGGCATGGGCGGGAATCTGCCCAGTCCGTACGACACCAGCTCGATGTACTCGAACGCGATGGCCG |
| FBgn0004647sechsim3 | X | 03 | X_03 | ATCAACAACGTTGCGGACTGGTATTTTGTGCCGGTGGGGCGCGAGGAGACGAACATTAATGCATGCAGCCTGTTTTTGTGCAAGTCCCGCGGCAATT<br>TGCTGCGCTACACGGAGCACAAGAAGTGGGAGCTGGTGGAGACCGCCTTCGAGGTGCAGCCGCGSTCGATCTTCTTCGGCCAGATTGTGTACGA<br>GTTCTACGGAGAGGGCAGGACCATTCAGCGAATGGCTGCCCTGCACATTATGGATGGTATCTGCCTGGGCGGCATCGATATCCGTAGACGTCCGTA<br>CCGCGAGCGCATGAGCATGTGCGATAAGTTTGCCAGGAGYCTAAACAAGCCGTACCGCAAGGAGCGCACTTTTGGAGCGCTGCGCAGCAAGCCC<br>CTCTTTCGCCTGCAGGATATGGGCAGTTTCTTTGCGAACATGCGCCATTATGTGCTAAAGGACAACTCGCAGAGATTCGGAATTGCGCTGGACGACAA<br>CAAGTTCTTTGTGCCCGGCGGCATAATGATGTTCTGCGAGCTGACCAAAAATTACGTGTCCGCCCACTCGCGGTCTCGTGGTCAGCTGTACTACTTTA<br>ACGTGAGGAATAAGGAGTCGTACTACAGTGATCAGATACCCTTGGAGAAGGCCAACGAGAT[C/A]TTCGCCTCGTTCCGGTTTAGCTTCTCGTGCCGC<br>CTGCTCTGGAAGTGGACGGACCTGCGCCAGGTGGATGAGCTGGCCACCGAGGACAATCCAAAGATCCTGTTCCGCAGCGACTTCGTCAAGTTCAT<br>TGCGGACAAGCTGGGCCACAGC |
| FBgn0029693sechsim3 | X | 03.9 | X_03.9 | GAGGAAGCAGCGCAAATTGAGTGGGGCGACGAGCTTCTCCTAAAGGCGCGCTTCGGCAGATCGTTTGACTTGGAACGTCGTCGCATCTACGAGGA<br>GAAGGAGCGCACAGGCACCACTCTGCATGTGCATCCAGAGAGCGATGACGAGGAAGTAAATAAAGAGTTTTCCTCAACCCTGATCACGCC[C/T]CC<br>CCGTCCGTCGAGCTACAATTTGAACTCACTGAAGTATGTGGAGGCGTGTGGCCTGTTRAACAAGCGTAATGAACATGTCTACGGCGGCCTTGACATG<br>GGCGAATCATGCGCCGTCCACCTCACTTTGGGCTATCGGAGCACCTTGCTGCCCGGCGGTAAAGTGAGCGGCTTTGAAATCTGCCTGGCTGTCTA<br>CGATCGGGGTCTGAAAAACCTTAAACGATCCACTCTAACTTCGCCACTGTGCAGCTTTTCCTTTGCACGACGGGCCAGCGCCCGATGTAACCCCAC<br>CACCGATGATGAGGGGGATACTGAGGAGGATTTACCGCCGCCAACAGCGAGGCGCTGCGTGCGTCGTACTTTATTTACAGAGGAGAACACTCAGAA<br>AGAAGAGGATCCCGATTCCATACCCCTGGAA |
| FBgn0003302sechsim3 | X | 05.9 | X_05.9 | ATGGGCGGTCTGTATACCCACATTCGAGAGGTGCATGCCCCCAAGGAGCGACGCCATCCCTGCGACCAGCCAGGCTGCGAGAGGATCTACACC<br>AGCCGCATAGCCATGCAGAAGCACAAAAGGCTGAAGCACAGTCCGCGCGACCGCGATGCCTTGAAGAAATTCAT[C/T]TGCGAGCAGTGCGGAGC<br>CAGCTTTAACCAGTCGGCGAATCTTAAGTACCAYCTTAAGACAAAGCATCCCACGGAGGACGAGGTGGCTGCCAGGGAGGGCGGCGCCGGAGAA<br>CGCCATTWCTGYGACATTTGTCAAAAGGAGTTCCATTCCCGATACACCCTCAAGTACCACACTTTGCAGCAGCACGAGGTGCAGGAAGAGTTGCCM<br>CACGAATGCCAGGTCTGTGGCCGCCGAATGGCCAAGAAGTTCATGCTGCTGCAGCACATGCTGATGCACTCGAGCGACAAGTTGCCCTGCGAGC<br>ACTGCGGCAGAAGATTTGCCAGGCGATTCGAGCTCGAAGCCCACGTGAGAGCGGTCCACT |
| FBgn0029928sechsim1 | X | 06.9 | X_06.9 | ATGGAACGCTGGCAGAATCGCGTTGCCGTCGTCACTGGCGCCAGTTCGGGCATCGGATCGGCCATCGCCAAGGATCTGGTCCTCGCCGGAATG<br>ACGGTGGTGGGTCTGGCCCGACGCGTCGATCGCGTCAAGGAGCTGCARAAGGAGCTGCCGGCGGAGAAGCGCGGCAAACTCTTTGCCCTCTA<br>CTGCGACGTGGGCAACGAGAGTTCCGTGAACGA[A/G]GCCTTCGATTGGATCATCCAAAAGTTGGGCGCCATCGATGTRCTGGTCAACAAT |

| | | | | |
|---|---|---|---|---|
| FBgn0026268sechsim2 | X | 11 | X_11 | TTGCCGGCACACACACATCCCTGGTGCGCGCCTCCCTGGCGAACAGCAATGGTGATAAGAGCGCCAATGGCACCATCGCCAGCCATCTGCAATTGATACCGAAGAAGGCGCCCAGCATACCCAAACCCAAGTGGCACGCACCCTGGAAGCTGTCGCGCGTCATCTCCGGCCACTTGGGCTGGGTGCGCTGCATCGCCGTGGAGCCGGGCAACGAGTGGTTCGCCACCGGCGCTGGTGATCGTGTGATCAAGATCTGGGACCTGGCCAGCGGCAAACTGAAGCTCTCGCTCACCGGTCATGTGAGCACWGTGCGCGGTGTGGCCGTCAGCACGAAGCATCCGTATCTGTTTAGCTGCGGCGAGGATCGGCAGGTCAAGTGCTGGGATCTGGAGTACAACAAGGTGATACGGCACTATCACGGTCACTTATCGGCTGTTTACTCGCTGGCCCTGCATCCCACCATCGATGTGCTGGCCACSTCCGGTCGCGACTCCACTGCCCGCATCTGGGATATGCGAACCAAGGCGAATGTGCACACACTGACGGGGCACACGAACACGGTGGC[C/T]AGTGTGGTGGCGCAGGCCACCAATCCGCAG |
| Fbgn0030365sechsim4 | X | 11.9 | X_11.9 | AGCAGCCTTCGCGGCACTGGAGCCGGCGGTGGCGGTGTTGGATCCGGCGGAGGCAGCAAGAGGTCAAATGAATCCCGCAGTTTGGGTGGCGGCGGTAGCAAAAGTGATTTTGCGACRGCCAAAAAGAAAAAGAAGCTGGCCGAAAAGATACTGCCGCAGAAGGTGCGGGATCTGGTGCCCGAATCGCAGGCGTACATGGATCTG[C/T]TGACGTTTGAGCGGAAACTGGATGCCACCATCATGCGCAAACGTCTGGACATCCAGGAGGCCCTCAAGCGCCCCATGAAGCAGAAGCGCAAGCTGCGC |
| FBgn0025462sechsim2 | X | 13 | X_13 | CAAGATGATCTGTGCAGCCAGGGATGCGAGAACACATCGGGCGGATATCGATGCGTCTGTGACGCGGGCTATCTGCTGGACAAGGATAATCGCACGTGTCGCGCCGTCGTTCACGGCTCTAAGGAACAGCAGCCCCTTCTGCTGTACACCACCCAGATGACGATTATGGGAATGCATCTGCGCGAGGACAACGTACGAAATCATGTCTACCAGGTGGCCGGGAATCTGAGCAAGGTCATTGGCGTGGCCTACGATGGCAGCCACATCTATTGGACCAACATCCAGAACGAAGCGGAGAGCATTGTGAAGGCAAACGGAGA[C/T]GGTTCGCATGCGGAGATTCTGCTGACCTCCGGCCTGGATGCACCAGAGGACCTGGCTGTCGATTGGCTGACGCAGAACATCTACTTCTCGGACAACATAATGCGGCACATAGCCGTGTGCTCGAACGATGGTCTAAACTGTGCCGTTCTGGTCACCCAGGATGTCCACCAGCCGCGATCRCTGGCTGTTTGGCCCCAGAAGGGCTTGATGTTCTGGACGGATTGGGGTGAGAAGCCCATGATAGGGCGCGCATCCATGGACGGCAGTCGGTCGCGACCCATTGTCAGCGACAACATCGAATGGCC |
| FBgn0004649sechsim1 | X | 14 | X_14 | ACAGCGACGAAATCGACATGACCCAGTTGATGCAATTCCGTAACCAACTGCARTCGGTGGCCAAAGAGAATGGCSTGCCCAAGCTCACGTTYATGCCGTTCTGCATCAAGGCTGCCAGCATTGCGTTGTCCAAATATCCAATTGTTAACAGTTCCCTGGATTTGGCCAGCGAATCGCTGGTCTTCAAGGGTGCCCACAACATAAGCGTTGCCATCGATACACCACAGGG[T/A]CTMGTGGTGCCGAATATTAAGAACTGCCAGACAAAGACTATTATCGAGATTGCCAAGGATCTGAATGCACTGGTGGAGCGCGGTAGGACT |
| FBgn0030612dsimech1 | X | 14.9 | X_14.9 | CTCAACTGGGGTCGCAAGGGCTCGATCTGGCCACTGACTTTCGGTTTGGCCTGCTGTGCCGTCGAAATGATGCACATCGCTGCTCCSCGATACGACATGGATCGATATGGTGTCGTGTTCCGTGCATCTCC[A/G]CGTCAGGCCGATGTCATCATCGTCGCTGGCACGYTGACCAACAAAATGGCACCGGCCCTGCGAAAGGTCTACGACCAAATGCCCGAGCCACGTTGGGTCATCTCCATGGGCAGCTGTGCCAACGGCGGTGGCTACTACCATTACTCGTACTCCGTCGTGCGTGGCTGCGATAGGATAATCCCCGTCGACATATACGTACCCGGTTG |
| FBgn0030718sechsim1 | X | 15.8 | X_15.8 | GCGCCTGTTTCTTGACAAGGCCGGTGGCCAGTTATCGCCACAGTGGGAGGAGGACGACGATCCCAGTCATCAGCTAAAGGAACAGCTGCTACATCAACAGCAGCCCCATGYTGCTAATGGAGGCTCCTCCTCCGGTAACCATCAGCCGTTTGCCCGATCCGGWCGSCTGTCGATGCAGTTTCTGGGCGATGGCAATGGCTACAAGTGGATGGAGGC[A/G]GCCGAGAAACTACAGAATCAGCCGCCAGCCCAGCAGAYATATCAGCAGGGTGGTCATCATGCTGGTCACGGTC |
| FBgn0000163sechsim4 | X | 17 | X_17 | ACCACATACAGCCAGGAGCTGGAGCAGAAGGTGGGCGAGATGAGTCGCCAGTTGACGGAACTGCTGCAGGAACGCGACAGGGAGAGGGACAAGGATAAGCCGCGCTTCGCAGCCCGCAGGCGGCGTKGCTGCGCCTGGCAAGCGGACTGCTCACGGCAGTCCCAATGTCCGACCCTCGCGCCTGGTGGCCTGGATGAACCTGTCCAAYTGGTTCAAGAGCAGACCCAGCGTGGAAACGCTGCGGGAACAGCGCATCTTCTTCGACGA[A/G]CCCTGCTTCGATACGGACCTGGAGATGGTCCTCAAGCACGACCAGCACCGCACTGTGCCGCGCATCGTTGTGGACTGCTGCGATCTTATCGAGCAAAAGTACCGCAGATCCACGCAGCCCATCGAGGGCATCTATCGGCAGTGCGG |
| FBgn0000084sechsim2 | X | 20 | X_20 | GTGCCCACGGTTAAGGACGCAGCTCCCTTCGACGCCGCCCAGGACGCCCAGGTGCTGCGGGCGGCGATGAAGGGATTCGGCACCGACGAGCARGAGATCATCGACGTGCTCGTCGGCAGGAGCAACCAGCAGAGGCAGACGATCAAGGCGGTTTACGAGGC[A/G]GAGTTCGAGCGCGACCTGGTSGACGATCTAAAGGACGAGCTGGGCGGCAAGTTCGAGGACGTGATCGTGGGTCTAATGATGCCRCCGGTGGAGTACCTGTGCAAGCAACTGCACGCAGCCATGGCGGGCATCGGCACCGAGGAGGCCACACTCGTCGAGATCCTGTGCACCAAGACCAACGAGGAGATGGCCCAGATCGTGGCCGTCTACGAGGAGCGCTACCAGCGTCCGCTGGCCGAGCAGATGTGCAGCGAGACCTCCGGCTTCTTCCGCCGCCTGCTCACGCTGATCGTGACCGGAGTGCGTGACGGACTGGACACGCCCGTCGACGTCGACCAGGCCAAAGAGCAAGCCGCCCAGCTCTACTCGGCCGGCGAGGCCAAGCTGGGAACGGACGAGGAGGTMTTCAACCGGATCATGTCGCACGCCAGCTTCCCACAGCTGCGACTCGTCTTCGAGGAGTACAAGGAGCTCTCCGGGCAGACCATCGAGCAGGCCATCAAGCACGAGATGTCCGACGAGCTGCACGAGGCCATGA |

| | | | | |
|---|---|---|---|---|
| FBgn0003950sechsim3 | X | 20.9 | X_20.9 | ACCCTGTGCAAGCGTGTGGAGCGGGACTCGCAATCGTCGGCGGAGGCGCGCGACCTCGTGMAGCAGATCTTCCGGCTCCTGGAGCACTACGA<br>GACACACCATCAGCAGCTGGCGGAGAAGATCCAGCAGACGGAGCTTGCGGCGGAGCAGGCGGCTCGCATCATGCCCATGTGGGCGGATGAAA<br>CTGGTGTTGGCTCCGCTTCTGGCAACTGGAACACGGCGATGTCTACCACGCTCATCTCCAGTGAGATTTGCACTCCGCCTCTGGCTCAGCCTGCC<br>GCTGATGAAGCGCTTCTCGGTGAAATCACTGGCAAGGATGTTGTTAAGYCCCATGTGGAACAGCACACGTATCCTGACCTGGGATACCCCTATCAATT<br>GTTG[A/G]CATTGGCTCCGCCAGGTGGAGTTCCAGTTGCAGGCACCTTTGTGCCTGTGACTCGCCTTCCTGCCCACTTGCTTGGGGCTTCTGAACA<br>CCCTGTTCCTCTGGTCCCAGTTGCAAGGGTCTACAGTGATCCTACGAAAACGTCGAGACAACCAGTTTTTCGGAGTAGCGCTCCTGC |
| FBgn0000719dsimech1 | X | 22.2 | X_22.2 | ATATTATCAAGTGAGGAAGACGAGTATACCACAGAGACGGCAACAACCACAGGGGACTCAACGGTTGCGGAGGCGTCAATGGATACATCGACTGCC<br>ACCTYCACCTCTGGCCAATCATCACCACATCCGCCAGAAGAACCCGAGATAGATGAACGTGAAAATCGCCTGGTCTTGATAAAATCGAAAGTGCAAC<br>CGGTTCAGTTAACGACAACAGCTTCGGCAACTGCAGCTGATGTTGCTGATTCAAGCTCCTCAACTGACCGGTTTCACTATCAGCATTTTGTCGAAGATG<br>AGGCCGCTAGCAGTACCACGACGGCAACACCAGAGCCCAGCA[A/G]CAGCACTCCCGGCGATCCGATCGAGCAGTCCGMTATGCCGGYGAGT<br>GACAATGACAATTTAATGACAAATACAATTGGCGGACGCGGYGACGACGATAATGATRGGGGCCATAAGGCTACCAGTGAAATTGATGTCCAACAGGAA<br>CTGCGGCTCATCAACGAGCTGGTAAAGGGCAAGCAAGGAAAGCAGCATCAGCCACAGAAACAAGAGCTTGAGCCAACCAGTACAGAAATAACTTCA<br>GCGTCAACATCAACAACAACTGAGGATGCCACCACCACAGCAGC |
| FBgn0000719dsimech4 | X | 22.2 | X_22.2 | ATATTATCAAGTGAGGAAGACGAGTATACCACAGAGACGGCAACAACCACAGGGGACTCAACGGTTGCGGAGGCGTCAATGGATACATCGACTGCC<br>ACCTYCACCTCTGGCCAATCATCACCACATCCGCCAGAAGAACCCGAGATAGATGAACGTGAAAATCGCCTGGTCTTGATAAAATCGAAAGTGCAAC<br>CGGTTCAGTTAACGACAACAGCTTCGGCAACTGCAGCTGATGTTGCTGATTCAAGCTCCTCAACTGACCGGTTTCACTATCAGCATTTTGTCGAAGATG<br>AGGCCGCTAGCAGTACCACGACGGCAACACCAGAGCCCAGCARCAGCACTCCCGGCGATCCGATCGAGCAGTCCGMTATGCCGG[C/T]GAGT<br>GACAATGACAATTTAATGACAAATACAATTGGCGGACGCGGYGACGACGATAATGATRGGGGCCATAAGGCTACCAGTGAAATTGATGTCCAACAGGAA<br>CTGCGGCTCATCAACGAGCTGGTAAAGGGCAAGCAAGGAAAGCAGCATCAGCCACAGAAACAAGAGCTTGAGCCAACCAGTACAGAAATAACTTCA<br>GCGTCAACATCAACAACAACTGAGGATGCCACCACCACAGCAGC |
